# DNA double-strand break end synapsis by DNA loop extrusion

**DOI:** 10.1101/2021.10.20.465154

**Authors:** Jin Yang, Hugo B. Brandão, Anders S. Hansen

**Affiliations:** Department of Biological Engineering, Massachusetts Institute of Technology, Cambridge, 02139; The Broad Institute of MIT and Harvard, Cambridge, 02142

## Abstract

DNA double-strand breaks (DSBs) occur every cell cycle and must be efficiently repaired. Non-homologous end joining (NHEJ) is the dominant pathway for DSB repair in G1-phase. The first step of NHEJ is to bring the two DSB ends back into proximity (synapsis). However, although synapsis is generally assumed to occur through passive diffusion, we show here that passive diffusion is unlikely to be consistent with the speed and efficiency of NHEJ observed in cells. Instead, we hypothesize that DNA loop extrusion facilitates synapsis. By combining experimentally constrained simulations and theory, we show that the simplest loop extrusion model only modestly facilitates synapsis. Instead, a loop extrusion model with targeted loading of loop extruding factors (LEFs), a small portion of long-lived LEFs as well as LEF stabilization by boundary elements and DSB ends achieves fast synapsis with near 100% efficiency. We propose that loop extrusion plays an underappreciated role in DSB repair.

## Introduction

DNA double-strand breaks (DSBs) can be caused by environmental agents such as radiation and drugs [1–3], and endogenous metabolism such as transcription and replication stress [4, 5]. For example, normal metabolism has been estimated to cause ∼1-50 DSBs per human cell per day [6, 7]. Consequently, fast and reliable DNA repair is necessary to prevent deleterious chromosomal rearrangements such as translocations, inversions, amplifications, and deletions [8]. The three major DSB repair pathways are non-homologous end-joining (NHEJ), alternative end-joining (Alt-EJ), and homologous recombination (HR) [9–11]. The specific DSB pathways choice depends on sequence, chromatin context, cell cycle phase, and the complexity of DSB ends [9, 11, 12]. Here we focus on NHEJ which is operational throughout the cell cycle and the dominant DSB repair pathway in G1-phase [13].

While much is known about the proteins that are recruited to DSB ends, their order of recruitment, and the molecular mechanisms involved in the repair [3, 14, 15], what all recruitment mechanisms have in common is that they are ***reactive***: recruitment of DSB repair factors begins after the DSB has occurred and been sensed. This introduces a time delay [16] during which the DSB ends can diffuse apart [17], which may delay repair, prevent repair, or result in aberrant ligation between distinct chromosomes causing translocations [18]. Indeed, prior experimental work has demonstrated that, in human cells, DSB ends can move several hundreds of nanometers apart within minutes after a DSB has occurred [19–22]. The DNA DSB repair process through NHEJ therefore requires two major steps: 1) bringing the DSB ends back into proximity (this process is called synapsis [23]) and 2) recruiting the necessary proteins to covalently ligate the synapsed DSB ends back together (**Fig. 1A**). While much is known about the second NHEJ step, the alignment and covalent linkage of synapsed broken DNA ends [11, 24], comparatively less is known about the first step, the process leading to DSB end synapsis.

**Figure 1.**
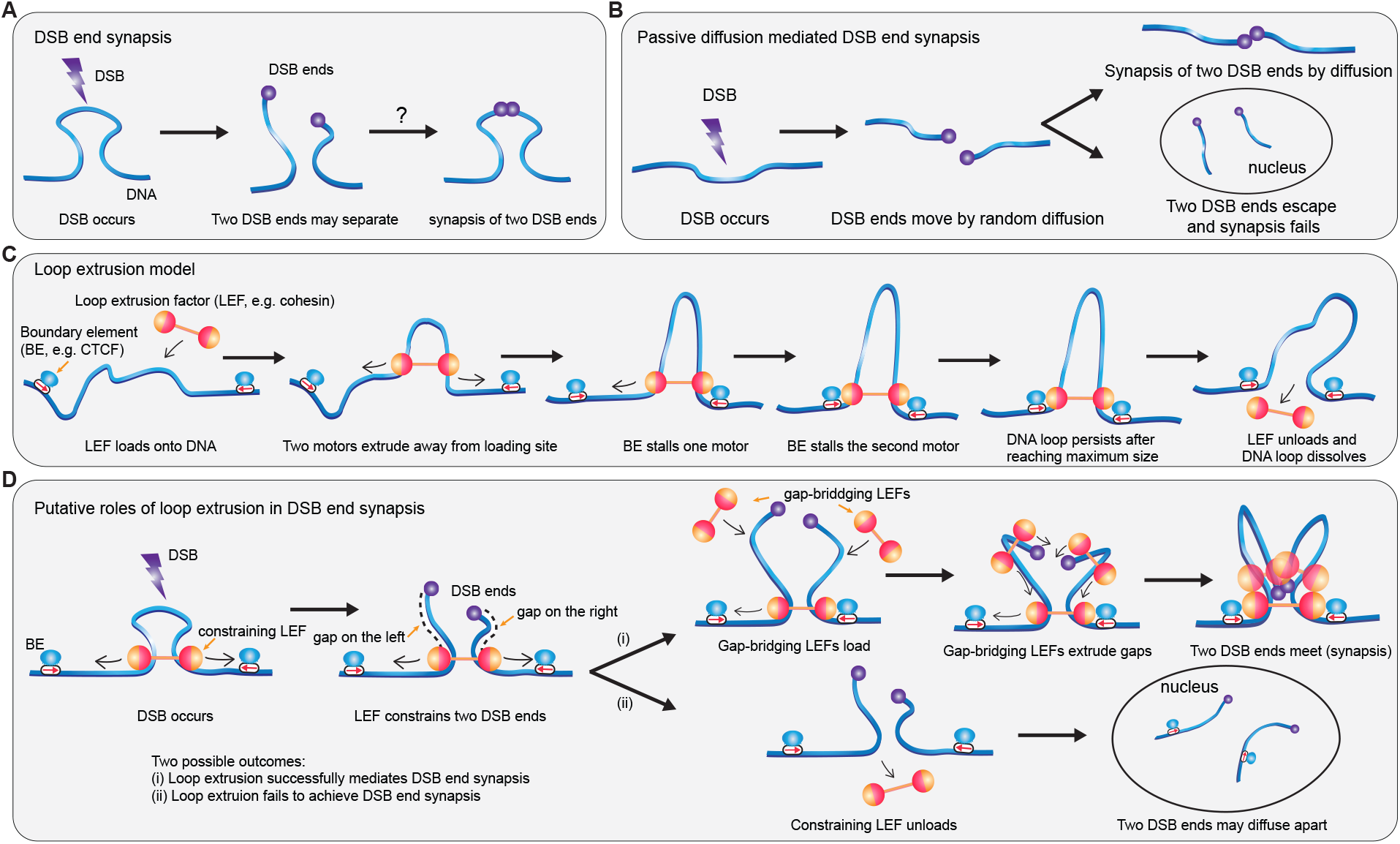
A model of DSB synapsis, mediated by DNA loop extrusion. (**A**) After the DSB has occurred, the two DSB ends may separate. How the two DSB ends are constrained from diffusing too far apart and brought back into proximity for downstream repair is not well understood. (**B**) Overview of DSB end synapsis mediated by passive diffusion. (**C**) Overview of the loop extrusion model. Loop extruding factors (LEFs) extrude bidirectionally away from the loading site; the two motors of a LEF extrude independently: after one motor is stalled by a boundary element (BE), the other motor can continue extruding until encountering a BE on the other side. (**D**) Loop extrusion may facilitate DSB end synapsis in two ways: (1) the constraining LEF prevents the two DSB ends from diffusing apart after DSB; (2) Additional gap-bridging LEFs loaded within the loop extruded by constraining LEF can extrude sub-loops to bring the two DSB ends into proximity. However, if the constraining LEF falls off before the two DSB ends are brought into proximity by gap bridging LEFs, the two DSB ends may diffuse apart. In our simulations, we assume LEFs cannot pass one another or DSB ends, and synapsis always fails once no constraining LEF remains for a given DSB.

Synapsis, the bringing of DSB ends back together for NHEJ, is generally assumed to be mediated by passive 3D diffusion [22] (**Fig. 1B**). However, here we show (see **Results** below) that passive diffusion is likely both too slow and too inefficient to be consistent with the speed and *>*95% efficiency of DSB repair by NHEJ in G1-phase in mammalian cells [25]. The inability of passive diffusion to explain the efficiency of synapsis *in vivo* suggests that alternative mechanisms must be operational inside the cell. Here, we hypothesize that DNA loop extrusion contributes to DSB repair by mediating fast and efficient DSB end synapsis. Since Alt-EJ also requires synapsis before the ligation of two DSB ends, the mechanism of synapsis facilitated by DNA loop extrusion we propose also applies to Alt-EJ [9, 26].

Loop extrusion is emerging as a universal mechanism that folds genomes into loops and domains [27]. In mammalian interphase, the primary loop extruding factor (LEF) is the cohesin complex, which is thought to extrude DNA bi-directionally at a rate of ∼0.5-2 kb/s until cohesin is blocked by a Boundary Element (BE) [28–30]. The primary BE in mammalian interphase is the insulator protein, CTCF [31]. By extruding the genome until it encounters CTCF boundaries, cohesin-mediated loop extrusion folds the genome into loops and domains known as Topologically Associating Domains (TADs) [32, 33] (**Fig. 1C**). Beyond cohesin, several other Structural Maintenance of Chromosomes (SMC) family complexes function as LEFs, including the condensin complexes that mediate mitotic chromosome compaction and bacterial SMC complexes that help resolve sister chromatids [27, 34–37].

We propose that loop extrusion likely plays a role in DSB repair for at least three reasons. First, loop extrusion is operational across the entire interphase genome. Thus, the loop extrusion and DSB repair machinery will necessarily encounter each other when a DSB occurs and have to interact. Second, experimental estimates suggest that most interphase DNA is inside an extruding cohesin loop at any given time [38–40] (**Supplementary Note 3.1**). Thus, a DSB is more likely to occur inside a cohesin loop than outside. Third, unlike known ***reactive*** DSB repair mechanisms, loop extrusion could function as a ***preemptive*** mechanism that prevents the DSB ends from diffusing apart and simultaneously accelerate the synapsis process, thereby promoting fast and efficient DSB repair. To test this hypothesis, we combine analytical theory and simulations, to quantitatively investigate the extent to which loop extrusion may help DNA repair by facilitating synapsis for NHEJ. We find that DNA loop extrusion can promote very fast (∼ 10 minutes) and efficient (≥ 95%) synapsis, and identify the parameter regimes required for efficient synapsis. Finally, we make several experimentally testable predictions to probe the relationship between the role of loop extrusion and DSB synapsis in NHEJ.

## Results

### 3D diffusion may be too inefficient to promote DSB end synapsis in mammalian cells

We began by investigating the prevailing model: that passive 3D diffusion mediates DSB synapsis [22]. The average synapsis time is estimated to be around 6-11 minutes in mammalian cells based on prior experimental data [41] (**Supplementary Note 1**). To see if 3D diffusion mediated synapsis is consistent with these values, we used an analytical formula [42] to estimate the mean synapsis time in a mammalian nucleus. Using experimental estimates for the chromatin diffusion coefficient *D* = (4 ± 1.9) × 10^*−*3^ μm^2^/s [43], a confinement radius of broken DNA of 990 ± 90 nm [44] and several other parameters (**Supplementary Note 1**), we calculated that the mean synapsis time for broken DNA ends via 3D diffusion is in the range of 20-90 minutes. We note that since the experimental parameter values of chromatin dynamics and diffusion before, during, and after a DSB has occurred are associated with significant uncertainty, our estimate is too. Nevertheless, for this reason and for additional reasons we discuss in **Supplementary Note 1**, our estimates strongly suggests that 3D diffusion of DSB ends alone will fail more frequently than expected experimentally (i.e. between 20% to 70% of the time).

### A plausible paradigm for DSB repair via DNA loop extrusion

Since 3D diffusion likely results in unphysiologically slow rates of DSB end synapsis, we reasoned that alternative mechanisms facilitate the synapsis process in parallel. One mechanism that is operational in parallel to 3D diffusion, and could facilitate DSB end synapsis is DNA loop extrusion by loop-extruding factors (LEFs) (such as cohesins, condensins, and other SMC complexes that may be operational inside the nucleus [37]). We hypothesize that loop extrusion facilitates DSB synapsis and repair in two ways.

First, since most of the genome is inside LEF loops [38–40], a DSB is statistically more likely to occur inside a loop than outside. If DSBs occur inside a LEF-mediated DNA loop, the DSB ends are constrained and unable to diffuse too far apart (**Fig. 1D**), and we call such a LEF a **constraining LEF**. The constraining LEFs’ presence on DNA provides a time window of opportunity for the two DSB ends to synapse either through passive diffusion [17] or through the action of **gap-bridging LEFs** (explained below), where we define a **gap** as the DNA segment between the constraining LEF and the broken DNA end (**Fig. 1D**).

Second, while the constraining LEF holds together the two pieces of DNA, **gap-bridging LEFs**, dynamically loaded between one DSB end and the constraining LEF can extrude loops that bring each DSB end into proximity with the constraining LEF (illustrated in **Fig. 1D**). If both sides of the DSB are extruded by gap-bridging LEFs, the DSB ends will be brought into spatial proximity, thereby achieving synapsis (**Fig. 1D**, top branch); the stochastic nature of LEFs binding to and unbinding from gaps means that multiple attempts may be required to simultaneously bridge both gaps via gap-bridging LEFs before synapsis is achieved (success) or until the constraining LEF unloads (failure) (**Supplementary Fig. 1**).

In the above, we assumed that LEFs stop extruding when they reach a DSB end (i.e., they do not “fall off” the DNA at the site of DSB) [45]. In addition, we note that any LEF can serve as “constraining” or “gap-bridging”, and that designation only depends on its current position with respect to the DSB. For example, a constraining LEF unloaded from one locus can reload at another DSB site to function as a gap-bridging LEF. Altogether, the action of constraining LEFs and gap-bridging LEFs, may constitute a new paradigm for thinking about the process of DSB end synapsis.

Thus, we set out to test our hypothesis that loop extrusion may facilitate DSB repair by facilitating synapsis by asking two questions: 1) Can the process of DNA loop extrusion achieve more efficient synapsis than 3D diffusion? 2) If so, can we identify physiologically plausible conditions for loop extrusion (e.g. LEF density on DNA or LEF processivity) that achieve near-perfect synapsis efficiency?

### A simple 3-parameter loop extrusion model predicts modest synapsis efficiency

To test our hypothesis that loop extrusion may promote DSB repair by facilitating DSB synapsis, we performed simulations of loop extrusion dynamics that incorporate DSBs, LEFs, and BEs. We simulated chromosomal DNA as a 1D lattice of sites that could be bound by LEFs. BEs were placed periodically on the lattice to create TADs of various sizes within the experimentally expected range [46–48] (**Fig. 2A**). In our simulated chromosomal DNA, we introduced DSBs approximately every 10 Mb, and followed the LEF dynamics on this lattice. LEFs were allowed to dynamically load to any lattice site not occupied by other LEFs, extrude loops, and dissociate. LEFs were not allowed to extrude past each other or DSB ends. We recorded the percentage of the DSB ends synapsed through extrusion (i.e., the synapsis efficiency) and the mean synapsis time (**Fig. 2A**). Example videos of successful and failed synapsis events can be found in **Supplementary Videos 1-2**. In its simplest form, 3 parameters are necessary to simulate loop extrusion (**Fig. 2B**): 1) the LEF separation, i.e., the total DNA length divided by the total number of LEFs on DNA; 2) the LEF processivity, i.e., the average length of DNA extruded by an unobstructed LEF (processivity = LEF residence time × extrusion speed); 3) the boundary strength, defined as the probability for a BE to stall LEF extrusion when the LEF encounters a BE, which can be conceptualized as the occupancy of CTCF binding sites. We then simulated loop extrusion and DSB synapsis across the whole range of boundary strengths (0 to 1) and a plausible range of processivities and separations, constrained by recent experimental measurements and models [32, 39, 40] (**Supplementary Note 3.1**).

**Figure 2.**
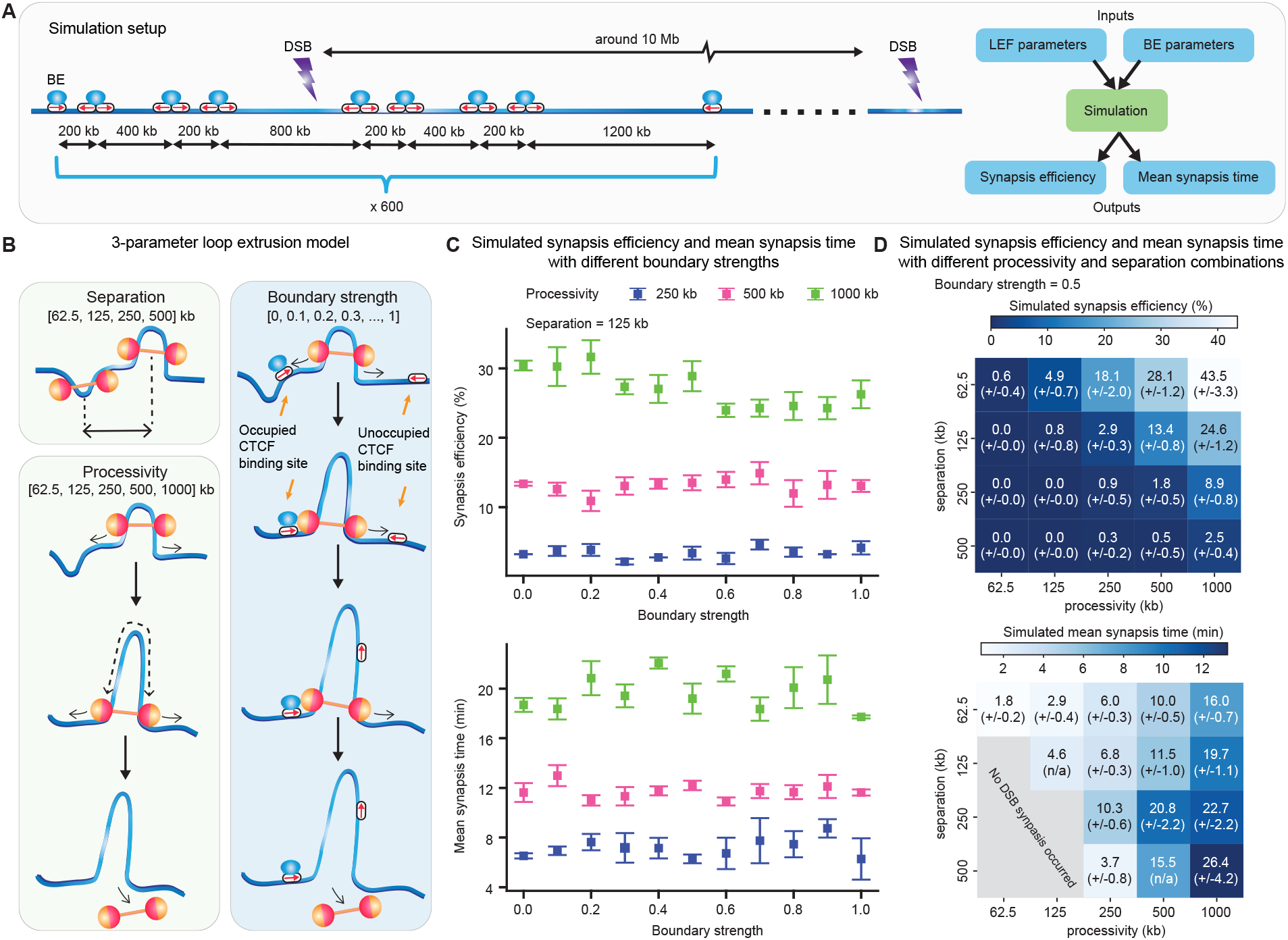
High LEF processivity and low LEF separation increase synapsis efficiency. (**A**) Overview of simulation setup. (**B**) 3-parameter model of loop extrusion-facilitated DSB end synapsis. LEF parameters (left panel): processivity and separation; BE parameter (right panel): boundary strength. (**C**) Boundary strength has little impact on DSB end synapsis under the 3-parameter model. The error bars represent the standard error of mean, *n* = 3 independent simulations. (**D**) Low synapsis efficiency under the 3-parameter model. Heatmaps of synapsis efficiency (top) and mean synapsis time (bottom, assuming total extrusion speed of 1 kb/s) for different LEF processivity and separation combinations (numbers in brackets show standard error of mean, *n* = 3 independent simulations, with 216-218 DSB events per simulation). The grey area in the bottom plot indicates no DSB synapsis occurred.

First, we studied the role of boundary strength (i.e. the probability that a LEF will stall at a BE) on synapsis efficiency. On one hand, we reasoned that boundary strength may help increase synapsis efficiency by shortening the gap between the constraining LEF and DSB ends, thus accelerating the gap-bridging process. On the other hand, the boundary strength could decrease synapsis efficiency by reducing the probability of DSBs occurring inside a loop since higher boundary strength will reduce the overall loop size, i.e., a smaller fraction of the chromosome will be extruded into loops. Surprisingly, our simulations revealed that both synapsis efficiency and mean synapsis times were largely independent of boundary strength (**Fig. 2C**). On investigating why boundary strength played such a small role on synapsis efficiency and mean synapsis times, we found that LEFs are more likely to be stalled by other LEFs before encountering BEs for the extrusion parameter values we investigated. Thus, BEs have a relatively small effect on LEF loop sizes (**Supplementary Fig. 2**). We conclude that boundary strength only mildly affects synapsis efficiency within the simple loop extrusion model and therefore we henceforth use a boundary strength of 0.5 (which is also in line with experimental estimates of CTCF binding site occupancy [39, 40]).

In contrast to boundary strength, however, simulations showed that LEF processivity and separation both strongly affect DSB synapsis efficiency and speed (**Fig. 2D**). Interestingly, when synapsis is achieved, the mean synapsis time is reasonably close to the physiological timescales of 6-11 minutes (**Fig. 2D**, bottom panel) [41]. Generally, we found that low separations and high processivities yield higher synapsis efficiency (**Fig. 2D**, top panel). The intuition is that low separations between LEFs means that there are more LEFs on the DNA to both help constrain the DSBs as well as bridge the gaps between the constraining LEF and the DSB ends. High processivities make it so that LEFs remain longer on the DNA affording a greater time window for LEFs to achieve synapsis, leading to higher mean synapsis time (**Fig. 2C**, bottom panel; **Fig. 2D**, bottom panel). Nevertheless, we find that the simple loop extrusion model only achieves moderately efficient DSB end synapsis (<45% synapsis) and requires a high processivity/separation ratio (e.g., processivity = 1000 kb; separation = 62.5 kb). These processivity and separation values are likely on the border of the physiologically plausible range for loop extrusion in mammalian interphase (**Supplementary Note 3.1**).

Together, these results show that whereas BE strength minimally affects synapsis, a high processivity/separation ratio facilitates DSB end synapsis with moderate efficiency. Thus, while loop extrusion can facilitate DSB synapsis, the simple loop extrusion model is too inefficient to do so in mammalian interphase.

### Probability theory elucidates two relative timescales underlying synapsis efficiency

To mechanistically and quantitatively understand why the simple loop extrusion model only achieved moderate synapsis efficiency, we formulated a bottom-up and mechanistically precise analytical theory that predicts synapsis efficiency as a function of loop extrusion parameters (**Supplementary Note 2**). We found that synapsis efficiency, *P*_synapsis_, can be decomposed into the product of two probabilities: the probability of the DSB occurring inside a DNA loop such that the broken DNA ends remain constrained by LEFs, *P*_constrained_, and the conditional probability of end-joining given that the DSB is constrained, *P*_end-joining|constrained_ (**Fig. 3A, Supplementary Note 2.1**).

**Figure 3.**
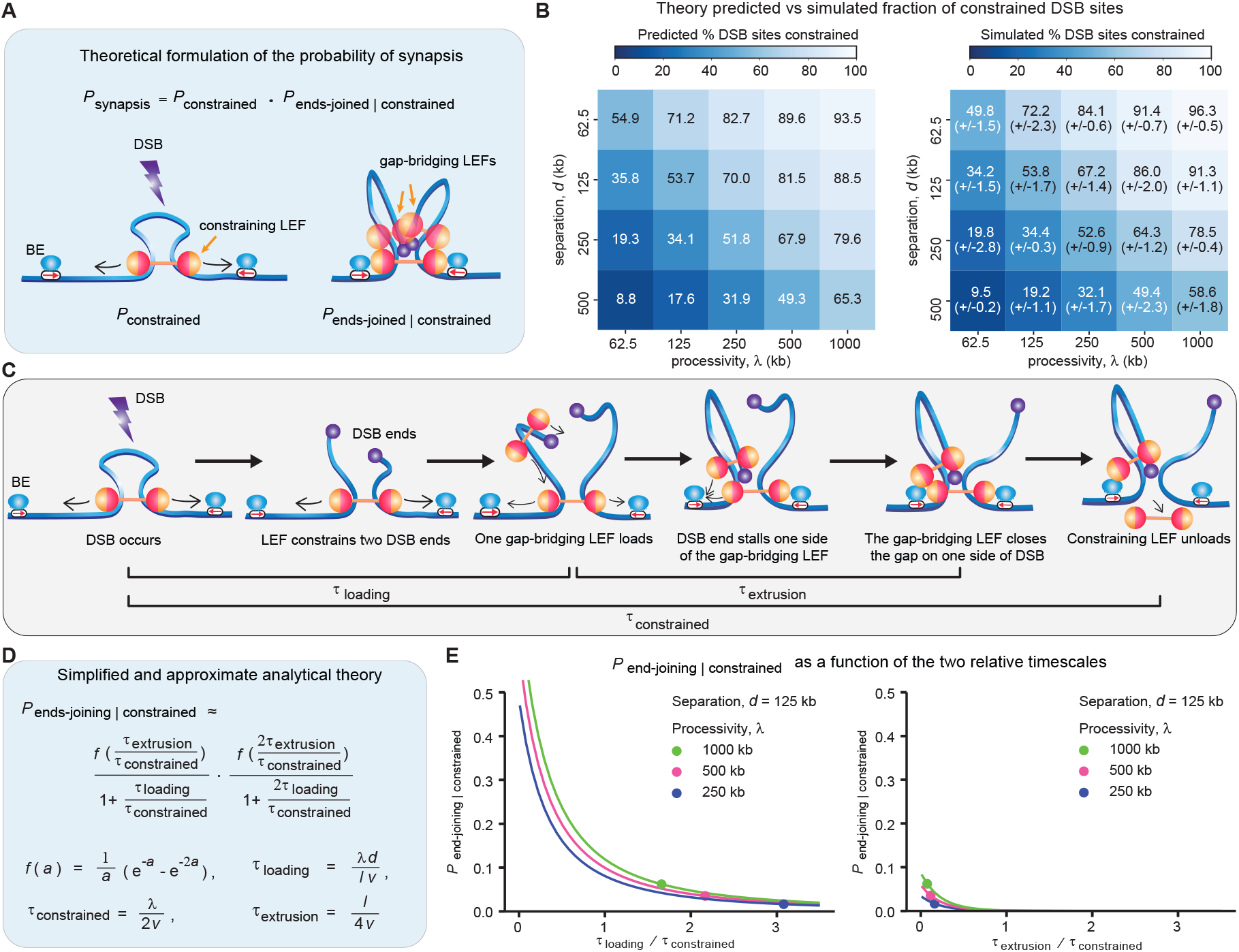
Synapsis can be quantitatively predicted and mechanistically understood using an analytical theory. (**A**) The probability of synapsis can be decomposed into the probability of being constrained and the conditional probability of gap-bridging given that the DSB was constrained. (**B**) *P*_constrained_ can be predicted by the ratio of processivity and separation. Heatmaps of predicted (left) and simulated (right; numbers in brackets show standard error of mean, *n* = 3 independent simulations, with 216-218 DSB events per simulation) fraction of constrained DSB sites with different combinations of processivity (y axis) and separation (x axis). Boundary strength = 0.5 was used in the simulations. (**C**) Three important timescales in DSB end synapsis. (**D**) *P*_end-joining | constrained_ is determined by two relative timescales: the ratio of loading time and constraining time, and the ratio of extrusion time and constraining time. *λ* is LEF processivity, *d* is LEF separation, *l* is the average LEF loop length (a function of *λ* and *d*), and *v* is the extrusion speed in one direction (i.e., 1/2 the total extrusion speed). (**E**) Larger improvement on synapsis efficiency can be achieved by reducing *τ*_loading_/*τ*_constrained_ than by reducing *τ*_extrusion_/*τ*_constrained_. The data points (circles) indicates the *P*_end-joining|constrained_ at the separation and processivity indicated in the legend, whereas the line plots show how *P*_end-joining | constrained_ varies with the ratio of loading time and constraining time (left panel) and the ratio of extrusion time and constraining time (right panel), while holding the other ratio constant at the values corresponding to the circle data points.

To build intuition, we examine **Fig. 3A**: *P*_synapsis_ depends on both *P*_constrained_ and *P*_end-joining|constrained_. On considering the temporal order of events, however, we note that end joining may only occur if the constraining LEF is present (hence the conditional probability). As such, *P*_constrained_ sets an upper bound for the *P*_synapsis_. Therefore, to achieve near-perfect synapsis efficiency, it is necessary to have *P*_constrained_ as close to 1 as possible, so we first consider *P*_constrained_.

*P*_constrained_ can be calculated as the fraction of the genome that is covered by LEF-mediated DNA loops. By accounting for the effect of BEs on LEFs, we derive an expression for *P*_constrained_ that accurately estimates the fraction of the genome inside loops (see (28) in **Supplementary Note 2.2**), as shown in **Fig. 3B**. Both our theory and simulations show that the larger the processivity, *λ*, and the smaller the separation, *d*, the higher the fraction of DSB sites constrained. This indicates that to achieve near-perfect synapsis efficiency mediated by DNA loop extrusion, it is first necessary that the genome has a high coverage by loops.

Next, to understand the role of gap-bridging LEFs on mediating synapsis, we examined how *P*_end-joining|constrained_ modulates the DSB synapsis efficiency. We derived a general analytical expression for *P*_end-joining|constrained_ that accounts for gap-bridging LEFs that load and finish extruding gaps on both sides of the DSB before the constraining LEF unloads. As simplifications, we assumed that only one gap-bridging LEF on each side of the DSB is bridging the gap, and that if one or more gap-bridging LEFs are present within the constraining LEF at the time of DSB occurrence, one of the two gaps is bridged already (Eqs.(68)-(71), see **Supplementary Note 2.3**). We found that two relative timescales dominate the simplified expression of *P*_end-joining|constrained_: the ratio of loading time *τ*_loading_ to the constraining time *τ*_constrained_, and the ratio of extrusion time *τ*_extrusion_ to the constraining time *τ*_constrained_ (**Fig. 3C-D, Supplementary Note 2.3**). *τ*_loading_ is the time it takes for gap-bridging LEFs to load into the gap between the DSB and the constraining LEF; *τ*_constrained_ is the time from when the DSB has occurred until the constraining LEF unloads; and *τ*_extrusion_ is the time for the gap-bridging LEFs to finish extruding the gap between the DSB end and the constraining LEF. We find that while reducing either relative timescale improves synapsis efficiency, larger improvement can be achieved by reducing *τ*_loading_/*τ*_constrained_ than by reducing *τ*_extrusion_/*τ*_constrained_ (**Fig. 3E**). Indeed, comparison of *τ*_loading_/*τ*_constrained_ and *τ*_extrusion_/*τ*_constrained_ across different combinations of processivity and separation show that loading of gap-bridging LEFs is generally the rate-limiting step of the synapsis process (**Supplementary Fig. 3**). Notably, biological processes that prolong the constraining LEF lifetime will contribute even more strongly to the efficiency of DSB end synapsis than processes that alter any single one of the relative time-scales (since *τ*_constrained_ is the denominator for both the *τ*_loading_ and *τ*_extrusion_ time-scales).

Our analytical theory allows us to understand both mechanistically and quantitatively how the various loop extrusion model parameters regulate synapsis efficiency, and reveal the relative impact of each parameter on synapsis efficiency. Recent experimental studies have proposed a wide range of possible mechanistic extensions to the simple loop extrusion models considered so far. We therefore next used our theory to estimate which of the most mechanistically plausible extensions would also likely affect synapsis efficiency, and we consider four extensions in the next sections.

### BE stabilization of LEFs and long-lived LEFs strongly improve synapsis efficiency

Having found that the synapsis efficiency depends most strongly on *τ*_constrained_, we first sought plausible biological mechanisms that would increase the constraining LEF lifetime.

First, we considered LEF stabilization by BEs (**Fig. 4A**, top panel; **Supplementary Video 3**). In mammalian interphase, cohesin and CTCF are the most prominent LEF and BE candidates *in vivo*, respectively. Recent work has demonstrated that CTCF may stabilize cohesin by protecting cohesin from WAPL-mediated dissociation and/or by facilitating ESCO1-mediated acetylation of cohesin, leading to an up to 20-fold increase in cohesin’s residence time [49, 50]. We carried out simulations where a LEF bound to a BE exhibits a 1-, 2-, 4-, 8-, or 16-fold increased residence time. By simulations and theory (extended to include the contribution of LEF-stabilization by BEs), we found that stabilization of LEFs at BEs strongly increases DSB synapsis efficiency (**Fig. 4A, Supplementary Note 2.4.1**). The extended theory provides intuition for the process (**Supplementary Fig. 4A, B**): first, stabilization of LEFs at BEs will improve *P*_constrained_ since the BE-bound LEFs have a higher processivity (which increases the average loop size, *l*) and results in a greater genome coverage by DNA loops (and hence the probability of a DSB occuring inside a loop); second, stabilization of LEFs at BEs increases *τ*_constrained_, and thereby prolongs the window of opportunity for gap-bridging LEFs to load and bridge the gaps, thus improving *P*_end-joining|constrained_ and synapsis outcome. However, stabilization of LEFs at BEs achieves increased synapsis efficiency at a cost of slower synapsis (**Supplementary Fig. 4F**). Thus, while stabilization of LEFs at BEs strongly improves synapsis efficiency, it was still insufficient (on its own) to achieve the *>*95% efficiency observed *in vivo* [25].

**Figure 4.**
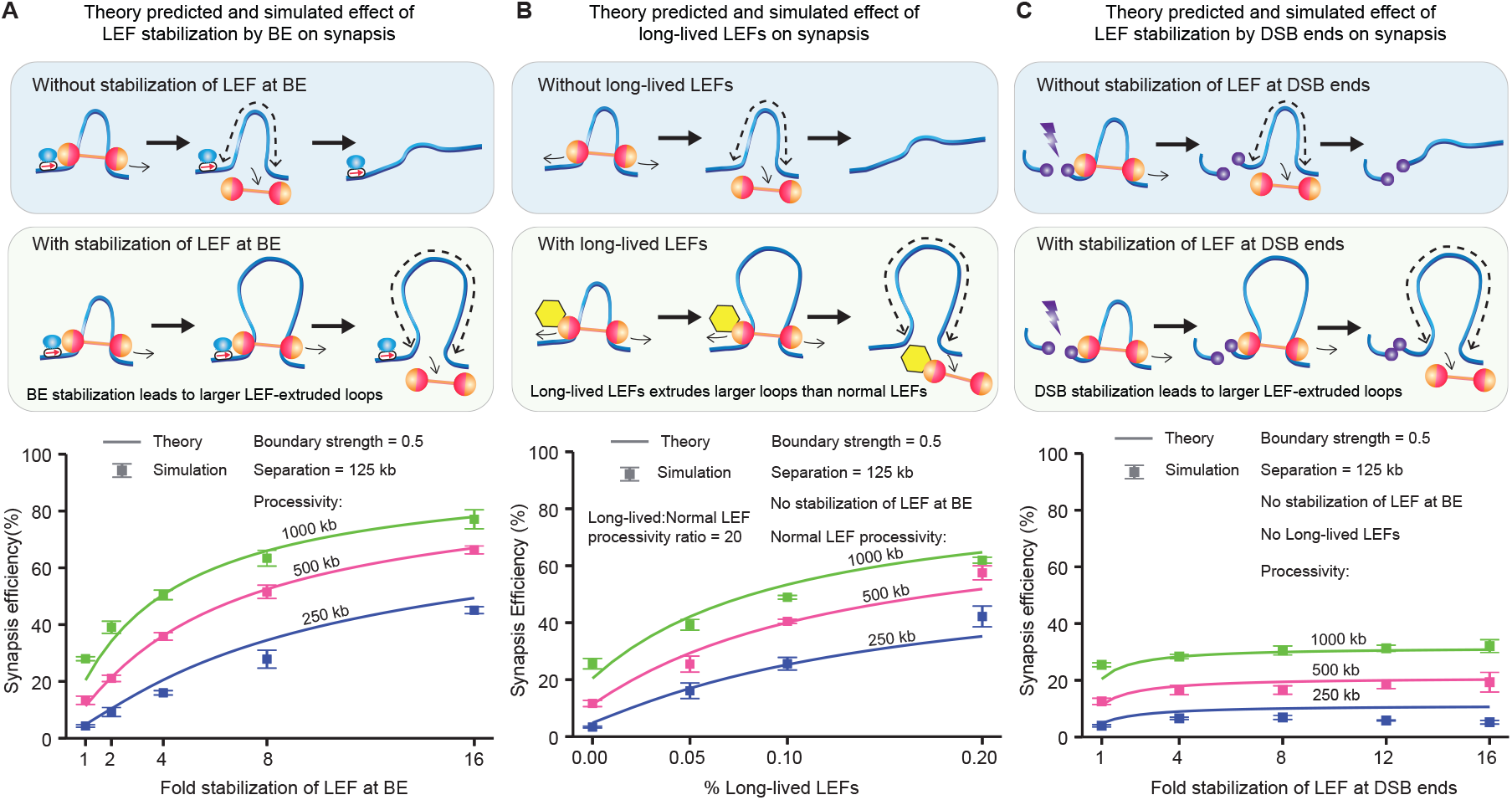
LEF stabilization by either BEs or DSBs improve synapsis efficiency, as does the presence of long-lived LEFs. (**A**-**C**) Schematic diagrams of the effects of stabilization of LEF by BE, having a small portion of long-lived LEFs, and stabilization of LEFs by DSB (top), and the corresponding synapsis efficiency (bottom) predicted by theory (lines) or obtained from 1D simulations (squares; The error bars represent the standard error of mean, n = 3 independent simulations, with 216-218 DSB events per simulation).

Second, we considered having a subpopulation of very long-lived LEFs (**Fig. 4B**, top panel; **Supplementary Video 4**). The most likely LEF candidate, cohesin, exists in multiple forms and recent work has shown that acetylated cohesin-STAG1 exhibits a much longer residence time than unacetylated cohesin-STAG1 [49]. Based on Wutz *et al*., we estimate that ∼30% of all cohesins could be acetylated, and exhibit up to 50-fold increase in residence time compared with the unacetylated cohesins (**Supplementary Note 3.3**). Thus, we carried out simulations where a sub-population of long-lived LEFs (5%, 10%, or 20%) exhibit 20-fold increase in processivity, and extended our analytical model accordingly (**Supplementary Note 2.4.2**). We found - both by simulations and our analytical model - that a small portion of long-lived LEFs improved the synapsis efficiency (**Fig. 4B**). The close agreement between our theoretical prediction and simulation results supports our mechanistic interpretations of long-lived LEFs’ role in synapsis: the long-lived LEFs facilitate synapsis mainly by acting as constraining LEFs (**Supplementary Fig. 4D,G**) and less frequently as gap-bridging LEFs (**Supplementary Fig. 4E,H**) [51]; the long-lived constraining LEFs provide a larger time window to attempt synapsis similar to stabilization of LEFs at BEs, as seen in the increased mean synapsis time (**Supplementary Fig. 4I**). Yet, once again, while long-lived LEFs strongly improve synapsis outcome, this mechanism is insufficient on its own to achieve the desired (*>* 95%) synapsis efficiency.

To understand the limitations of the two above proposed mechanisms, we looked at how they separately affect *P*_constrained_ and *P*_end-joining|constrained_. We found that stabilization of LEFs at BEs and long-lived LEFs can realize ∼100% *P*_constrained_ (**Supplementary Fig. 4J,K**), suggesting that the failure to achieve near-perfect synapsis efficiency is due to inefficient gap-bridging by these mechanisms. We thus turned our attention to additional mechanisms, which could potentially improve *P*_end-joining|constrained_, and ultimately increase *P*_synapsis_.

### DSB end stabilization of LEFs only modestly improves synapsis efficiency

To identify mechanisms that facilitate the gap-bridging process (i.e. increase *P*_end-joining|constrained_), we searched for bottlenecks within the sequence of events leading to synapsis. Aside from retention of the constraining LEF, the foremost bottleneck for efficient LEF-mediated synapsis is that it requires the simultaneous bridging of the gaps on both sides of the DSB. That is, once the DNA on one side of the gap is extruded into loops (by one or more LEFs), extrusion on the other side of the DSB needs to finish before the LEFs on the first side unload (**Supplementary Fig. 4L**). Therefore, we reasoned that factors stabilizing the association of LEFs to the DSB can help facilitate the synapsis process (**Supplementary Video 5**).

Recent experimental work [45] has suggested that cohesins associated with DSB ends are stabilized resulting in increased processivity (DSB stabilization of LEFs; **Fig. 4C**, top panel), in line with our hypothesis. We thus extended our theory to account for DSB stabilization of LEFs (**Supplementary Note 2.4.3**), and performed simulations where a LEF in contact with a DSB end experiences a 1-, 4-, 8-, 12-, or 16-fold increase in residence time. We found that while DSB stabilization does lead to improved synapsis outcome, the effect is much milder than stabilization of LEFs at BEs (**Fig. 4C**, bottom panel).

We identified two reasons for the poor performance of DSB stabilization in increasing synapsis efficiency. First, DSB stabilization does not modify *P*_constrained_ or the constraining LEF lifetime (see **Supplementary Note 2.4.3**), which limits any efficiency gains obtained by having longer-lived gap-bridging LEFs. Moreover, DSB stabilization only improves *P*_end-joining|constrained_ by reducing the pre-factor 2 in front of the two relative timescales in **Fig. 3D** (Eq.(78)) down to 1 (in the limit of infinite-fold stabilization, **Supplementary Note 2.4.3**), which has a very modest effect on *P*_end-joining|constrained_.

Together, these results suggest that *maintenance* of the end products of gap-bridging (i.e. proteins that give gap-bridging LEFs a boost in residence time) is not the bottleneck of synapsis. Instead, this suggests that *establishment* of the end products of gap-bridging are the limiting step; this led us to search for mechanisms that accelerate the rate of gap bridging.

### Targeted loading of LEFs at DSB improves synapsis efficiency by accelerating loading of gap-bridging LEFs

Guided by the objective of finding mechanisms that accelerate gap-bridging product formation, we arrived at the idea of increasing the loading rate of LEFs to the gap. This can be achieved through targeted loading of LEFs to the DSB ends (**Fig. 4C**, top panel; **Supplementary Video 6**). Experimental support for targeted-loading of LEFs at DSB ends come from recent studies that observed accumulation of SMC1 at DSBs sites leading to a ∼2-10-fold cohesin enrichment at restriction-enzyme induced DSB sites [45, 52] though two other studies reported cohesin enrichment only for S/G2 cells but not G1 cells via laser induction [53, 54], suggesting that further experimental work is needed. To test how targeted loading of LEFs to DSB ends aids synapsis, we implemented simulations where loading of LEFs within 1 kb of the DSB is 250-, 500-, 750-, 1000-, 5000- or 10000-fold more likely than at other similarly-sized genomic loci; this corresponded to about 5%, 9%, 13%, 17%, 50% or 67% of LEF loading events occurring at the DSB sites in our simulations. To test the physiological plausibility of these parameter values, we generated ChIP-seq-like data from our simulations for the accumulation of LEFs around the DSBs, and compared this to experimental ChIP-seq data [45]. We found good agreement between the ∼2-fold experimental enrichment of SCC1 in the DSB-containing TADs and our simulated enrichment of LEFs (with 250X targeted loading) (**Supplementary Fig. 5A,B**).

From our simulations, we found that targeted loading of LEFs significantly improved synapsis efficiency, and that the effect saturated quickly at ∼750-fold increased targeted loading (**Fig. 5B**), which corresponds to ∼2.5-fold enrichment of LEFs in DSB-containing TADs (**Supplementary Fig. 5A**). We additionally extended our theoretical model to include the targeted loading mechanism and found consistent results (**Fig. 5B**). Thus, accelerated loading at DSBs makes gap-bridging LEFs more likely to finish synapsis before the constraining LEF unloads (**Supplementary Note 2.4.4**); this leads to higher synapsis efficiency (**Fig. 5B**) and lower mean-synapsis time compared with the case of no targeted loading (**Fig. 5C**). Nevertheless, although targeted loading increases both the speed and efficiency of synapsis and agrees with experimental data, it still falls short of the necessesary *>* 95% synapsis efficiency seen *in vivo*.

**Figure 5.**
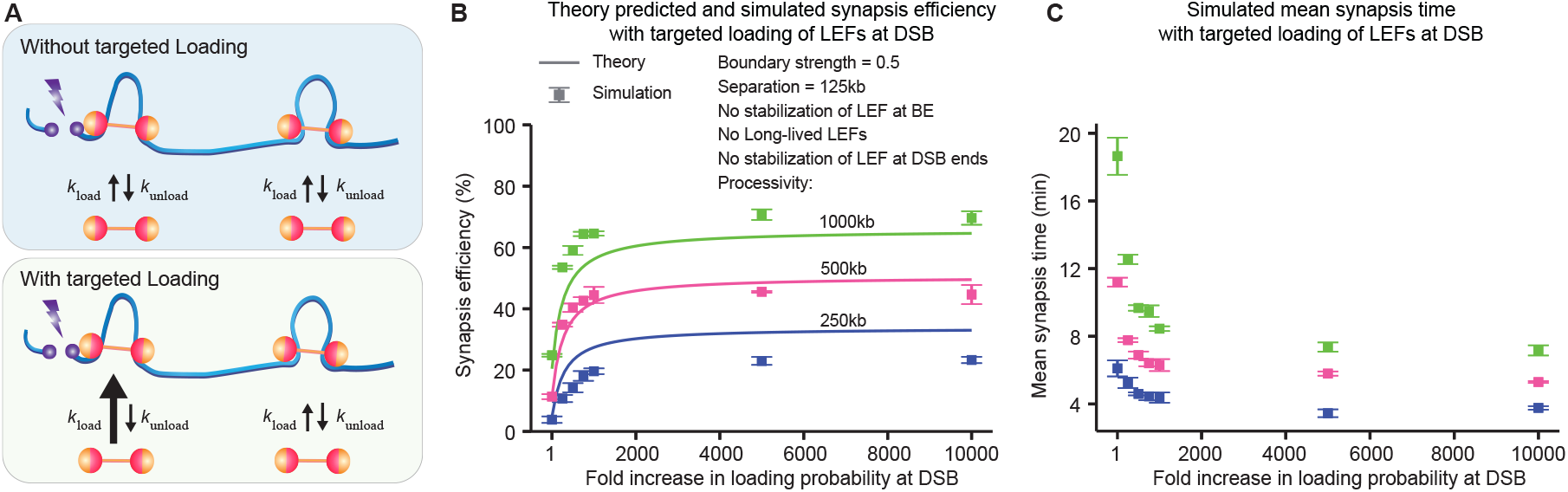
Targeted loading of LEFs at DSB accelerates synapsis and improves synapsis efficiency. (**A**) Schematic diagram of DSB synapsis without targeted loading (top panel) and with targeted loading (bottom panel). (**B**) Targeted loading of LEFs at DSB sites significantly improves DSB end synapsis efficiency. The error bars represent the standard error of mean, *n* = 3 independent simulations, with 216-218 DSB events per simulation. (**C**) Targeted loading of LEFs at DSB sites significantly reduces the mean synapsis time. Conversion of simulation time steps to synapsis time assumes total extrusion speed of 1 kb/s. The error bars represent the standard error of mean, *n* = 3 independent simulations, with 216-218 DSB events per simulation.

### Large scale simulations identify the parameter regime required for synapsis with near-perfect efficiency

Thus far, we considered four extensions of the simple loop extrusion model, and found that each individually improves synapsis outcomes, but falls short of the *>*95% synapsis likely required *in vivo*. We therefore asked whether combinations of our proposed mechanisms can achieve the necessary synapsis efficiency. To test this, we carried out a systematic large-scale sweep of all 5 model parameters which included 1) the fold stabilization of LEFs at BEs, 2) the fraction of long-lived LEFs, 3) the fold stabilization of LEFs at DSB ends, 4) the fold increase in loading rate at the DSB, 5) the ratio of LEF processivity to LEF separation. This yielded a total of 768 different parameter combinations (**Fig. 6A, Supplementary Note 3**).

**Figure 6.**
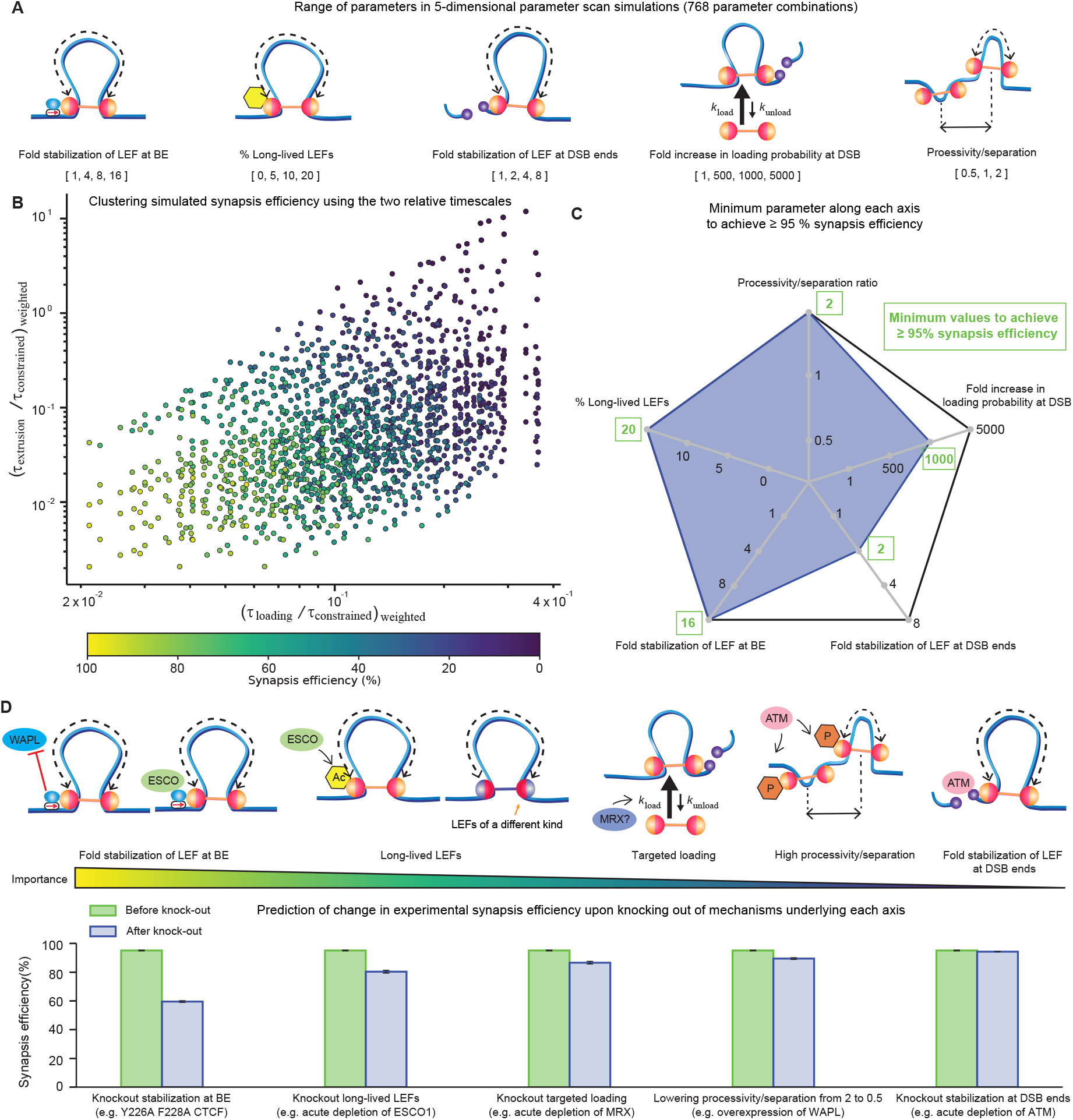
Large scale simulations reveal a physiologically plausible parameter regime that achieves synapsis with *≥* 95% efficiency. (**A**) Parameters scanned along each of the 5 dimensions. (**B**) The two relative timescales in **Fig. 3D** can cluster the 5D parameter scan data points based on synapsis efficiency. (**C**) Among all the parameter combinations that achieve a synapsis efficiency average *≥* 95% (*n* = 3 independent simulations, with 216-218 DSB events per simulation), we recorded the minimum parameter along each of the 5 dimensions. (**D**) The 5 aspects of synapsis ranked ordered based on the predicted reduction in synapsis efficiency upon knocking out the corresponding mechanism. Schematic diagram shows a plausible mechanistic basis for each of the five aspects of synapsis. The bar plot shows the average synapsis efficiency before and after knocking out each of the five mechanisms. The error bars represent the standard error of mean (*n* = 3 different parameter combinations that achieved *≥*95% synapsis efficiency).

Interestingly, the synapsis efficiency of all 768 parameter combinations separated neatly along two axes (**Fig. 6B**) composed of the two relative, weighted time-scales (*τ*_loading_/*τ*_constrained_)_weighted_ and (*τ*_extrusion_/*τ*_constrained_)_weighted_ defined by Eqs.(148)-(149) (**Supplementary Note 2.6**). This representation of our simulation results demonstrates that despite large mechanistic differences between the models, our theory identifies a universal pair of time-scales that captures the central features of the LEF-mediated DSB synapsis process.

We next focused on the models that achieve high synapsis efficiency, and sought to understand more mechanistically how our four proposed extrusion mechanisms may combine for efficient synapsis. Among the parameter combinations that met the 95% efficiency criterion, we plotted the minimum required value of each parameter (**Fig. 6C, Supplementary Video 7**). First, we found the minimal required value for fold stabilization of LEFs at BEs was 16-fold, consistent with experimental estimates of up to 20-fold stabilization [49] (see **Supplementary Note 3.2**). For the fraction of long-lived LEFs, we found 20% by our simulation sweep, and experimentally it is estimated that up to ∼30% of cohesins are acetylated (and have a longer DNA-bound residence time, see **Supplementary Note 3.3**). For the fold stabilization of LEFs at DSB ends, we found the minimal required value was 2, similar to the ∼2-4 range suggested by Arnould et al. [45] (see **Supplementary Note 3.4**). For the fold increase in loading probability at DSB, we needed a value of 1000 (compared to ∼250 suggested by our analysis of the Arnould et al. data [45], see **Supplementary Fig. 5A**). Finally, we found that we needed a processivity/separation ratio of 2 (which is within the range of ∼0.3-26 suggested by Cattoglio et al. [39] and Holzmann et al. [40], see **Supplementary Note 3.1**). We note that the three models with ≥ 95% synapsis efficiency finish synapsis within 12-14 minutes on average (assuming total extrusion speed of 1 kb/s), consistent with the 6-11 minutes synapsis time estimated from prior data [41]. While synapsis efficiency is independent of extrusion speed, synapsis time calculated from our simulations is inversely proportional to extrusion speed. For example, if we use the upper bound of experimentally observed extrusion speed of 2 kb/s instead [29], the mean synapsis time for the three models with ≥ 95% synapsis efficiency will be 6-12 minutes. Therefore, highly efficient LEF-mediated DSB end synapsis is achievable within the parameter ranges suggested by experimental data.

We next performed 1D polymer simulations to generate chromosome conformation capture (Hi-C)-like contact maps from the models which achieve high synapsis efficiency. Consistent with the experimental Hi-C maps reported by Arnould et al. [45], we find that DSBs result in an X-shaped stripe pattern in the vicinity of the break site (**Supplementary Fig. 6A**). Our simulations suggest that the stripe pattern grows quickly over time and reaches a steady state by 1 hr after DSB occurrence (**Supplementary Fig. 6A**). Quantitative differences in contact frequency between our simulations and the experiments may be due to differences in simulation implementation versus experimental conditions (see **Methods**). Moreover, these results suggest that in order to capture the temporal dynamic changes to 3D genome structure caused by DSB formation, it is necessary to perform Hi-C at shorter time-intervals post-DSB formation (e.g. at 5 min intervals), and points to the need to have fast mechanisms of inducing DSBs at specified genomic locations to better study the effect of DSB end synapsis on 3D genome organization [55].

In summary, our simulations show that with experimentally plausible parameter values, loop extrusion can achieve fast and ≥95% efficient synapsis. We thus propose that loop extrusion plays a previously unrecognized role in mediating DSB synapsis as part of NHEJ in mammalian cells.

## Discussion

Synapsis is the first step of DSB repair by NHEJ, which is the dominant repair pathway in the G1-phase. Synapsis has largely been assumed to occur by passive 3D diffusion [22]. However, we estimate that passive diffusion would lead to unphysiologically slow, or inefficient synapsis in mammalian nuclei.

Here we propose that protein-mediated DNA loop extrusion may promote fast and efficient synapsis in cells. We emphasize that loop extrusion can be a preemptive mechanism facilitating synapsis, in contrast to the reactive recruitment of DSB repair machinery after the DSB has occured where DSB ends may diffuse apart during the recruitment period. We built a probabilistic theoretical framework by which to understand LEF-mediated DSB end synapsis. The most simplistic loop extrusion model fell short of the synapsis efficiency observed *in vivo*, but did predict synapsis timescales consistent with experimental data. Guided by our analytical theory we explored four plausible extensions to the simple loop extrusion model that constituted mechanistically distinct ways to improve synapsis outcome and tested the theory with simulations of the extrusion-mediated DSB synapsis process, finding they were in good agreement. We found that while each mechanistic extension could moderately improve synapsis efficiency above the 3-parameter model baseline, it was on combining all four mechanisms that we found a regime with *>*95% synapsis efficiency. Our theory demonstrates that loop extrusion is a viable and efficient way to mediate the first step of the NHEJ process, DSB end synapsis, by co-opting the cell’s chromosome organization machinery. A broader role for loop extrusion in DNA repair is beginning to emerge. Recent experimental studies have proposed that DNA loop extrusion may facilitate DSB repair foci formation by mediating *γ*H2AX spreading [45, 55–57] and by forming structural scaffolds with 53BP1 and RIF1 to protect DSB ends from aberrant processing [58]. In addition, loop extrusion by cohesins (one of the best studied LEFs) has been proposed to facilitate V(D)J recombination [59–61] and class switch recombination (CSR) [62] by aligning the genomic loci to be recombined after activation of the DSB machinery. Our model can be readily generalized to encompass V(D)J recombination and CSR, since V(D)J recombination and CSR can be considered as special cases of NHEJ synapsis where two instead of one DSBs are induced [11]. In addition, the proposed role of LEFs in mediating *γ*H2AX spreading is synergistic with our proposed gab-bridging mechanism (**Fig. 5**). Our work therefore provides a framework to extend, integrate, and test several models.

Importantly, we note some limitations of our study. Though 3D diffusion likely occurs simultaneously and synergistically with loop extrusion, we have not considered it here. First, if two DSB ends are held together by a constraining LEF, they cannot diffuse apart and would, all other things being equal, achieve faster and higher-efficiency diffusive synapsis than without a constraining LEF. Second, with diffusion, gap-bridging LEFs may not need to achieve perfect DSB end proximity. Instead, if the gap-bridging LEFs bring the DSB ends sufficiently close, 3D diffusion may be sufficiently fast and reliable to mediate synapsis. Thus, our quantitative estimates of loop extrusion mediated synapsis should be taken as lower bounds, and we propose that loop extrusion and 3D diffusion simultaneously contribute to DSB end synapsis in cells. Finally, this study has assumed a particular set of rules by which LEFs interact with one another and other obstacles on the genome. We note, however, that as additional features of LEF-mediated extrusion come to light, such as the ability of LEFs to bypass large steric obstacles [63, 64], or each other [65, 66], one can envision extending our developed framework to account for such mechanisms.

We note several experimental directions worthy of further investigation that can help falsify or substantiate our model. Specifically, we identified five mechanistic features of loop extrusion that mediate efficient DSB synapsis: 1) stabilization of LEFs at BEs, 2) a mixture of long-lived and short-lived LEFs, 3) high LEF processivity/separation ratio, 4) targeted loading and, 5) DSB stabilization. 1)-3) all help increase the probability of a DSB occurring inside a loop (*P*_constrained_) and the constraining LEF residence time (*τ*_constrained_), whereas 4) and 5) both accelerate synapsis by increasing the chance of simultaneous gap-bridging on both sides of the DSB (**Fig. 3**). Crucially, these mechanistic extensions are all biologically plausible. First, we suggest that stabilization of LEFs at BEs could be mediated by CTCF-mediated cohesin protection from its unloader WAPL [50] or ESCO-mediated acetylation [49]. Second, we suggest that the LEF processivity/separation ratio may increase following a DSB perhaps through ATM-mediated phosphorylation of cohesin subunits SMC1 and SMC3 [67], consistent with the global stabilization of cohesins observed experimentally [45]. Third, long-lived LEFs may correspond to acetylated STAG1-cohesins [49], or LEFs of a different kind that exhibits higher processivity. Fourth, we suggest that targeted LEF loading at DSBs may be mediated by the MRX complex as knockdown of MRX led to a significant decrease in cohesin loaded at DSBs [52]. Fifth, we speculate that LEF stabilization by DSBs may be mediated by the ATM complex, which phosphorylates cohesins and accumulates cohesins in the DSB-containing TAD [45]. Finally, we note that our mechanistic understanding of loop extrusion and DSB repair is advancing rapidly, such other factors and mechanisms are likely to be found to play a role beyond the ones mentioned above.

To facilitate the experimental testing of our model, we used our theory and simulations to make specific and quantitative predictions (assuming that cohesin and CTCF play the main role of LEF and BE; **Fig. 6D, Supplementary Fig. 6B**). Our theory predicts that reducing *τ*_constrained_ would most strongly decrease synapsis efficiency. Indeed, our simulations predict loss of stabilization of LEFs at BEs to have the strongest effect: loss of stabilization of LEFs at BEs would reduce LEF-mediated synapsis efficiency to ∼60%. Experimentally, this may be tested by mutating Y226 and F228 in the N-terminus of CTCF since this is predicted to eliminate CTCF-mediated stabilization of cohesin without affecting CTCF binding to DNA [50]. Next, we predict that eliminating long-lived LEFs would have the second-strongest effect, reducing LEF-mediated synapsis efficiency to ∼80%. For example, this could be achieved through acute auxin-inducible degron (AID) depletion of ESCO1 or STAG1. Knocking out targeted loading would have the third-strongest effect, reducing LEF-mediated synapsis efficiency to ∼87% and drastically increasing the mean synapsis time by ∼181%. Experimentally, acute depletion of MRX would be one way of testing this. Lastly, although we predict lowering the processivity/separation ratio from 2 to 0.5 and knock-out of DSB stabilization to be relatively mild, reducing LEF-mediated synapsis efficiency to ∼89% and ∼94% respectively, their knock-out would slow down synapsis substantially increasing the mean synapsis time by ∼67% and ∼39%, respectively. Note that synapsis is just the first step of the NHEJ pathway, and if downstream NHEJ processes rely on a fast synapsis step, e.g. if NHEJ-related protein assembly is only stable for a limited duration, then significant slow down of synapsis due to knock-out of targeted loading and DSB stabilization could translate into even lower overall NHEJ efficiency than the predicted synapsis efficiency in **Fig. 6D**. Finally, given the high redundancy between the mechanisms considered in our study, we also predicted the quantitative effect of double knock-outs and alteration of the extrusion processivity/separation (**Supplementary Fig. 6B**).

Broken DNA-end synapsis is a key but understudied step in DSB repair. Our theory provides a new framework for rethinking this initial step of the NHEJ process in the context of our current understanding of 3D chromosome organization by loop extrusion. In summary, we predict that DNA loop extrusion plays a previously underappreciated role in DNA repair by mediating DNA double-strand break synapsis.

## Methods

### Time steps and lattice set-up

We use a fixed-time-step Monte Carlo algorithm for 1D simulations as described in previous work [68]. Each lattice site corresponds to 1 kb of DNA, and we define the chromosome as a lattice of *G* = 2164800 sites. Loop extruding factors (LEFs) are comprised of two motor subunits that move bidirectionally away from each other one lattice site at a time. Like most cohesin simulations [37], we assume LEFs cannot bypass each other upon encounter. LEFs also cannot extrude past the first and the last lattice sites so that LEFs do not “walk off” the chromosome.

### Boundary elements

Each boundary element (BE) occupies a lattice site. BEs are directional (indicated by the red arrow in schematics) and only if the LEF motor subunit’s extrusion direction is convergent with the direction of a BE, will the motor subunit be stalled by the BE with a probability equivalent to boundary strength *b*. Unless specified otherwise, a boundary strength of *b* = 0.5 is used, in line with experimental estimates of CTCF binding site occupancy [39, 40]. Once a motor subunit is stalled by a BE (i.e. the subunit stopped at the BE lattice site), no further movement of the subunit is allowed until the LEF dissociates from the locus and reloads back onto the chromosome somewhere else. Only one motor subunit can occupy a BE lattice site at a time. We place BEs on the chromosome so that TADs of the sizes shown in **Fig. 2A** are achieved.

### DSB sites

Each DSB site occupied two lattice sites on the chromosome, each of which corresponded to a DSB end. We introduced DSBs approximately every 10 Mb: we first randomly picked the DSB site in the very first TAD on the chromosome, and then we found the TAD 10 Mb to the right of the first DSB site, and randomly induced the second DSB in the TAD (so that the distance between DSBs and BEs were randomized), and so forth. This results in altogether 216-218 DSB sites on the chromosome. We first ran 100 thousand time steps of the 1D simulations, and then introduced all the DSBs simultaneously. After DSB occurrence, LEFs were not allowed to extrude past DSB ends. We allowed multiple motor subunits to occupy the DSB end lattice site at the same time, to enable simulations with targeted loading of LEFs to DSB sites.

### LEF association and dissociation rates

All 1D simulations were performed with a fixed number of LEFs, determined by the ratio of chromosome length *G* and LEF separation *d* (i.e., the inverse of LEF density). The dissociation rate was linked to the LEF processivity *λ* (i.e., the average length of DNA extruded by an unobstructed LEF before it dissociates) (**Supplementary Note 2.3.2**). After a LEF dissociates from the chromosome, it immediately and randomly reloads onto a lattice position on the chromosome that is not occupied by other LEFs’ motor subunits. The only lattice sites where co-occupancy and loading of multiple LEFs were allowed is at DSB ends.

### Monitoring of synapsis events

After DSBs occur, at every time step, for each DSB site, we first checked whether there was at least one LEF whose two motor subunits were on opposite sides of the DSB (i.e., whether the DSB site was constrained by at least one LEF). For the constrained DSB sites, we counted the number of lattice sites between the innermost constraining LEF that was not extruded into loops (i.e., the gap size), and if there were only 2 lattice sites (corresponding to 2 kb DNA) not extruded into loops for both gaps, we counted synapsis is being achieved and we stopped monitoring this site. We scored all DSB sites that were not initially constrained by LEFs as having failed to achieve synapsis, and did not continue monitoring these sites in later time steps.

### Modifications to LEF dynamics with additional mechanisms

With stabilization of LEFs at BEs, the LEFs with at least one motor subunit at the lattice sites representing BE had *w*-fold reduction in dissociation rate. With stabilization of LEF at DSB ends, the LEFs with at least one motor subunit at the lattice sites representing DSB ends had *r*-fold reduction in dissociation rate. With a small fraction *α*_*o*_ of long-lived LEFs, the long-lived LEFs had a dissociation rate 20 fold smaller than the normal LEFs (non long-lived LEFs). When there was a subpopulation of long-lived LEFs, the separation *d* referred to the separation of long-lived LEFs and normal LEFs combined. With targeted loading of LEFs at DSB, the loading probability at the lattice sites representing DSB ends was *F* fold higher than anywhere else on the chromosome.

### Simulated ChIP-Seq data

We first divided the genome into 5-kb bins, and then we counted the number of LEF motor subunits in each bin using the stored LEF positions 10 min post DSB, and wrote this to a BED file containing each bin’s score (i.e. number of LEFs). Along with another BED file containing the DSB coordinates, we used the plotHeatmap command from deepTools [69] to generate the ChIP-seq heatmaps shown in **Supplementary Fig. 5B**. For the fold enrichment calculated in **Supplementary Fig. 5A**, we counted the number of LEF motor subunits in each DSB-containing TAD across the different time points, and finally normalized the average LEF subunit counts post DSB by the corresponding average LEF subunit counts before DSB occurrence. The boundaries of the chr20 DSB-containing TAD of Dlva cells were determined by CTCF binding sites adjacent to the DSB site using CTCF ChIP-seq data [45].

## Data availability

Simulation data are available in the GitHub repository https://github.com/ahansenlab/DNA_break_synapsis_models/tree/main/Data.

## Code availability

Simulation and analysis codes, as well as analytical theory formulated in Mathematica are available in the GitHub repository https://github.com/ahansenlab/DNA_break_synapsis_models.

### Simulated Hi-C contact maps

We first normalized the LEF positions on the lattice sites by subtracting the positions of the closest DSB sites, so that the LEF positions were all relative to the closest DSB sites. We only included LEFs that were within ±2.5 Mb of the DSB sites. We then calculated the contact probability maps directly from the LEF positions, by utilizing a Gaussian approximation developed previously to simulate bacterial Hi-C maps [68]. Iterative correction was then applied to the calculated contact maps to generate the final contact maps [70]. Note in our 1D simulations, unlike experiments, targeted loading of LEFs to DSB ends continued even if they are already synapsed (this was so that the LEF abundance at each DSB sites did not depend on when the DSB sites are synapsed). Moreover, in experiments, the stripe pattern measured by Hi-C [45] might be weaker because certain DSBs sites could have already been synapsed/repaired in a fraction of the cells, whereas we did not simulate the DNA repair process downstream of synapsis.

## Supporting information

Supplementary Video 1 (successful synapsis)_NoExtendedMechanismHighProc_v5

Supplementary Video 2 (failed synapsis)_NoExtendedMechanism_v5

Supplementary Video 3_BEstabilization_v5

Supplementary Video 4_LonglivedLEFs_v5

Supplementary Video 5_DSBstabilization_v5

Supplementary Video 6_TargetedLoading_v5

Supplementary Video 7_MechanismCombined_v5

## Acknowledgement

We thank Dr. Leonid Mirny, Dr. Ed Banigan, Dr. Thomas Graham, Dr. Assaf Amitai, Dr. Kyoko Yokomori, Dr. Miles Huseyin, Sarah Nemsick and the Hansen lab for insightful discussions and comments on the manuscript. J.Y was supported by the MathWorks Engineering Fellowship and a graduate fellowship from the Ludwig Center at MIT’s Koch Institute for Integrative Cancer Research. This work was supported by the National Institutes of Health (grant numbers R00GM130896, DP2GM140938).

## Author Contributions

H.B.B. conceived of the project; J.Y., H.B.B and A.S.H designed the project; J.Y. performed the research with joint guidance from H.B.B. and A.S.H; J.Y. and H.B.B. developed the theory and simulation codes with input from A.S.H.; J.Y. performed the simulations and analyses with input from H.B.B and A.S.H; J.Y. drafted the manuscript with input from H.B.B. and A.S.H. and all authors edited the manuscript.

## Supplementary Information

### Supplementary Information

This file contains Supplementary Notes 1-3 and Supplementary Figures 1-6.

**Video 1**

Example of a successful synapsis event, simulated with the simple 3-parameter loop extrusion model. Simulation parameters: separation = 125 kb, processivity = 1000 kb, boundary strength = 0.5, and no mechanistic extensions.

**Video 2**

Example of a failed synapsis event, simulated with the simple 3-parameter loop extrusion model. Simulation parameters: separation = 125 kb, processivity = 1000 kb, boundary strength = 0.5, and no mechanistic extensions.

**Video 3**

Example video of synapsis with stabilization of LEF at BE. Simulation parameters: separation = 125 kb, processivity = 250 kb, boundary strength = 0.5, fold stabilization of LEF at BE = 16, and no other mechanistic extensions.

**Video 4**

Example video of synapsis with a small fraction of long-lived LEFs. Simulation parameters: separation = 125 kb, normal LEF processivity = 250 kb, boundary strength = 0.5, % long-lived LEFs = 20, long-lived:normal LEF processivity ratio = 20, and no other mechanistic extensions.

**Video 5**

Example video of synapsis with stabilization of LEF at DSB ends. Simulation parameters: separation = 125 kb, processivity = 250 kb, boundary strength = 0.5, fold increase in loading probability at DSB = 1000, and no other mechanistic extensions.

**Video 6**

Example video of synapsis with targeted loading of LEFs at DSB. Simulation parameters: separation = 125 kb, processivity = 250 kb, boundary strength = 0.5, fold stabilization of LEF at DSB ends = 4, and no other mechanistic extensions.

**Video 7**

Example video of synapsis with all four mechanistic extensions combined. Simulation parameters: separation = 125 kb, normal LEF processivity = 250 kb, boundary strength = 0.5, fold stabilization of LEF at BE = 16, % long-lived LEFs = 20, long-lived:normal LEF processivity ratio = 20, fold stabilization of LEF at DSB ends = 4, and fold increase in loading probability at DSB = 1000.

## Supplementary Information for

### 1 Supplementary Note 1

#### Overview

DNA Double-Strand Break (DSB) synapsis is generally assumed to be mediated by passive 3D diffusion [1]. However, due to the experimental difficulty associated with monitoring the spatio-temporal kinetics of the DSB repair process at a single locus in live mammalian cells by NHEJ (i.e. starting with DSB induction through synapsis and repair) [2], there has not been direct *in vivo* evidence indicating DSB synapsis is mediated solely by 3D diffusion.

DSB synapsis step can be described as a two-stage process [3–5]: the two DSB ends are first brought into proximity in a long-range (LR) synaptic complex, in which the two DSB ends are held ∼ 11.5 nm apart; the LR synaptic complex is then transitioned to a short-range (SR) synaptic complex that ligates the two DSB ends. While much has been uncovered about the processes downstream to the formation of the LR complex, comparatively less is known about the initial engagement of the two DSB ends, i.e., how the two DSB ends are brought into proximity in the LR complex in the first place. In our study, we use the term, synapsis, to refer to the process starting with DSB occurrence to the initial engagement of the two DSB ends in the LR synaptic complex. Given that autophosphorylation of the catalytic subunit of the DNA-dependent protein kinase (DNA-PKcs) is suggested to signal the transition from the LR to SR synaptic complex [4], the average synapsis time can be estimated around 6-11 minutes in mammalian cells based on prior experimental data [6]. To see if 3D diffusion is consistent with an average synapsis time of 6-11 minutes, we used an analytical formula for our calculations [7]. Based on [7], we estimate that synapsis would take on average between 20 to 90 minutes if mediated by passive 3D diffusion in chromosomes organized by SMC complexes into loops of average size ∼ 250 kb.

This raises a crucial point: assuming that loop extrusion contributes to DSB end synapsis through the combined action of constraining LEFs and gap-bridging LEFs as we propose, all current experimental estimates in mammalian cells already indirectly take into account the effects of loop extrusion. However, the speed and reliability of DSB end synapsis by pure 3D diffusion without the action of loop extrusion has then never been estimated before, and therefore the true efficiency of DSB end synapsis by pure 3D diffusion remains unknown.

The synapsis estimates based on pure 3D diffusion are significantly larger than the average synapsis time of 6-11 minutes in mammalian cells estimated from experimental data. Further, given that normal metabolism causes ∼ 1-50 DSBs per human cell per day [8, 9], in the nuclear “soup” of DSB ends, 3D diffusion alone cannot ensure the correct pairing of DSB ends. This points to the possibility of alternative mechanisms, beyond passive 3D diffusion, that help mediate DSB synapsis. The calculation of the DSB synapsis rates (and its limitations) using the analytical formula are described below.

#### 1.1 Estimation of the mean synapsis time for diffusion mediated synapsis

After a DSB occurs, recruitment of repair factors is initiated. However fast the recruitment process is, this duration of time gives the two DSB ends the opportunity to separate by diffusion. Indeed, experiments show that DSB ends can diffuse apart by several hundred nanometers [2, 10]. We asked whether 3D diffusion would be sufficient to bring the separated DSB ends back together within physiological time scales. Amitai and Holcman [7] derived an analytical formula to calculate the mean time, ⟨*τ*_*h*_⟩, for diffusion mediated synapsis in a confined volume:

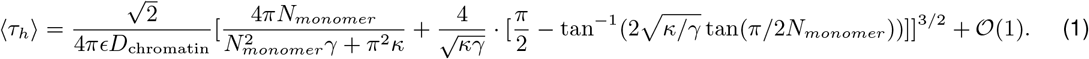

The encounter radius, *∈*, is the 3D distance the DSB ends must achieve in order for synapsis; here, we estimate *∈* = 11.5 ± 0.4 nm based on crystallographic models of the long-range synaptic complex that is thought to bring the two DSB ends into proximity [4, 5]; the uncertainty in our estimate comes from the resolution uncertainty associated with the crystal structure. The diffusion coefficient of chromatin in mammalian chromosomes has been estimated to be *D*_chromatin_ = (4 ± 1.9) × 10^*−*3^ μm^2^/s [11]. The number of monomers used to discretize the chromosome was calculated as, *N*_*monomer*_ = 2500, by assuming chromatin “monomers” of *δ* ≈ 30 nm fiber and that the DSB occurs inside of a loops of an average size of 250 kb, and using 0.3 nm/bp [12]. 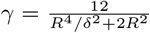 is a parameter dependent on the confinement radius *R* and the standard deviation of the distance between adjacent monomers *δ* [7]. Here we use a confinement radius of *R* = 990 ± 90 nm calculated from a broken locus in yeast [13], in agreement with another independent study [14], which is about twice of the confinement radius without a DSB [12–14]. *κ* = 3*/δ*^2^ is the spring constant. 𝒪 (1) is a constant error term. Given the aforementioned range of the chosen parameters, the mean synapsis time for synapsis driven by diffusion was calculated to be about 20-90 minutes.

#### 1.2 Estimation of the synapsis efficiency for diffusion mediated synapsis

With the average synapsis time of ⟨*t*_expt_⟩ ≈ 6 − 11 min and synapsis efficiencies estimated from experimental data *P*_synapsis,experiment_ ≈ 95%, we can estimate (as a first-order approximation) the upper bound on the total synapsis time, *t*_upper_, allowed by a cell:

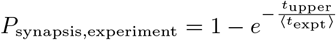

By setting *P*_synapsis,experiment_ = 0.95, and letting ⟨*t*_expt_⟩ = 11 min, we get ⟨*τ*_upper_⟩ ≈ 33 min as an estimate for the upper bound on time afforded by the cell to fix a DSB. Then, we can ask - given *τ*_upper_ = 33 min, how much does our calculated estimate for the first-passage time due to 3D diffusion decrease the probability of synapsis? Thus,

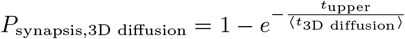

where ⟨*t*_3D diffusion_⟩ ≈ 20 − 90 min as calculated above. This calculation results in *P*_synapsis,3D diffusion_ ≈ 30 − 80%, suggesting that 3D diffusion alone is too error prone (i.e. failing 20% 70% of the time) to be consistent with the experimentally determined synapsis efficiency.

We end by emphasizing that essentially all of the above parameters are associated by great uncertainty and that our estimate therefore is too. Moreover, the above calculation assumes an equilibrium distribution of starting positions for the DSB ends, where as in a real system, the initial condition is that the two DSB ends are in close physical proximity. This assumption may partially reduce the first-passage times. However, the estimated ∼ 30 − 80% synapsis efficiency mediated by 3D diffusion above suggest that a purely diffusion driven synapsis mechanism is likely to be too error prone to be consistent with the *>* 95% efficiency of synapsis and NHEJ observed in mammalian cells.

### 2 Supplementary Note 2

#### Overview

To understand how the probability of DSB synapsis is affected by loop extruding factors (LEFs), we developed a probability theory framework for the process and used it to derive an analytical solution for the probability of synapsis. Briefly, we focus on the repair of DNA DSBs by non-homologous end joining (NHEJ). NHEJ repair involves two steps. First, the two DSB ends must be brought into proximity (synapsis). Second, they must be ligated back together. Here we focus on the first step, DSB synapsis. Please note that our goal is not to obtain the most precise analytical expression, but rather to derive a sufficiently accurate expression that we can use to obtain mechanistic intuition for how various loop extrusion mechanisms and parameters affect the efficiency of synapsis. Therefore we make approximations whenever necessary to simplify the mathematical form.

We first consider how the simplest loop extrusion model, which involves just 3 parameters, may facilitate DSB synapsis. After that, we extend this simple model by adding the four individual mechanisms discussed in the main text that improve synapsis efficiency: LEF stabilization by boundary elements (BEs); the presence of a subpopulation of long-lived LEFs; LEF stabilization by the DSB ends; and targeted loading of LEFs at DSB ends. Finally, we derive an analytical expression that combines all four additional mechanisms.

We note that our theory does not consider the effect of passive diffusion. Since passive 3D diffusion likely contributes to synapsis in cells in a manner that is synergistic with loop extrusion, we note that our estimates of DSB synapsis efficiencies should be considered lower bounds.

#### 2.1 The probability of synapsis mediated by loop-extruding factors

**Supplementary Note 2 Figure 1.**
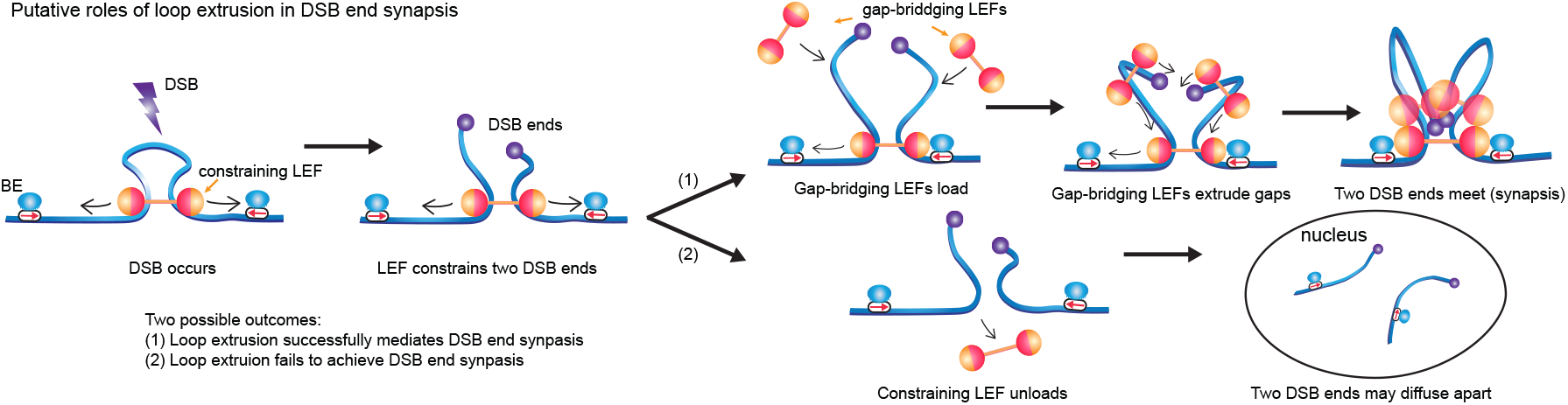
Pathway for successful LEF-mediated DSB synapsis. (1) Loop extrusion may facilitate successful DSB end synapsis in two steps: (i) the constraining LEF prevents the two DSB ends from diffusing apart after DSB and (ii) additional gap-bridging LEFs loaded within the loop extruded by constraining LEF can extrude sub-loops to bring the two DSB ends into proximity (2) If the constraining LEF falls off before the two DSB ends are brought into proximity by gap bridging LEFs, the two DSB ends may diffuse apart. In our simulations, we assume synapsis always fails once no constraining LEF remains for a given DSB.

We describe a general form for the probability of synapsis mediated by LEFs, *P*_synapsis_. For the purpose of understanding the limitations of what LEFs can and cannot do for synapsis, we omit contributions from 3D diffusion in all subsequent calculations of *P*_synapsis_.

Therefore, in order for LEF-mediated synapsis to happen, at least one constraining LEF must reside over the DSB site at the time when the DSB occurs. We denote the probability of having at least one constraining LEF as *P*_constrained_, which can be calculated as the probability that the DSB occurs inside a DNA loop extruded by a LEF. If the condition of having at least one constraining LEF over the DSB is met (**Supplementary Note 2 Figure 1, left**), then additional LEFs loaded into the gap between the DSB and the edges of the constraining LEF can extrude loops to bring the DSB ends into proximity to achieve synapsis (**Supplementary Note 2 Figure 1 path (1)**. We refer to this process as gap-bridging and the LEFs that mediate gap-bridging as gap-bridging LEFs. We note that at least one constraining LEF needs to remain throughout the gap-bridging process until synapsis is simultaneously achieved on both sides. If the constraining LEF dissociates before gap-bridging on both sides is achieved, we assume that the two DSBs ends can diffuse apart, and we consider this a failed LEF-mediated synapsis event (**Supplementary Note 2 Fig. 1 path (2)**). We denote the conditional probability of finishing gap-bridging given that the DSB ends are constrained at the time of DSB occurrence as *P*_end-joining|constrained_.

Thus, the probability of synapsis, *P*_synapsis_, can then be expressed as the product of *P*_constrained_ and *P*_end-joining|constrained_:

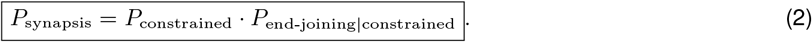

In the following sections, we derive analytical estimates for both *P*_constrained_ and *P*_end-joining|constrained_.

#### 2.2 The probability of DSB ends being constrained

As stated above, *P*_constrained_ is simply the probability that the DSB occurs inside a DNA loop such that the broken DSB ends remain constrained by at least one LEF. Since we assume that DSBs occur homogeneously throughout the genome, *P*_constrained_ is equivalent to the fraction of the genome that is extruded into loops by LEFs. For the scenario without BEs, the fraction of the genome inside loops can be estimated by considering the nesting of LEFs [15]:

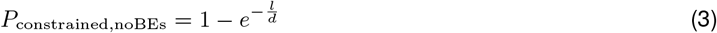

where *l* is the average DNA loop size, and *d* is the LEF separation, i.e., the average linear distance between LEF loading sites. This expression accounts for the increasing chance of LEFs loaded into existing loops and thus not contributing to increasing the fraction of genome inside loops.

To incorporate the effect of BEs on *P*_*constrained*_, we account for how BEs decrease the amount of DNA that is inside loops by prematurely stalling LEFs. Therefore, the fraction of genome extruded into loops becomes:

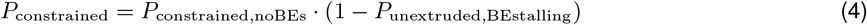

where *P*_unextruded,BEstalling_ is the probability of DNA becoming unextruded (i.e. unlooped) due to stalling of LEFs at BEs.

To calculate *P*_unextruded,BEstalling_, we adapted the mean-field theoretical model previously derived by Banigan and Mirny [16] for the fraction of genome coverage by loops. Therefore,

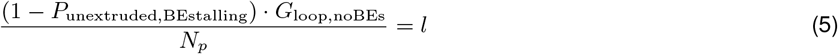

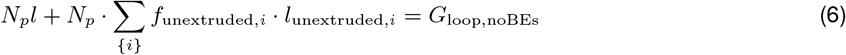

where *G*_loop,noBEs_ is the average length of genome inside loops with no BEs present, and thus if the total genome length is *G, G*_loop,noBEs_ = *G P*_constrained,noBEs_. The fraction of *G*_loop,noBEs_ that remains looped in the presence of BEs is given by 1 − *P*_unextruded,BEstalling_. *N*_*p*_ is the total number of parent loops as defined in [16] (i.e. it is the total number of loops minus the number that exist within a larger loop). *f*_unextruded,*i*_ is the probability of adjacent parent LEFs being in a state *i*, where the set of possible {*i*} constitutes all configurations of two adjacent LEFs and BEs which result in unextruded DNA between the LEFs because of the presence of BEs (see **Supplementary Note 2 Figure 2**). Finally, *l*_unextruded,*i*_ is the average length of unextruded DNA for the configuration *i*. Here the size of parent loops are approximated as the average loop size of all LEFs. Combining Eqs.(5)-(6), we can solve for *P*_unextruded,BEstalling_:

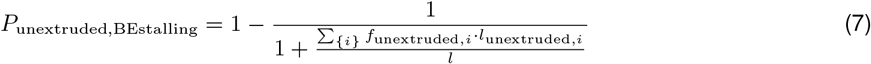

The unknown quantities now become *f*_unextruded,*i*_ and *l*_unextruded,*i*_ which we can calculate by enumerating the types of configurations which lead unextruded DNA because of a BE (**Supplementary Note 2 Figure 2**). Instead of explicitly enumerating and calculating all the possible configurations, we take a mean field approach and consider the leading three configurations (**Supplementary Note 2 Figure 2**) as being representative of the much larger space of configurations.

We first consider the case where the loading sites of two adjacent LEFs are separated by a BE (i.e. are in adjacent topologically associated domains, (TADs)). We refer to this case as Configuration I. In our mean field approach, this scenario arises if the average distance between the BE and the loading site of LEFs (i.e. 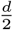) is smaller than the TAD size, *D*, i.e.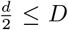 (**Supplementary Note 2 Fig. 2A**). In this case, unextruded DNA arises if and only if one LEF is stalled by BEs and the other LEF fails to reach BE. The probability of a LEF motor subunit being stalled by BEs can be estimated by the fraction,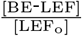, where [BE-LEF] is the concentration of stalled LEF-BE complexes and [LEF_o_] is the total density of LEF motor subunits extruding in one direction:

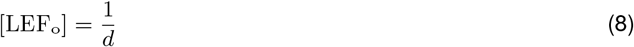

Thus the overall fraction of LEFs in *Configuration I* can be calculated as the following:

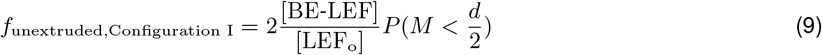

where *M* is the random variable representing the extrusion distance of one LEF motor subunit before the LEF unloads. The pre-factor of 2 accounts for the fact that the LEF stalled by BE could be either on the left or on the right of the BE. If we assume *M* is exponentially distributed, since each LEF motor subunit (a LEF is composed of two subunits) on average travels *l/*2, then *M* ∼ Exp(*l/*2). Thus the probability that *M* is smaller than 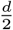 can be calculated as:

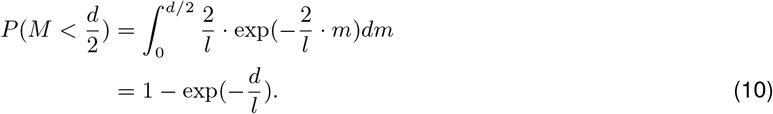

The corresponding average unextruded DNA length can be calculated as the following (**Supplementary Note 2 Fig. 2A**):

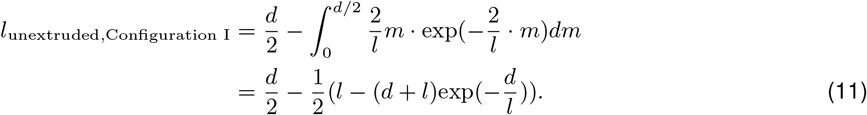

**Supplementary Note 2 Figure 2.**
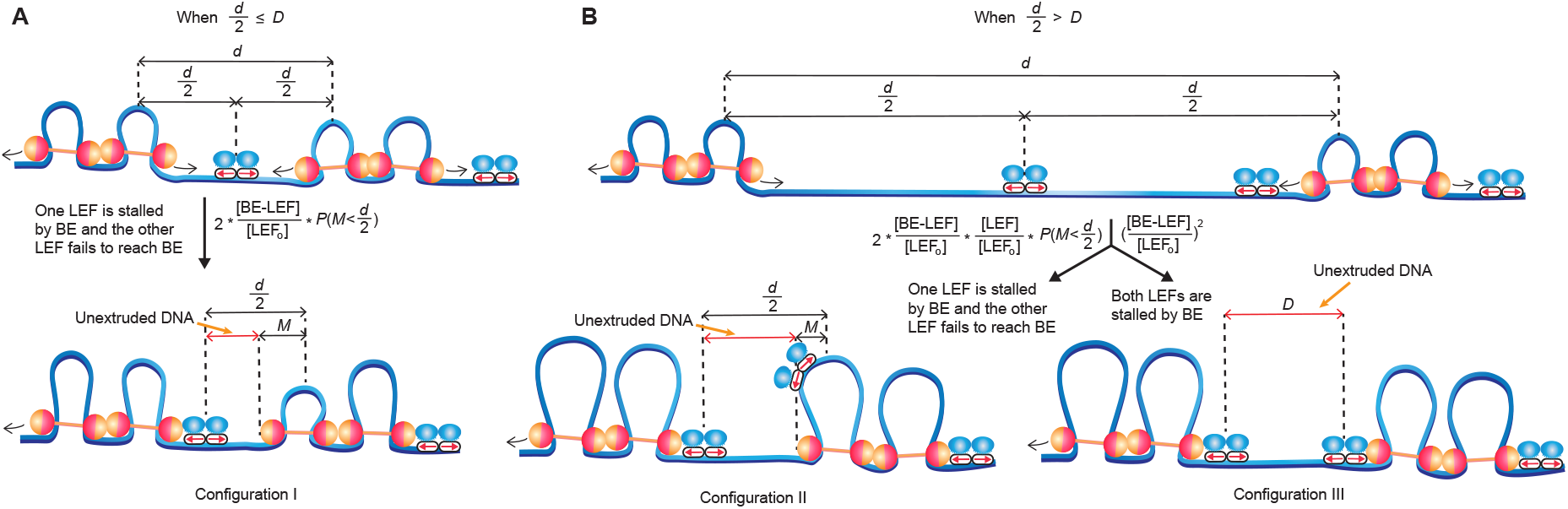
BE stalling can generate three types of LEF pair configurations leading to unextruded DNA. (**A**) When 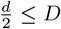, the loading sites of the LEF pair are in adjacent TADs, separated by a distance of *d*. On average, the BE is equidistant to both LEF loading sites. Unextruded DNA arises if and only and if one LEF is stalled by BE and the other LEF fails to reach BE, leading to an unextruded segment of average length calculated in Eq.(11).(**B**) When 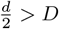, the loading sites of the LEF pair are separate by a TAD. Unextruded DNA can arise with two different LEF pair configurations. The first configuration is identical to the configuration in (A), which occurs when one LEF is stalled by BE, and the other LEF is not stalled by the additional BE in between but fails to reach the leftmost BE. The second configuration occurs when both LEFs are stalled by BEs, resulting in an unextruded segment with the same length as the TAD size.

Next we consider the cases where the loading sites of the adjacent LEFs are separated by a TAD (i.e. two BEs). This occurs when 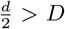. There are altogether two such types of configurations which we call Configuration II and Configuration III. Configuration II, occurs when one LEF is stalled by one of the BEs forming a BE-LEF complex, whereas the other LEF is not stalled, but fails to reach the BE-LEF complex thereby leaving an unextruded gap (**Supplementary Note 2 Fig. 2B**, left branch). Let [LEF] be the concentration of extruding LEFs not bound to BE. Configuration II type of scenarios occur with the following frequencies and result in the following lengths of unextruded DNA:

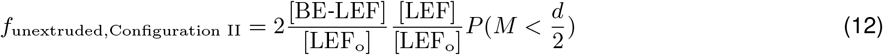

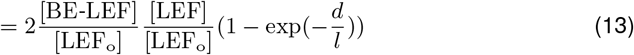

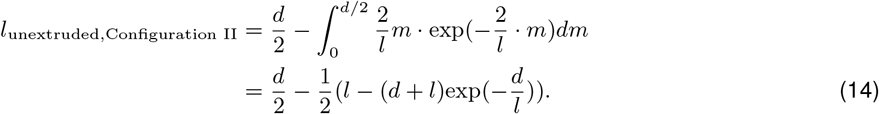

Configuration III arises when both LEFs are each stalled by one of the BEs, leading to an unextruded DNA segment of average length that equals the TAD size *D* (**Supplementary Note 2 Fig. 2B**, right branch). Note that while in reality there could be more than one TAD is in between the loading sites of two adjacent LEFs, we do not consider those scenarios within our parameter space under mean-field theoretical model, since the largest separation *d* we consider is 500 kb, and the smallest total size of two adjacent TADs in our simulations is 600 kb (200 kb and 400 kb TAD next to each other), and thus on average there would not be more than one TAD between the loading sites of two adjacent LEFs. Configuration III type of scenarios occur with the following frequencies and result in the following lengths of unextruded DNA:

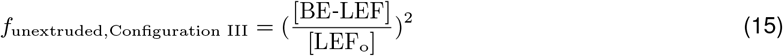

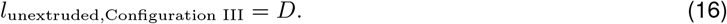

Combining Eqs.(3)-(4),Eqs.(7)-(16), we obtain the expression for *P*_constrained_:

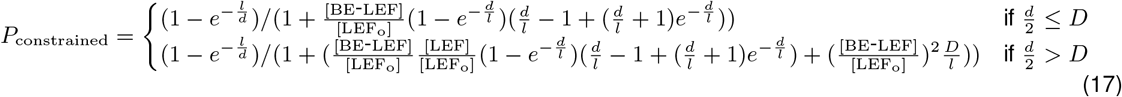

The average DNA loop size *l* has been previously estimated using processivity *λ* and separation *d* with ∼ 1% precision for 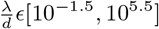, by applying the following expression obtained through fitting a 7-th degree polynomial to simulation results [15]:

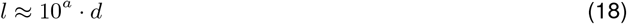

where:

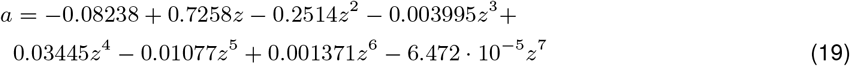

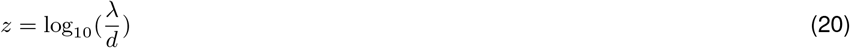

Now the only unknown in Eq.(17) is 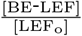. To calculate 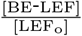, we consider the following system:

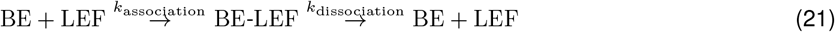

where BE-LEF is the complex of BE and LEF formed upon a LEF being stalled by a BE, with rate constant *k*_association_. LEF in complex with BE dissociates from the DNA and reloads on the genome with rate constant *k*_dissociation_. We assume one BE can only associate with one LEF at a time. Although we consider two-sided LEFs, the extrusion along both directions are identical and independent processes, thus we can consider the model in Eq.(21) only for extrusion in one direction, without loss of generality.

Since the rate of BE-LEF formation is determined by the rate of collision between BE and LEF, the association rate constant *k*_association_ can be written as the following:

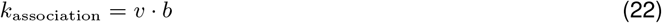

where *v* is the unobstructed extrusion speed in one direction (i.e., 1/2 the total extrusion speed), and *b* is boundary strength, defined as the probability of BE stalling LEF extrusion upon encountering (with probability 1 − *b*, LEF extrudes past BE).

Once a LEF is stalled by a BE, the average duration before the LEF dissociates from the DNA is given by 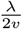, where *λ* is the processivity, i.e., the average length of DNA extruded by an unobstructed LEF. Thus the dissociation rate constant *k*_dissociation_ can be written as the following:

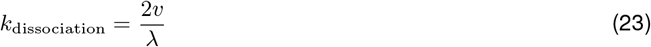

We can write the following differential equation to describe the rate of formation of BE-LEF:

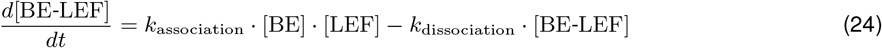

At steady state, we have:

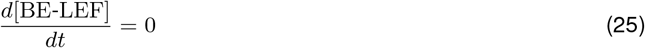

Combining Eq.(22)-Eq.(25), we get:

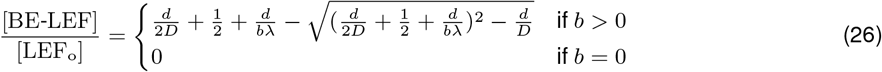

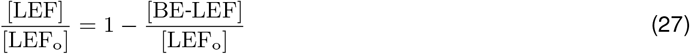

Now we can substitute Eqs.(26)-(27) into Eq.(17) to solve for *P*_constrained_. In the simulations we have four different TAD sizes of *D*_*j*_ *∈*{200 kb, 400 kb, 800 kb, 1200 kb}, each of which appears with a frequency of *ω*_*j*_ *∈*{0.5, 0.25, 0.125, 0.125} respectively, and thus *P*_constrained_ can be computed by summing Eq.(17) with different TAD sizes weighted by the frequency of the TAD size (assuming *b >* 0):

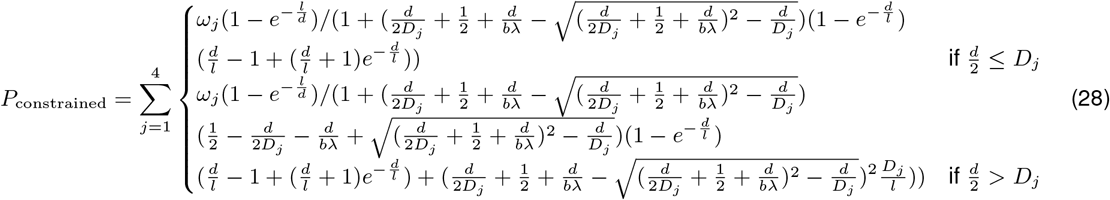

The theoretical expression we derived for *P*_constrained_ in Eq.(28) can predict the percentage of DSB sites constrained by LEFs with reasonably high accuracy (**Supplementary Note 2 Fig. 3**). For the rest of the paper, we use the weighted average TAD size of 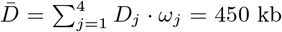 · *ω*_*j*_ = 450 kb to simplify equations and calculations, unless specified otherwise.

**Supplementary Note 2 Figure 3.**
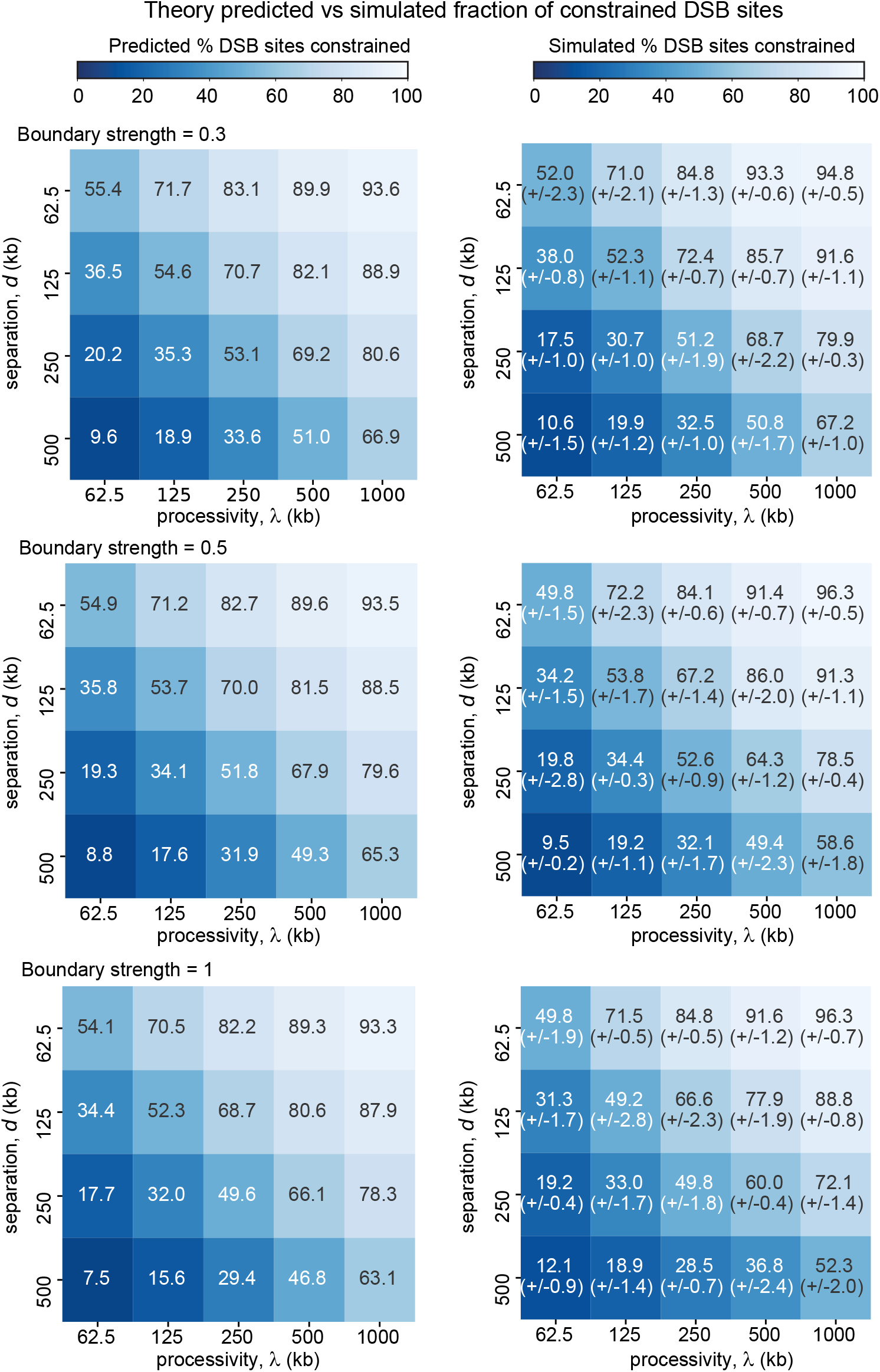
Theory prediction of the fraction of DSB sites constrained by LEFs is consistent with simulation results across different boundary strengths. The middle panel with boundary strength = 0.5 is the same as main text **Fig. 3B**. *P*_constrained_ can be predicted given processivity, separation, and boundary strength. Heatmaps of predicted (left) and simulated (right; numbers in brackets show standard error of mean, n=3 independent simulations, with 218 DSB events per simulation) fraction of constrained DSB sites with different combinations of processivity (y axis) and separation (x axis). Each row corresponds to boundary strengths of 0.3, 0.5 and 1 respectively.

##### 2.2.1 Simplified expression for the probability of DSB ends being constrained

By using a linear approximation (tangent line approximation) of 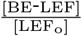, we can simplify the expression for *P*_constrained_. To this end, we define:

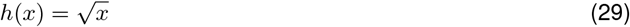

Then the linear approximation of *h*(*x*), H(*x*), can be written as:

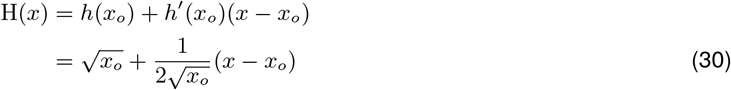

Substituting 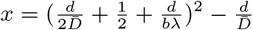 and 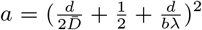 into Eq.(30), we obtain the linear approximation of Eq.(26):

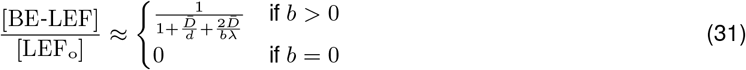

Substituting Eq.(27) and Eq.(31) into Eq.(17), we get the simplified expression for *P*_constrained_:

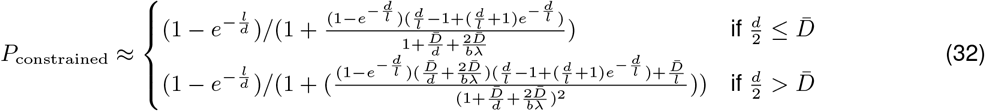

Eq.(32) shows *P*_constrained_ is a monotonic increasing function of *l* and *λ*, and a monotonic decreasing function of *d*.

#### 2.3 The probability of end joining given that DSB ends are constrained

To calculate the probability of gap-bridging given that DSB ends are constrained, *P*_end-joining|constrained_, we want to compute how often simultaneous gap-bridging on both sides of the DSB happens before the constraining LEF unloads. In other words, we want to determine how frequently the time it takes to achieve synapsis is shorter than the lifetime of the constraining LEF. Since gap-bridging on both sides of the DSB are independent of each other, it is more mathematically tractable to consider the probability of gap-bridging on each side, and then take into account that synapsis requires the gap-bridging on both sides of the DSB to happen at the same time. Therefore, we need to formulate the time to bridge the gap and the lifetime of constraining LEFs respectively, which we will discuss next.

##### 2.3.1 The gap-bridging time distribution

We define the gap-bridging time, *T*, as the duration between DSB occurrence and the first time that the gap is bridged on one side of the DSB. Thus, let the first-passage time, *T*, be a random variable and the probability distribution of *T* be given by *f*_*T*_. To simplify the theory, we initially assume there is no gap-bridging LEFs present between the constraining LEF and DSB ends at the time of DSB occurrence (we modify this assumption later on considering the full probability of gap-bridging), and we assume only one gap-bridging LEF carries out the bridging from beginning to end. Therefore, we can conceptualize gap-bridging on one side of the DSB for a given DSB site as a two-step process: first, a gap-bridging LEF must load between the DSB end and the constraining LEF; second, the gap-bridging LEF must extrude to bridge the gap between the DSB end and the constraining LEF. We define the loading time random variable as *X*, corresponding to the time it takes to finish the first step, and the extrusion time random variable as *Y*, corresponding to the time it takes to complete the second step. Thus,

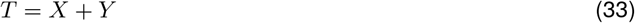

###### 2.3.1.1 The loading time distribution

Generally, we can assume that the process of loading gap-bridging LEFs into the gap between DSB end is Markovian, and thus exponential, with parameter, *k*_load_ = ⟨*τ*_load_ ⟩^*−*1^, where *k*_load_ is the LEF loading rate, and ⟨*τ*_load_⟩ is the average loading time. We assume LEFs reload randomly (uniform loading probability across the genome) as soon as they unload.

Let, *L*, be the length of the gap, i.e., the length of DNA between the DSB end and the edge of the constraining LEF. We assume the LEF lifetime is exponentially distributed with an average lifetime of 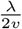. The probability density function(PDF) of the loading time *X*, defined on the interval *x∈* [0, ∞), can be described by the following exponential function:

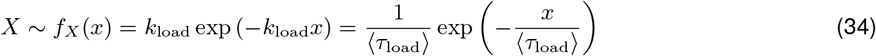

in which:

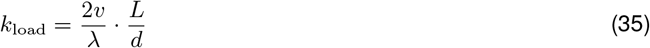

where, as a reminder, *v* is the extrusion speed in one direction (i.e., 1/2 the total extrusion rate), *λ* is the processivity (i.e., the average length of DNA extruded by an unobstructed LEF), *L* is the length of the DNA between the DSB end and the edge of the constraining LEF, and *d* is the average linear distance between LEF loading sites.

As can be told from Eq.(35), the longer the gap, *L*, between the DSB and the edge of the constraining LEF, the faster the loading of a gap-bridging LEF.

###### 2.3.1.2 The extrusion time distribution

Provided that a gap-bridging LEF has been loaded in the gap, we can then determine the distribution of extrusion times. Since we assume that only one gap-bridging LEF extrudes the gap, the extrusion time is determined by the time it takes to extrude to whichever of the DSB and constraining LEF is furthest away: 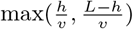 where *h∈* [0, *L*] is the loading point of the LEF, as shown in the diagram below. Thus, the minimum extrusion time of 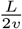, is achieved when the gap-bridging LEF loads right in the center of the gap, whereas the maximum extrusion time of 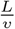 is achieved when the gap-bridging LEF loads at either boundary of the gap. Since we assume the loading of the LEFs is spatially homogeneous, the extrusion time 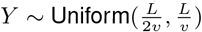.

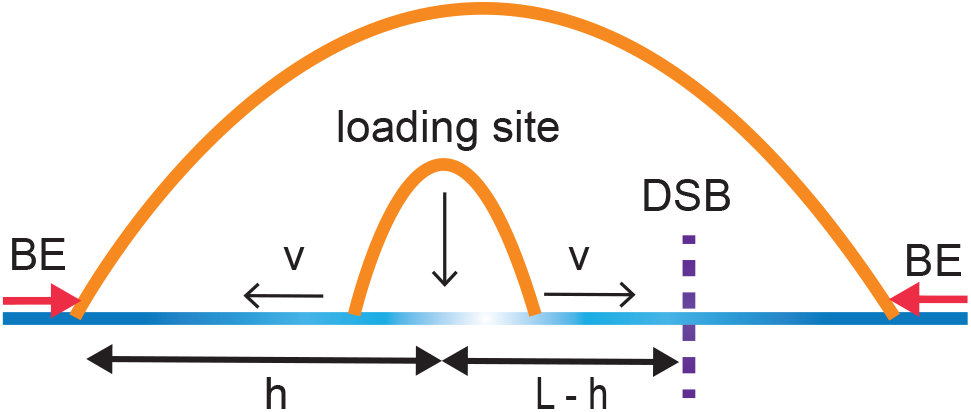

Thus, the PDF of the extrusion time *Y* is given by the following uniform distribution:

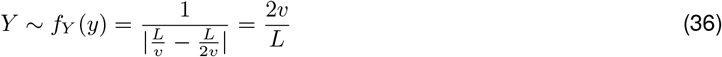

defined on the interval 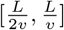.

We can rewrite Eq.(36) as the following to simplify expressions later on:

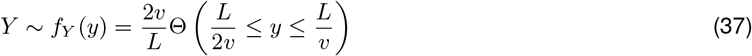

where,

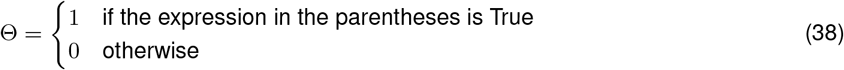

###### 2.3.1.3 Computing the gap-bridging time distribution through convolution

Having determined the loading time distribution, *f*_*X*_, and the extrusion time distribution, *f*_*Y*_, we can now compute *T* = *X* + *Y*. To do so we need to compute the convolution of *f*_*X*_ and *f*_*Y*_:

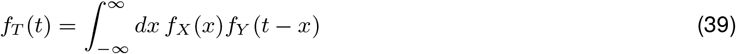

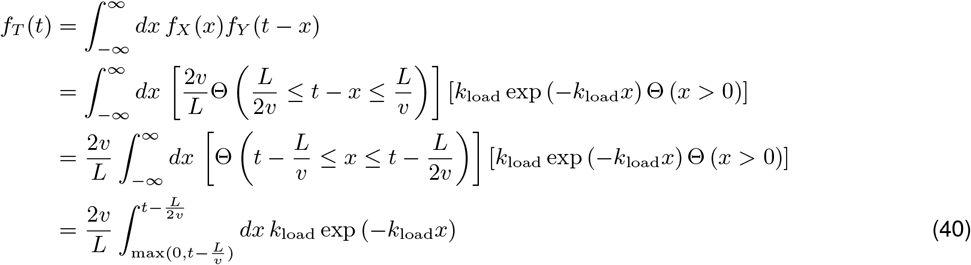

The solution has two parts, making it piece-wise continuous:

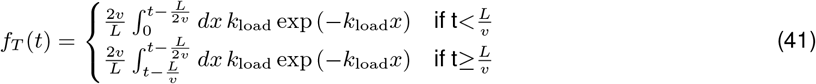

which results in:

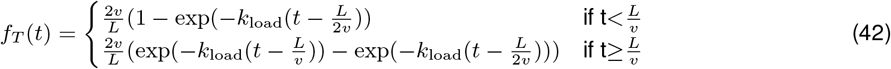

##### 2.3.2 The constraining LEF lifetime distribution

Since gap-bridging must take place before the constraining LEF unloads, we need to also compute the distribution of the constraining LEF lifetime. Let the lifetime of constraining LEFs be given by the random variable *C*. The probability distribution of *C* is given by *f*_*C*_. As above, we assume the lifetime of constraining LEFs are exponentially distributed, and ⟨*τ*_*c*_⟩ is the average lifetime of constraining LEFs. The PDF of the constraining LEF lifetime *C* can be expressed as:

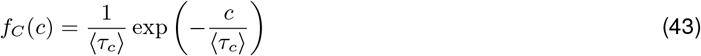

in which:

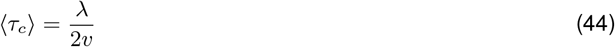

##### 2.3.3 The probability of gap-bridging on one side of the DSB

Having determined the gap-bridging time distribution *f*_*T*_, and the constraining LEF lifetime distribution *f*_*C*_, we can now compute the probability of gap-bridging on the side of the DSB that is bridged first. The calculation of *f*_*T*_ above assumes there is no gap-bridging LEF present at the time of DSB occurrence. However, there is a nonneglibible probability that gap-bridging LEFs are already present in the gap. The average number of gap-bridging LEFs within the constraining LEF, *n*, can be approximated as the following using Eq.(18) [15]:

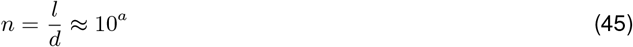

where *a* is defined in Eq.(19)-(20).

Let the number of gap-bridging LEFs inside the constraining LEF, *N*, be a random variable. Then *N* follows Poisson distribution:

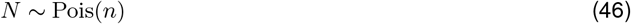

Thus the probability of having no gap-bridging LEFs inside the constraining LEF is:

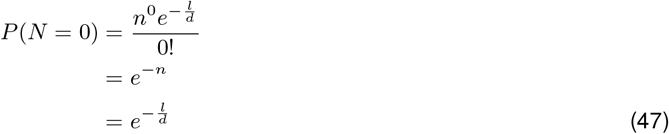

Conversely, the probability of having one or more gap-bridging LEFs is:

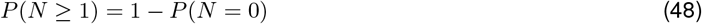

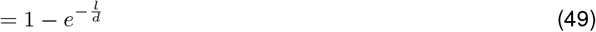

As an approximation, we assume if one or more gap-bridging LEFs are present within the constraining LEF at the time of DSB occurrence, then one gap will be successfully bridged with probability 1. This approximation is motivated by the observation that the loading of gap-bridging LEF is the rate-limiting step of synapsis (**Supplementary Fig. 3**). If one or more gap-bridging LEFs are present within the constraining LEF, one of the two gaps are likely closed by the pre-existing gap-bridging LEFs. However, if initially there is no gap-bridging LEF inside the constraining LEF, the gap-bridging time *T* must be smaller than the constraining LEF lifetime *C*. We note *f*_*T*_ is a function of the gap length *L*, and we denote the gap length on the side bridged first as 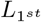. Therefore,

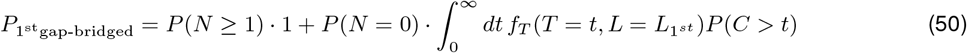

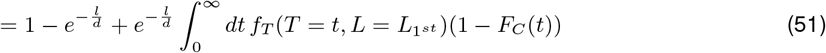

where *F*_*C*_ is the cumulative distribution function of the constraining LEF lifetime:

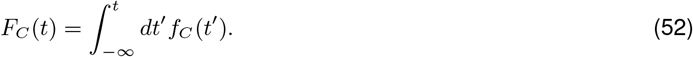

Because *f*_*T*_ is piece-wise continuous, we can compute this by breaking it up into two parts:

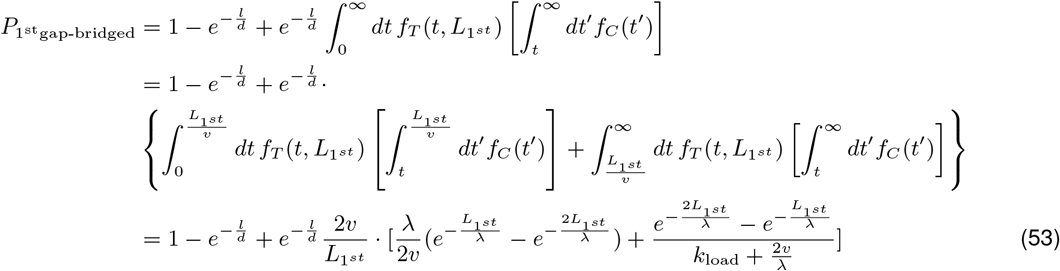

##### 2.3.4 Simultaneous gap-bridging on both sides of the DSB and the gap-bridging LEF lifetime distribution

In order to achieve synapsis, the gaps on both sides of the DSB must be bridged simultaneously. In other words, once the gap on one side of the DSB is bridged, the gap on the other side of the DSB must be bridged before the gap-bridging LEF on the side bridged first unloads. Let the lifetime of gap-bridging LEFs that have already finished gap-bridging on the side of DSB bridged first be given by the random variable *G*, whose PDF is *f*_*G*_. Then the probability of gap bridging on the second side (while the gap-bridging LEF on the first side remains bound) can be written as:

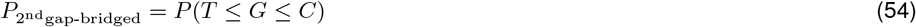

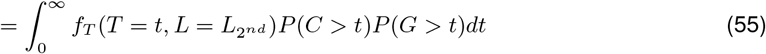

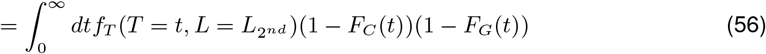

where:

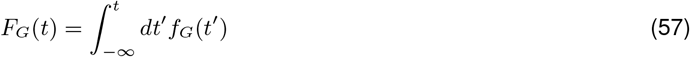

Utilizing the memoryless property of exponential distribution, the PDF of the lifetime *G* of the gap-bridging LEF that have already finished gap-bridging on the side bridged first, can be expressed as:

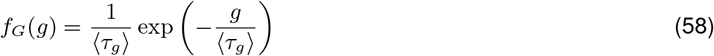

in which:

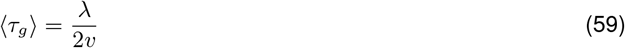

Note the PDF of gap-bridging LEF lifetimes is mathematically identical to the PDF of constraining LEF lifetimes, since these two kinds of LEFs are essentially identical, and their identity are assigned based on their locations relative to a DSB. We assign different notations here to facilitate our discussions of extensions to the loop extrusion model later where distinguishing them becomes helpful. Now we can compute the probability of bridging the gap on the other side of the DSB while the gap-bridging LEF on the side bridged first remains:

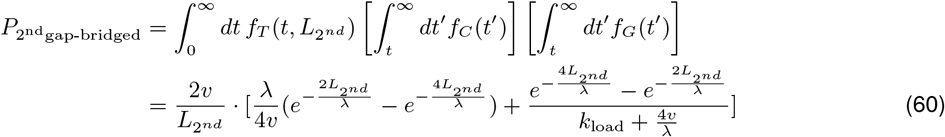

##### 2.3.5 Computing the probability of joining DSB ends given DSB ends are constrained

Since gap-bridging on the first side and the second side of the DSB are independent processes, the probability of both gaps being bridged upon the first try is the product of Eq.(53) and Eq.(60). However, gap-bridging does not need to be achieved upon the first try: as long as the constraining LEF remains, even if the gap-bridging LEF on the side bridged first unloads, the gap-bridging process can continue until simultaneous gap-bridging on both sides of the DSB is fulfilled. Thus the probability of joining DSB ends given DSB ends are constrained for a given DSB site whose gap lengths are 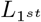 and 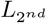 can be written as the following infinite sum:

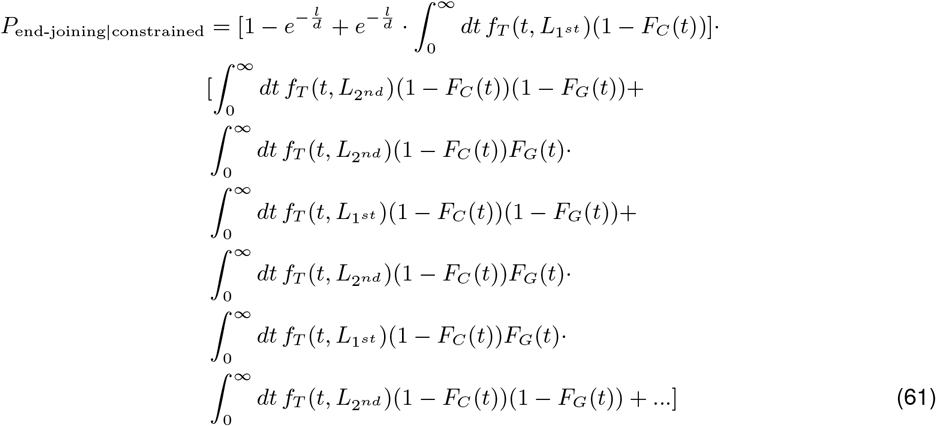

Utilizing the formula for the infinite sum of a geometric series, the equation above can be reduced to:

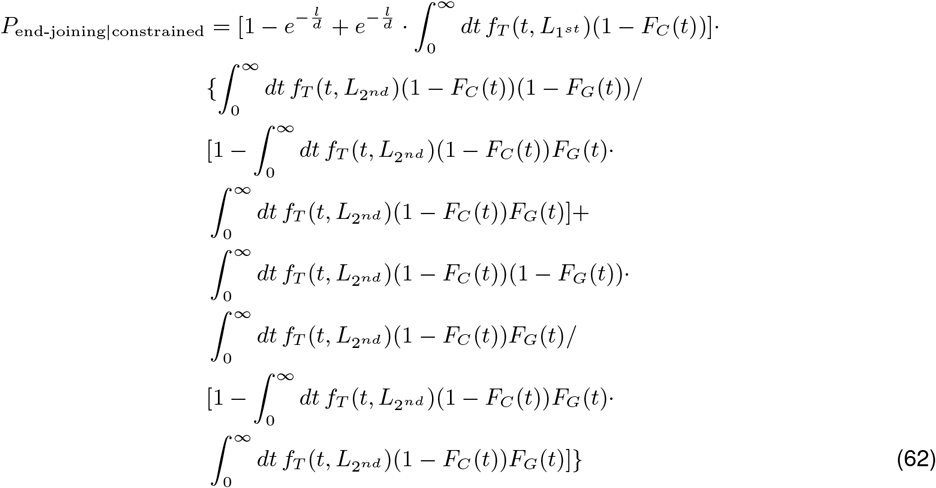

Eq.(62) is written for a specific DSB site with gap lengths of 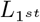 and 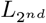 for the side bridged first and the side bridged second respectively. We assume the total length of the gaps on both sides of the DSB (the sum of 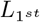 and 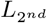), is the average DNA loop size *l*:

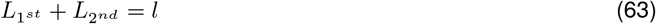

We can then integrate over 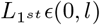 to generalize Eq.(62) for any DSB site, given that the location of DSB is random:

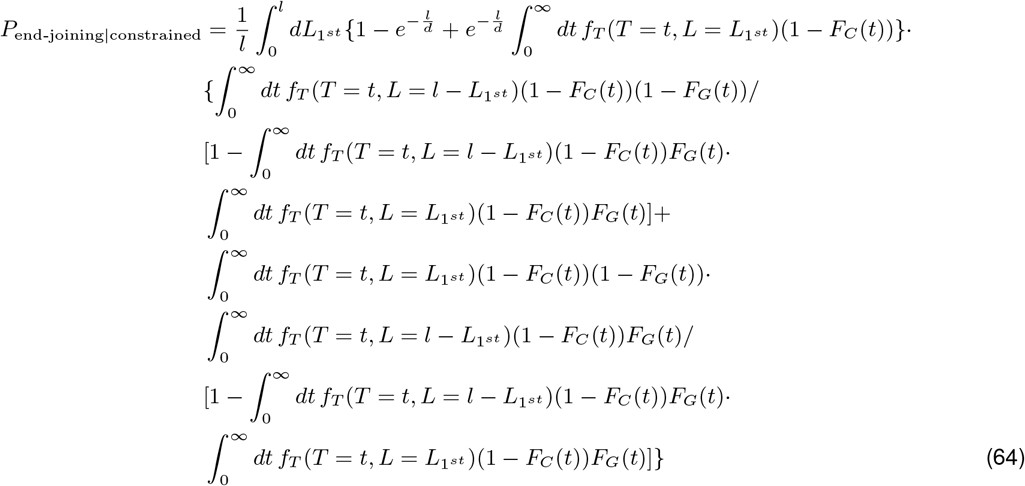

We define:

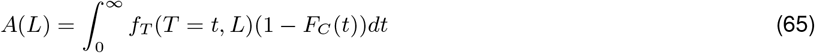

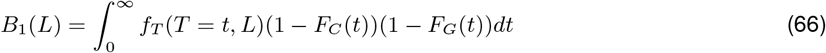

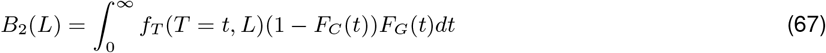

Thus Eq.(64) can be rewritten as:

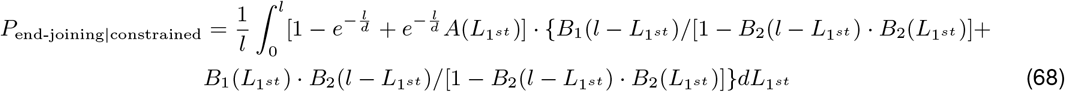

We have computed Eq.(65) and Eq.(66) for specific gap lengths of 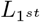 and 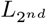 respectively in Eq.(53) and Eq.(60). We rewrite them below for the general gap length of *L*:

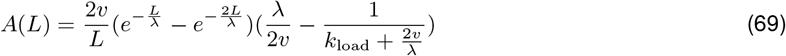

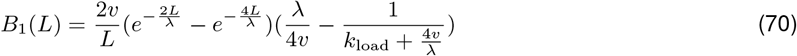

Notice Eq.(67) is simply the difference between *A*(*L*) and *B*_1_(*L*):

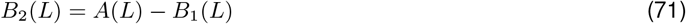

Now we can substitute Eqs.(18)-(20) and Eqs.(69)-(71) into Eq.(68) and perform numerical integration to obtain *P*_end-joining|constrained_.

The probability of synapsis *P*_synapsis_ can then be determined by multiplying the numerically integrated *P*_end-joining|constrained_ and the *P*_constrained_ computed in Eq.(28). Note that *P*_end-joining|constrained_ and *P*_constrained_ both only depend on the LEF processivity *λ* and the LEF separation *d*, and thus *P*_synapsis_ can also be determined as long as we know *λ* and *d*. Below we compare the synapsis efficiency predicted by our analytical expression for *P*_synapsis_ and the simulated synapsis efficiency with different combinations of *λ* and *d*. This comparison shows that our theory is reasonably accurate (**Supplementary Note 2 Fig. 4**).

**Supplementary Note 2 Figure 4.**
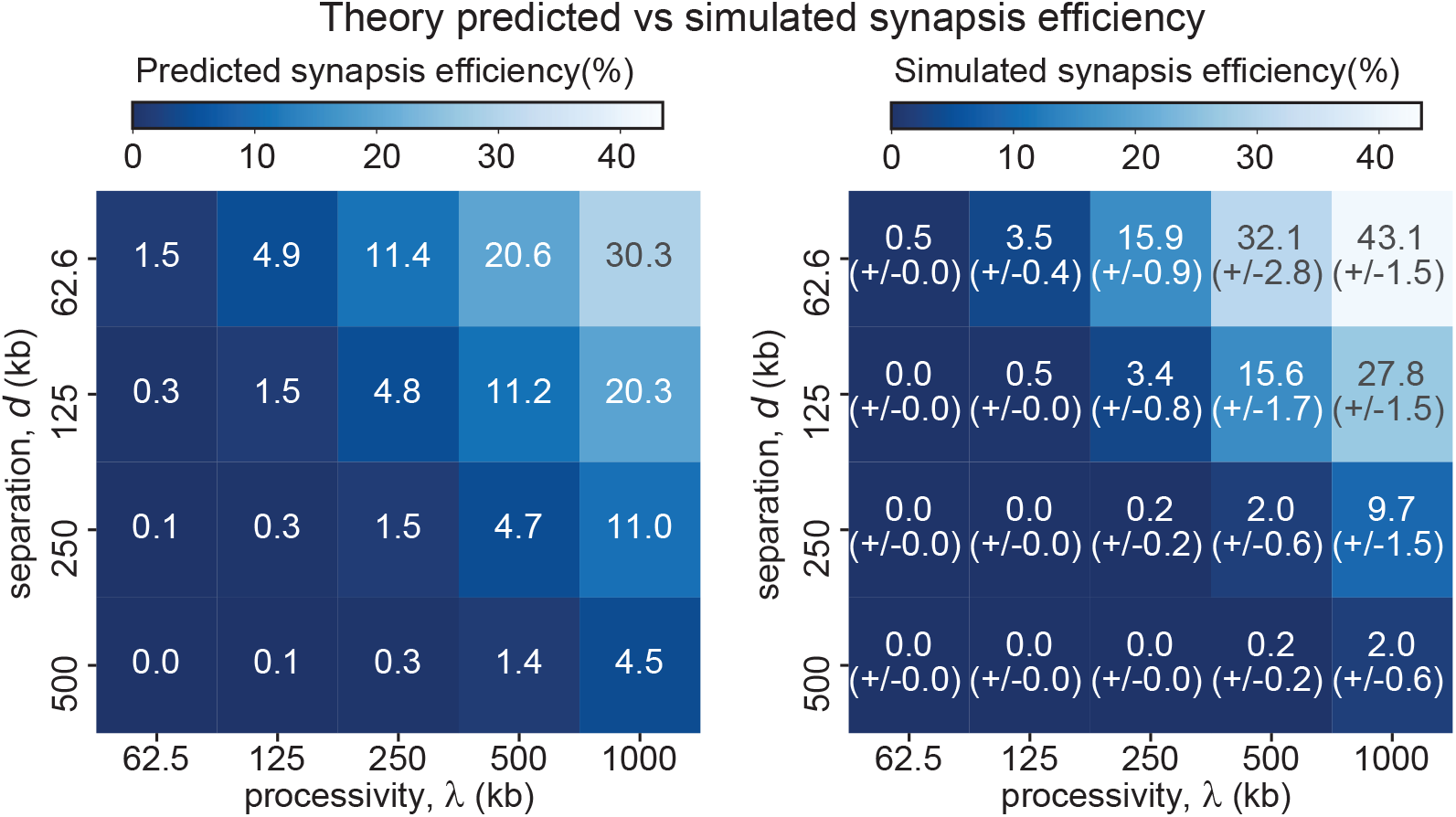
Theory prediction of synapsis efficiency is consistent with simulation results. Same as **Supplementary Fig. 3**. Heatmaps of predicted (left) and simulated (right; numbers in brackets show standard error of mean, n=3 independent simulations, with 218 DSB events per simulation) synapsis efficiency with different combinations of processivity (y axis) and separation (x axis). Boundary strength = 0.5 was used in the simulations.

##### 2.3.6 Simplified expression for the probability of end-joining given the DSB is constrained and two important relative timescales

Our theory above highlights that *P*_synapsis_ can be determined from just *λ* and *d*. However, despite this seemingly simple picture and the reasonable agreement between our theory and simulations, our theoretical formulation for *P*_synapsis_ is still too complicated to be written in a compact form. Therefore, it is challenging for someone to gain mechanistic intuition for what factors underlie efficient synapsis. In this section, we aim to simplify the expression for *P*_synapsis_ with the goal of gaining mechanistic intuition.

Let us consider the scenario where gaps are bridged upon the first try. Further, let us consider a DSB site with the break right in the center of the constraining LEF, and thus with gap length 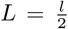 on both sides of the DSB. We assume there is no gap-bridging LEF present in the constraining LEF at the time of DSB occurrence. Then *P*_end-joining|constrained_ can be approximated as the following expression simplified from the product of Eq.(53) and Eq.(60):

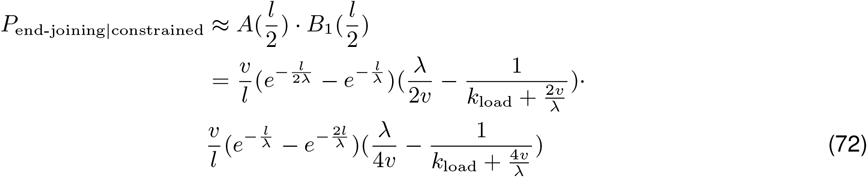

We can rewrite Eq.(72) as:

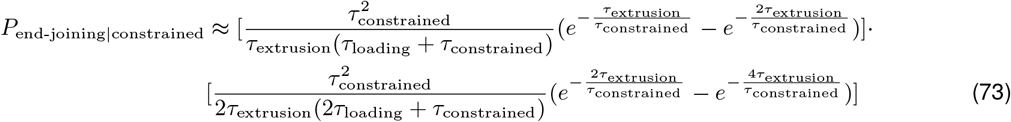

where:

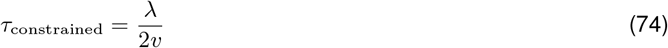

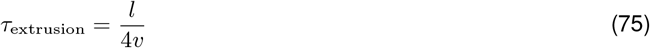

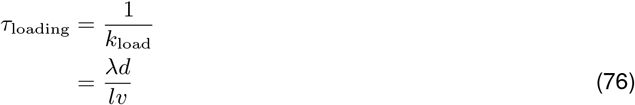

We can further simplify Eq.(73) by defining:

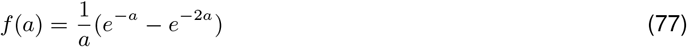

Then Eq.(73) can be written as:

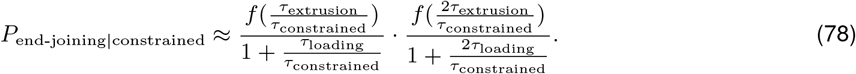

The simplified formula for *P*_gap-bridged|constrained_ suggests that given the DSB ends are constrained, synapsis efficiency is dictated by two relative timescales: the ratio of loading time and the constraining LEF lifetime and the ratio of extrusion time and the constraining LEF lifetime. Further, *P*_gap-bridged|constrained_ monotonically decreases with the two relative timescales, so decreasing either or both of the two relative timescales will improve synapsis efficiency.

Substituting Eq.(32) and Eq.(78) into Eq.(2), we can obtain a final simplified expression for the synapsis efficiency:

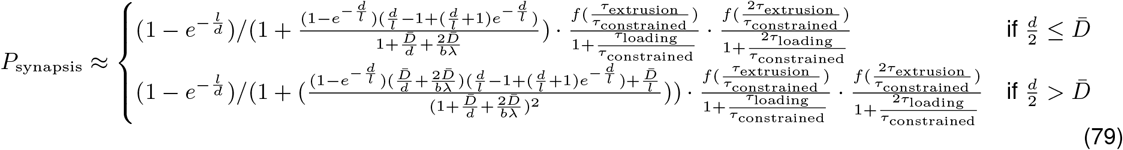

#### 2.4 Synapsis with additional mechanisms

In this section, we will discuss four experimentally plausible extensions of the simple loop extrusion model that improve synapsis efficiency: stabilization of LEFs by BEs, the presence of a small fraction of long-lived LEFs, stabilization of LEFs by DSB ends, and targeted loading of LEFs to DSBs. We start by investigating how the addition of each of the four mechanisms modifies the expressions derived above. Finally, we end by showing the combined effect of all four mechanisms on synapsis efficiency.

##### 2.4.1 Synapsis with stabilized LEFs by BEs

One mechanism that improves synapsis efficiency by increasing *P*_constrained_ is the stabilization of LEFs by BEs. We define, *w*, as the fold increase in LEF lifetime through stabilization by BEs. We assume LEFs with one or both motors bound to a BE have identical fold increase in lifetime, *w*. Intuitively, *P*_constrained_ increases with BE stabilization of LEFs since the LEFs stabilized by BEs extrude larger loops; therefore, DSBs are more likely to happen inside loops. To incorporate the effect of BE stabilization on *P*_constrained_, we modify the rate of BE-LEF dissociation in Eq.(23) to the following:

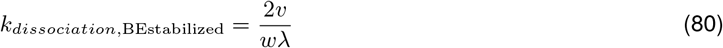

The updated BE-LEF dissociation rate in Eq.(80) in turn updates the fraction of LEF motor subunits extruding in one direction that are bound to BEs in Eqs.(26)-(27) to the following:

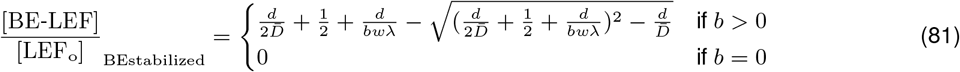

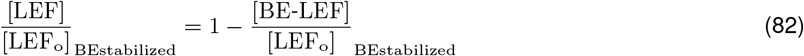

To account for the effect of BE stabilization of LEFs on parameters such as the average LEF processivity and average LEF loop size, we first need to know what fraction of LEFs are stalled at BEs. Let, *β*, be the fraction of LEFs that have at least one motor subunit bound to BEs, then *β* can be calculated as:

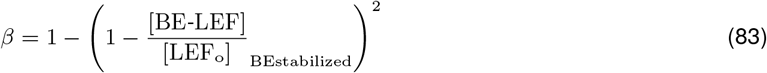

Eq.(83) can reasonably well predict the fraction of LEFs stabilized by BEs (**Supplementary Note 2 Fig. 5**).

**Supplementary Note 2 Figure 5.**
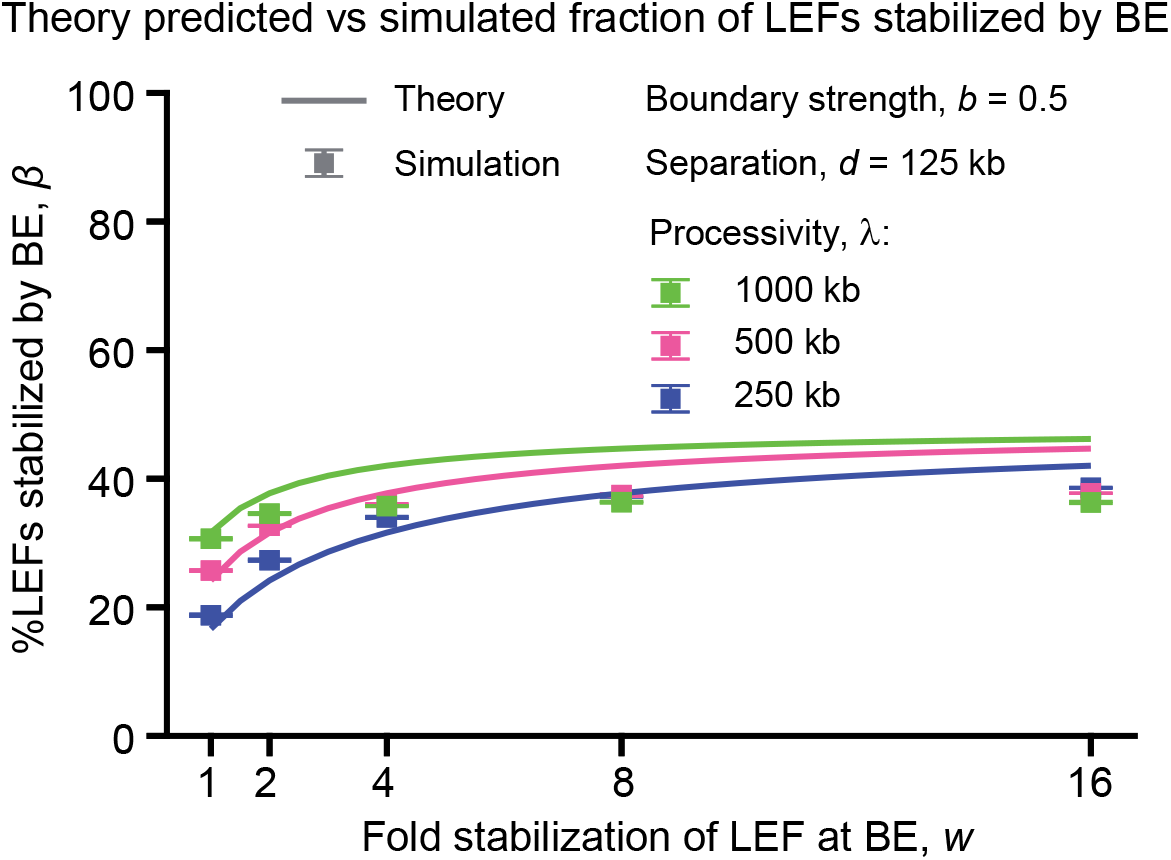
The percentage of LEFs stabilized by BEs predicted by our theory is largely consistent with the simulation results. The error bars represent the standard error of mean, n = 3 independent simulations, with 218 DSB events per simulation.

We can then use Eq.(83) to compute the weighted average processivity and the calculate the average loop size:

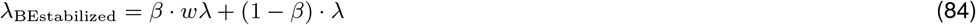

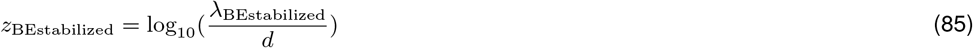

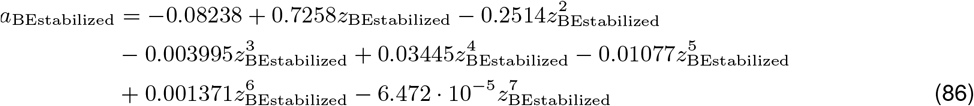

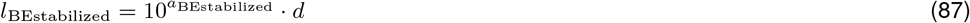

Eqs.(81)-(82) and Eq.(87) in turn update the probability of DSB happening in loops defined in Eq.(17) to the following:

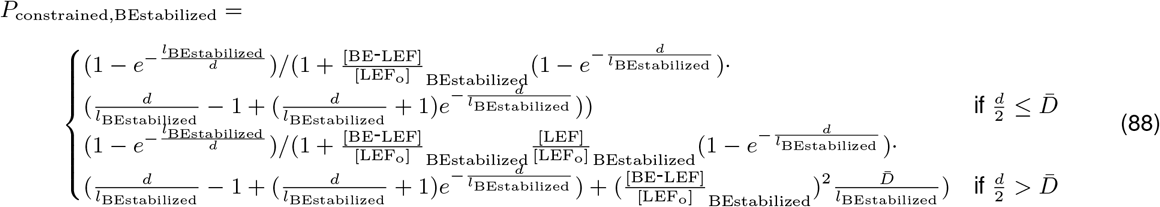

We next aim to compute *P*_end-joining|constrained,BEstabilized_. The first consideration is to account for the effect of BE stabilization on the distribution of constraining LEF lifetimes, *f*_*C*_. We thus sought to calculate the probability that a DSB is flanked by constraining LEFs. As a first-pass, we calculate the probability that a DSB occurs in a TAD held together by at least one constraining LEF. Since a TAD is defined by a pair of convergent BEs, the probability,*P*_stabilized_, of a TAD having at least one BE occupied, is:

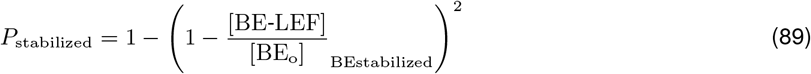

Eq.(89) can accurately predict the fraction of TADs with stabilized LEFs (**Supplementary Note 2 Fig. 6**). As shown in **Supplementary Note 2 Fig. 6**, most of TADs have at least one LEF stabilized by BE in the parameter space considered. Since the LEFs stabilized by BE will likely constrain DSB ends in a DSB-containing TAD due to their prolonged lifetime, we make the simplifying assumption that with LEF stabilization by BE, all constraining LEFs obtain a *w* fold increase in lifetime.

Thus, the modified constraining LEF lifetime distribution is:

**Supplementary Note 2 Figure 6.**
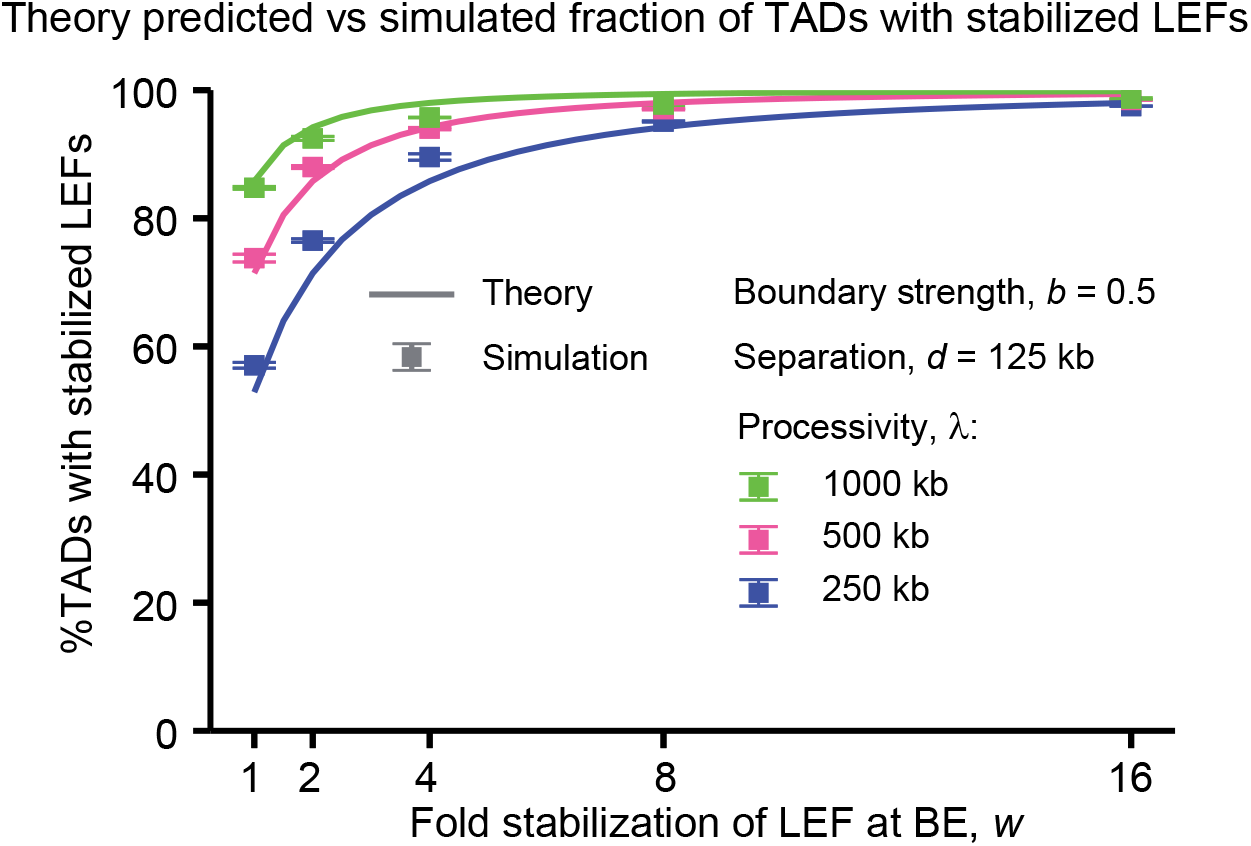
Most TADs have at least one LEF stabilized by BE. The error bars represent the standard error of mean, n = 3 independent simulations, with 218 DSB events per simulation.

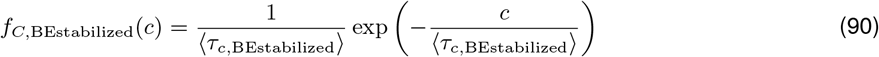

where:

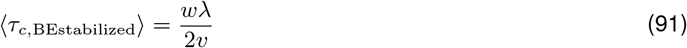

As a further simplifying approximation, we neglect the effect of BE stabilization on loading time distribution, since only the fraction of LEFs stabilized by BEs have slower dynamics.

The modified *f*_*C*_ updates Eqs.(68)-(71) to the following:

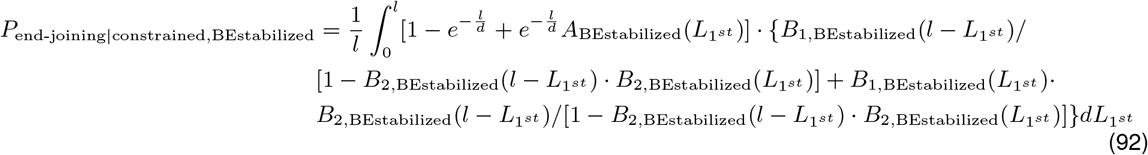

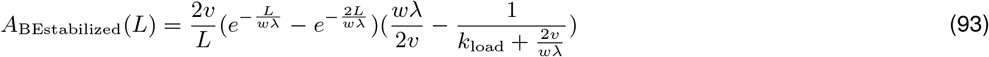

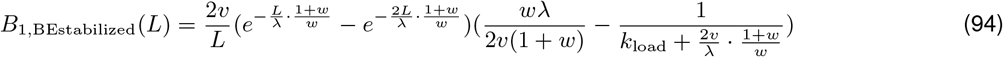

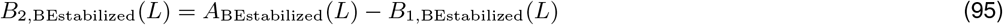

*P*_synapsis,BEstabilized_ can then be computed as:

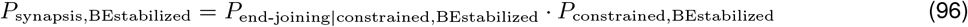

Eq.(96) can accurately predict the synapsis efficiency with BE stabilization, validating our mechanistic explanation of BE stabilization facilitating synapsis by improving the chance of DSB happening inside a loop and increasing the lifetime of constraining LEFs (**Supplementary Note 2 Fig. 7**).

**Supplementary Note 2 Figure 7.**
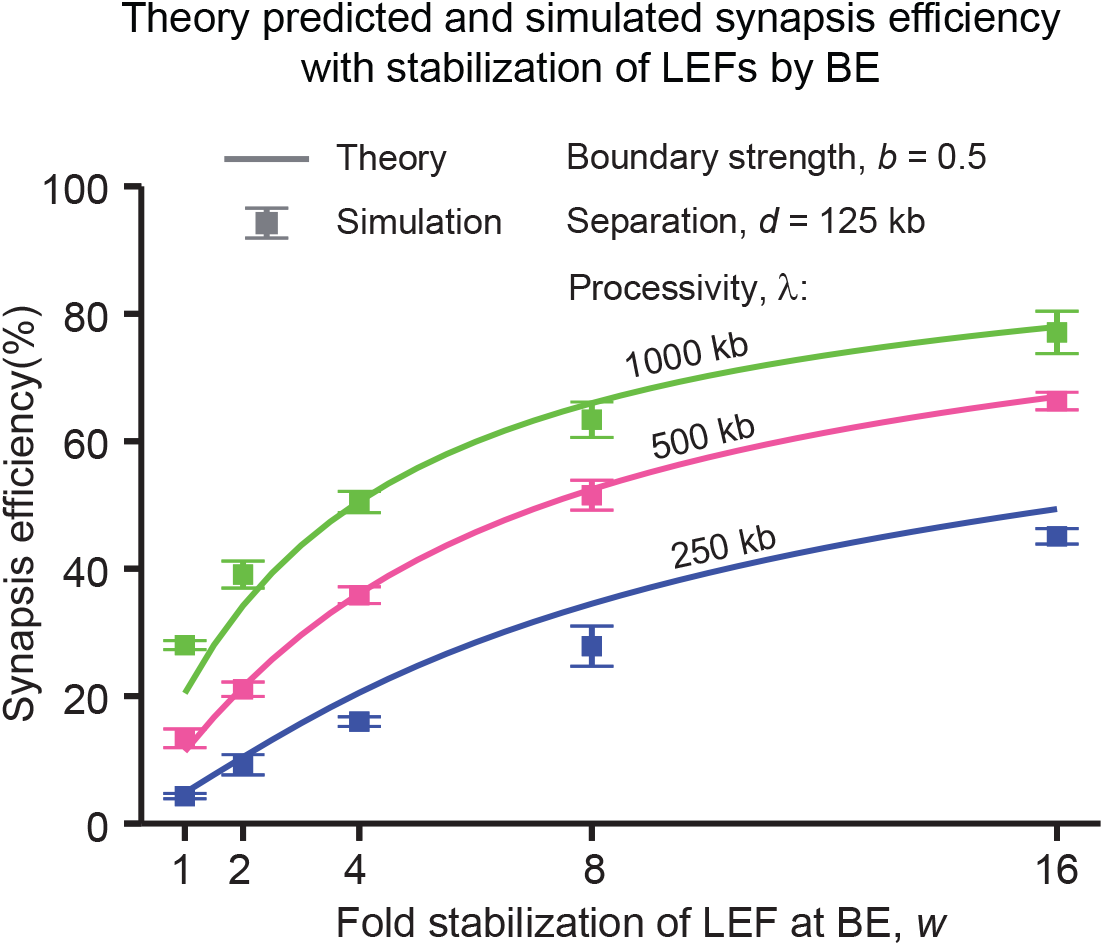
Theory predicts the synapsis efficiency with BE stabilization. Same as the bottom panel in main text **Fig. 4A**. The error bars represent the standard error of mean, n = 3 independent simulations, with 218 DSB events per simulation.

##### 2.4.2 Synapsis with a small fraction of long-lived LEFs

Another mechanism that increases both the chance of DSB happening in loops and *τ*_constrained_ is the presence of a small fraction of long-lived LEFs. Let, *α*_*o*_, be the fraction of long-lived LEFs. Let, *s*, be the fold increase in long-lived LEFs’ lifetime compared with normal LEFs. To incorporate the effect of the presence of a small fraction of long-lived LEFs on *P*_constrained_, we use an weighted average processivity to update the average loop size and the fraction of LEF motors extruding in one direction that are bound to BEs:

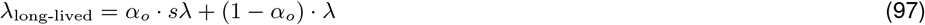

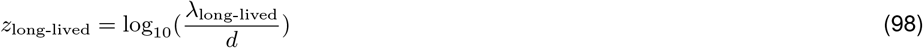

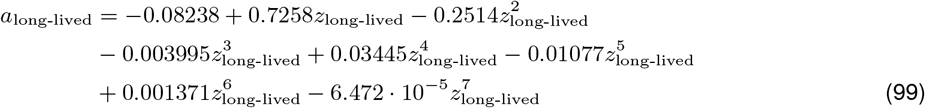

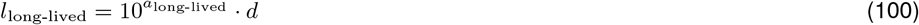

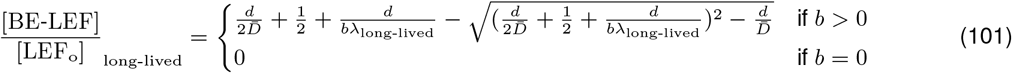

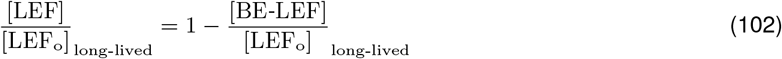

Eqs.(100)-(102) in turn update the probability of a DSB happening in loops defined in Eq.(17) to the following:

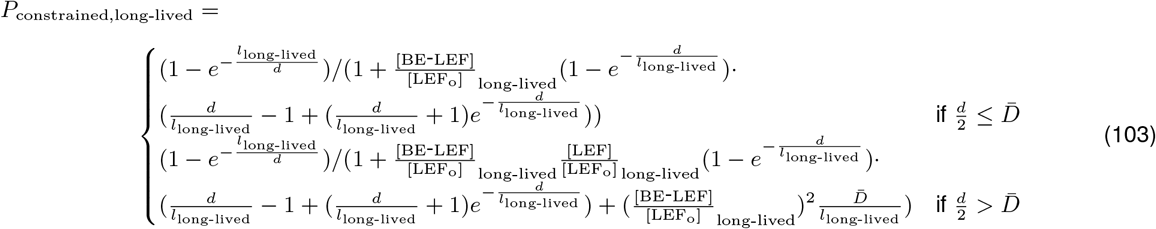

Long-lived LEFs are over-represented in the population of constraining LEFs as they extrude larger loops. We therefore approximate the fraction of constraining LEFs that are long-lived LEFs, *α*, as the fraction of DNA extruded by long-lived LEFs among the DNA extruded by all LEFs assuming all LEFs are unobstructed:

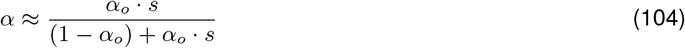

Then PDF of the constraining lifetime will be modified to the following:

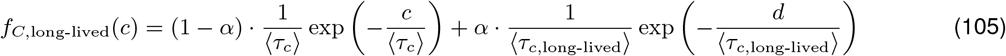

where:

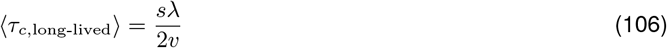

As we only consider a small fraction of long-lived LEFs, we neglect its effect on the loading time distribution as a simplifying approximation.

The modified *f*_*C*_ updates Eqs.(68)-(71) to the following:

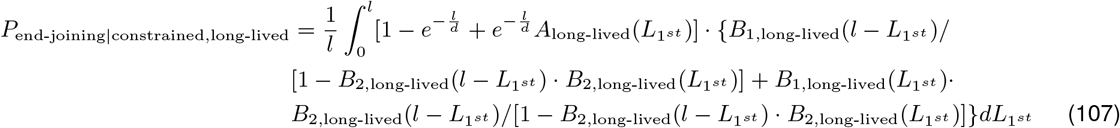

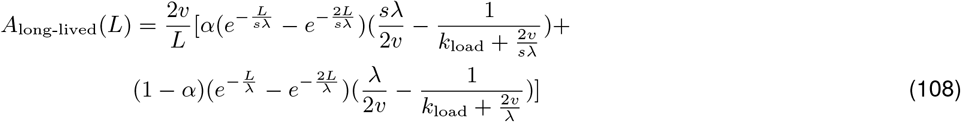

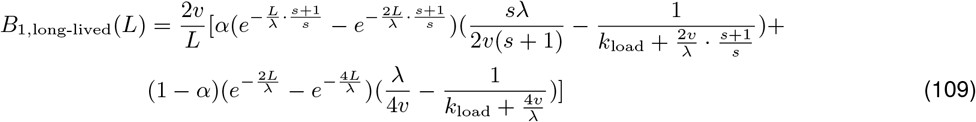

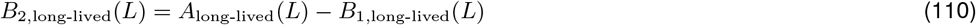

*P*_synapsis,long-lived_ can then be computed as:

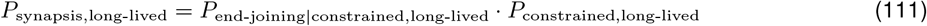

Eq.(111) can accurately predict the synapsis efficiency with a small fraction of long-lived LEFs, validating our mechanistic explanation of a small fraction of long-lived LEFs facilitating synapsis by improving the chance of DSB happening inside a loop and increasing the lifetime of constraining LEFs (**Supplementary Note 2 Fig. 8**).

##### 2.4.3 Synapsis with LEFs stabilized by DSB ends

Mechanisms that stabilize gap-bridging LEFs will increase the likelihood of simultaneous gap-bridging events on both sides of the DSB. One such mechanism is the stabilization of LEFs by DSB ends which affects the lifetime, *G*, of the subpopulation the gap-bridging LEFs that have come into contact with the DSB end. Let, *r*, be the fold increase in *G*. We note that *G* was introduced in Section 2.3.4, to account for the lifetime of gap-bridging LEFs that have already finished gap bridging - therefore, as a first order approximation to simplify our calculations, *r* only affects gap-bridging LEFs that have fully bridge 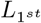, but does not help promote the initial bridging process. The modified PDF of the gap-bridging LEF lifetime can be written as:

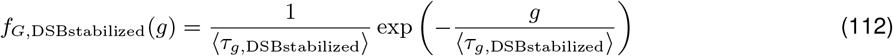

in which:

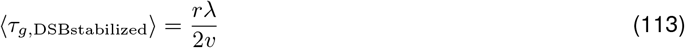

The modified *f*_*G*_ updates Eq.(68) and Eqs.(70)-(71) to the following:

**Supplementary Note 2 Figure 8.**
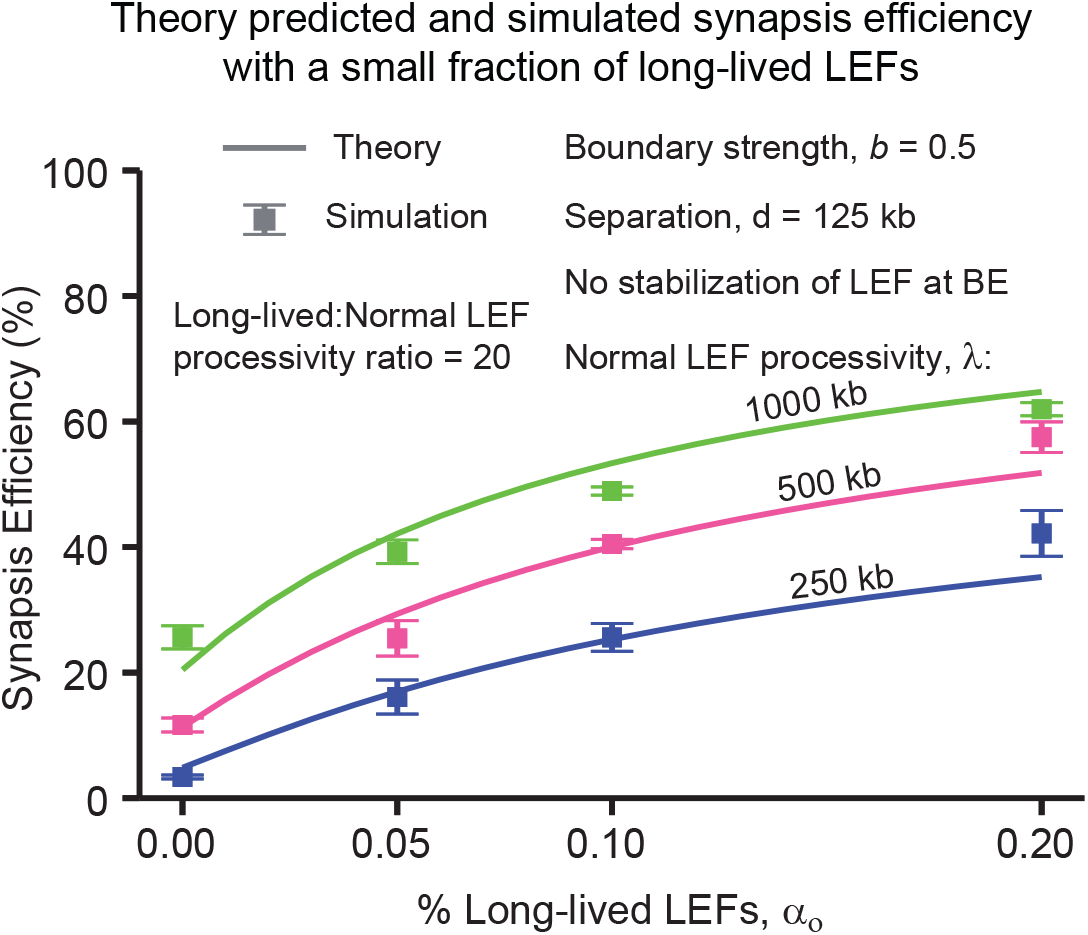
Theory predicts the synapsis efficiency with a small fraction of long-lived LEFs. Same as the bottom panel in main text **Fig. 4B**. The error bars represent the standard error of mean, n = 3 independent simulations, with 218 DSB events per simulation.

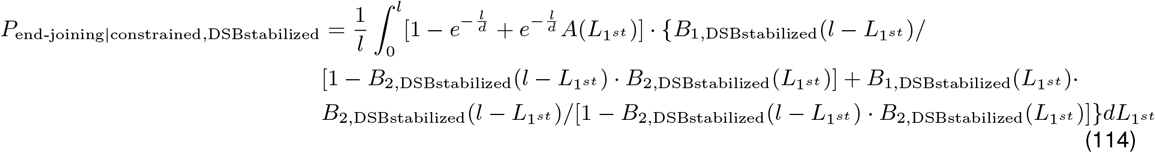

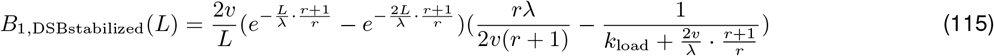

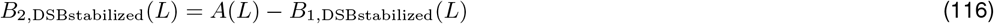

Note that since DSB stabilization is a reactive mechanism that only acts post DSB occurrence, it does not change the probability of DSB occurring in loops. Thus *P*_synapsis,DSBstabilized_ can then be computed as:

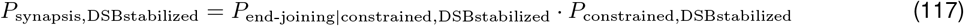

Eq.(117) can accurately predict the synapsis efficiency with DSB stabilization, validating our mechanistic explanation of DSB stabilization facilitating synapsis by increasing chance of simultaneous gap-bridging on both sides of the DSB through prolonged lifetime of gap-bridging LEFs (**Supplementary Note 2 Fig. 9**).

**Supplementary Note 2 Figure 9.**
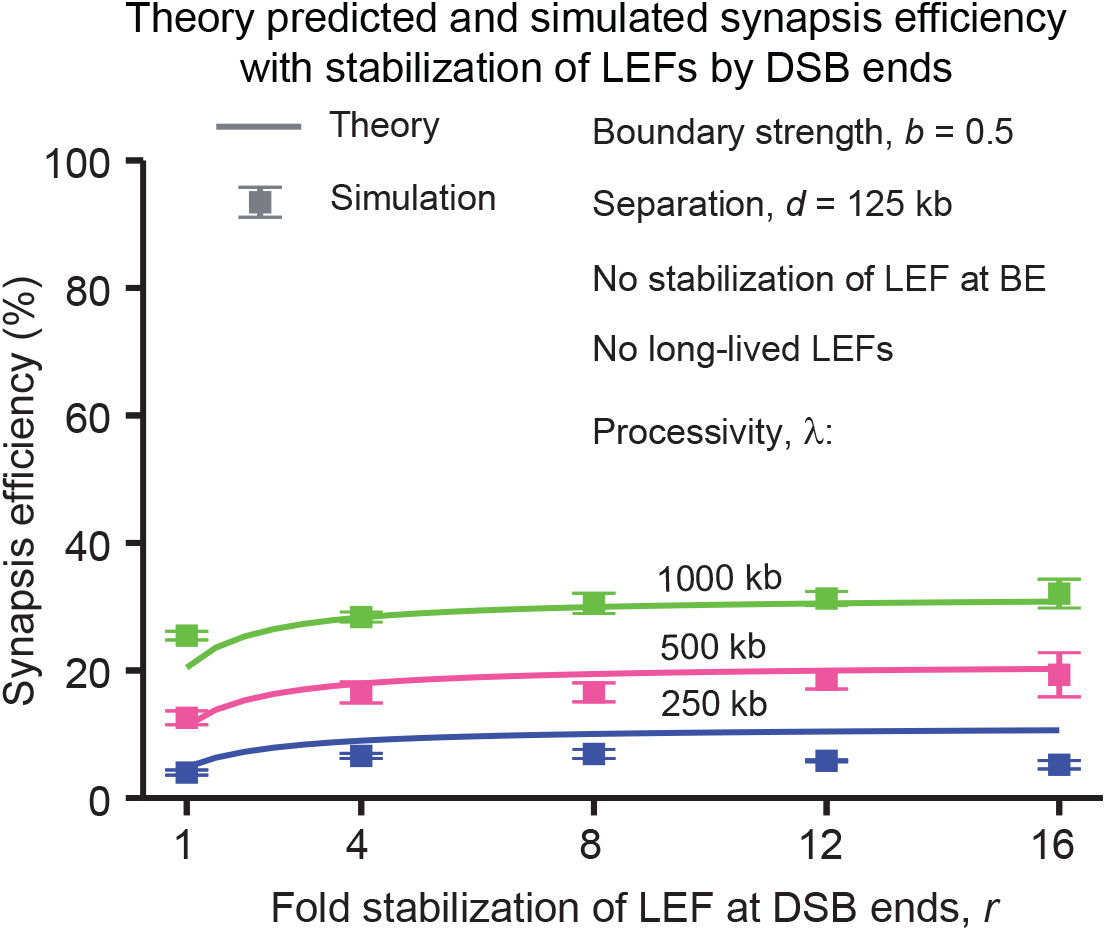
Theory predicts the synapsis efficiency with DSB stabilization. Same as the bottom panel in main text **Fig. 4C**. The error bars represent the standard error of mean, n = 3 independent simulations, with 218 DSB events per simulation.

##### 2.4.4 Synapsis with targeted loading of LEFs to DSB ends

One mechanism that can reduce *τ*_loading_ and thereby increase *P*_end-joining|constrained_ is targeted loading of LEFs to the DSB site. Let, *F*, be the targeted loading factor (i.e., the fold increase in loading probability at the DSB compared with anywhere else in the genome). Let, *U*, be the average distance between two adjacent DSBs in kb, and in all simulations of the paper, we use *U* = 10000 kb, which corresponds to one DSB occurring every 10 Mb. The PDF of the loading time *X* can then be modified as the following:

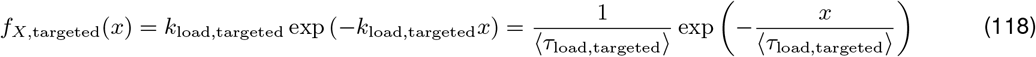

where:

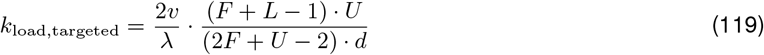

The modified ⟨*τ*_load_⟩ updates Eqs.(68)-(71) to the following:

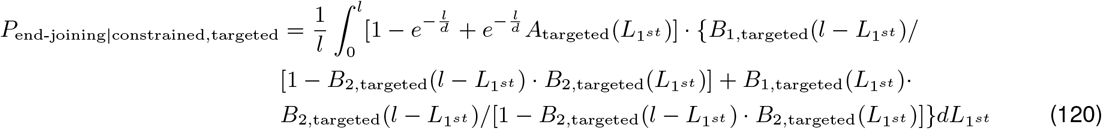

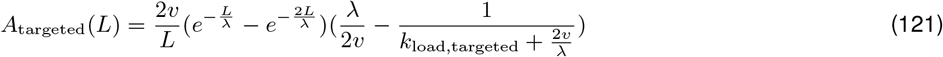

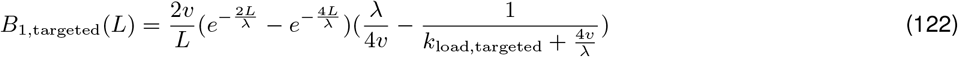

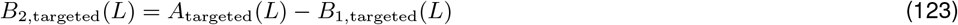

Like DSB stabilization, targeted loading is also a reactive mechanism that only acts post DSB occurrence, it does not change the probability of DSB occurring in loops. Thus *P*_synapsis,targeted_ can then be computed as:

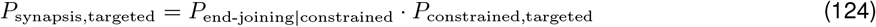

Eq.(124) can accurately predict the synapsis efficiency with targeted loading of LEFs at DSB, validating our mechanistic explanation of targeted loading facilitating synapsis by reducing *τ*_loading_ (**Supplementary Note 2 Fig. 10**)

**Supplementary Note 2 Figure 10.**
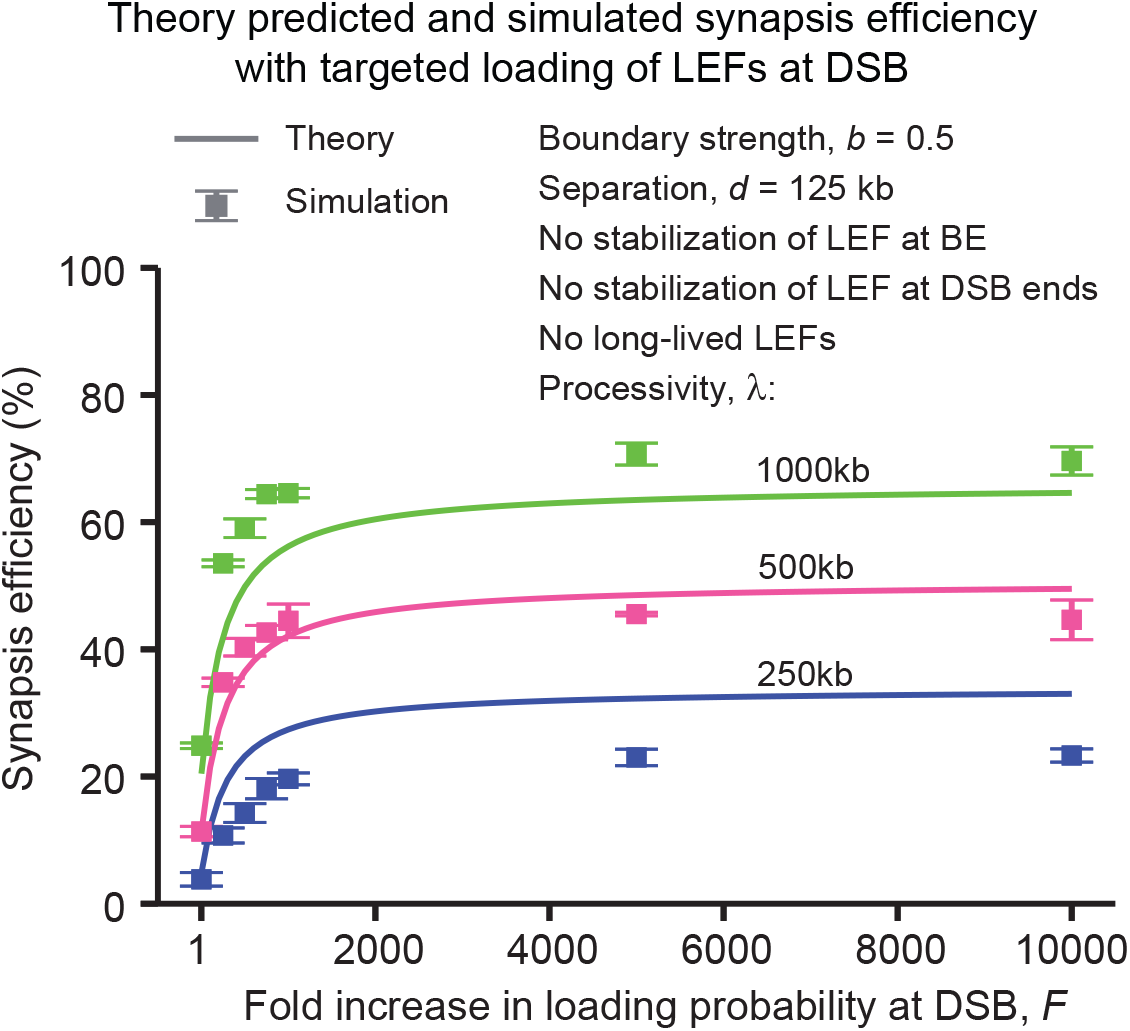
Theory predicts the synapsis efficiency with targeted loading of LEFs at DSB. Same as main text **Fig. 5B**. The error bars represent the standard error of mean, n = 3 independent simulations, with 218 DSB events per simulation.

##### 2.4.5 Synapsis with all four mechanisms combined

Now that we have explored how individual mechanisms facilitate synapsis, we can determine their combined effects. Only BE stabilization and the presence of long-lived LEFs affect *P*_constrained_, as both DSB stabilization and targeted loading are reactive mechanisms. To determine the combined effect of BE stabilization and the presence of long-lived on *P*_constrained_ without further complicating the mathematical form of the solutions and to maximally utilize the framework developed above, we first replace *λ* in Eq.(81) with the weighted average *λ*_long-lived_ defined in Eq.(97) to obtain an updated expression of 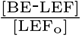 and 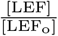:

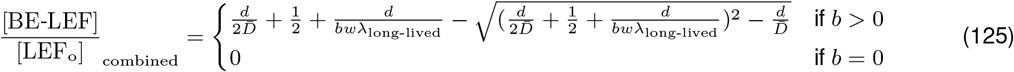

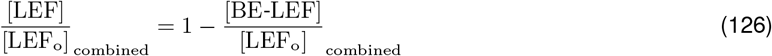

Then the fraction of LEFs stabilized by BEs, *β*, can be updated accordingly:

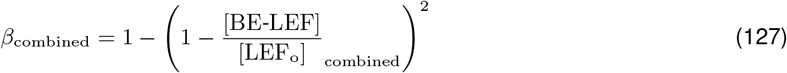

Now we can use the following weighted average processivity to calculate the average loop size, as an approximation of the combined effect of BE stabilization and long-lived LEFs on *P*_constrained_:

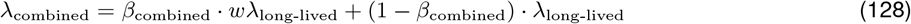

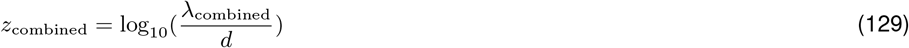

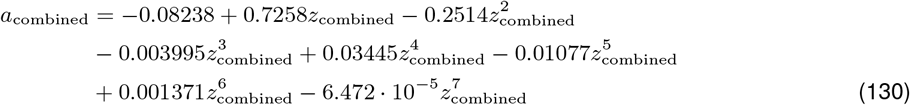

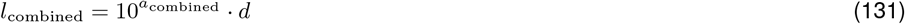

Eqs.(125)-(126) and Eq.(131) in turn update the probability of DSB happening in loops defined in Eq.(17) to the following:

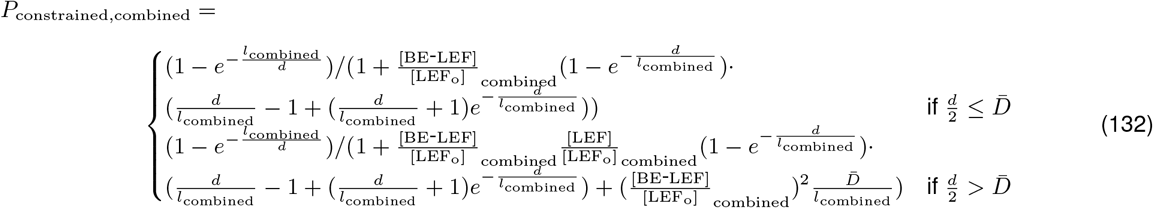

For simplicity, we neglect the effects of BE and DSB stabilization, and long-lived LEFs on the loading time as an approximation:

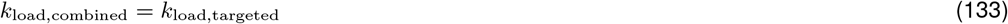

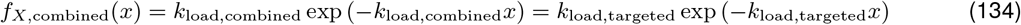

We approximated the combined effect of long-lived LEFs and BE stabilization on the constraining LEF lifetime distribution as the following:

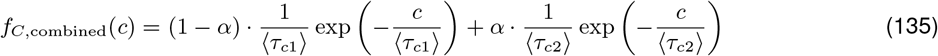

in which:

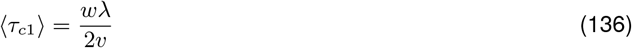

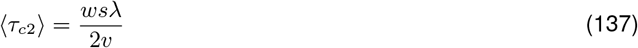

As an approximation, we neglect the scenario of long-lived LEFs acting as gap-bridging LEFs, since long-lived LEFs are much less likely to be loaded at a DSB to function as gap-bridging LEFs given their slow dynamics. Thus the PDF of the gap-bridging LEF lifetime remains the same as Eq.(112):

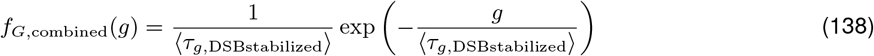

Subsequently, with all four mechanisms combined, Eqs.(68)-(71) are modified to the following:

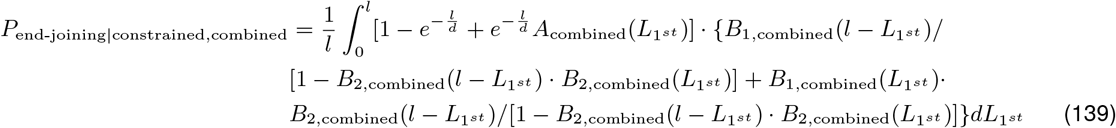

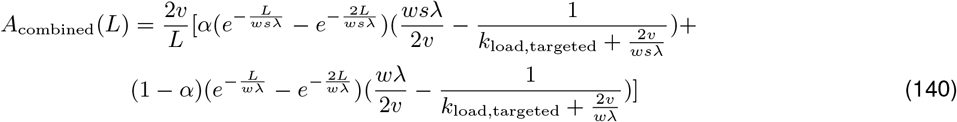

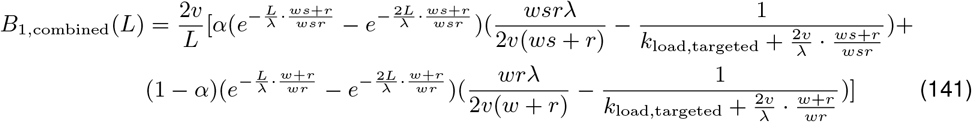

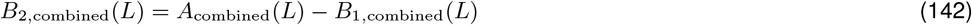

*P*_end-joining|constrained,combined_ can now be calculated by substituting Eqs.(140)-(142) into Eq.(139) and performing numerical integration (see examples in the Mathematica notebooks in **Supplementary Materials**).

*P*_synapsis,combined_ can then be computed as:

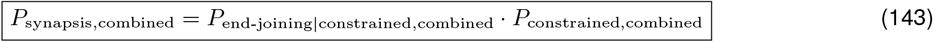

Eq.(143) can predict the synapsis efficiency with all four additional mechanisms combined with reasonable accuracy, supporting our mechanistic explanation of how these mechanisms come together to facilitate DSB end synapsis (**Supplementary Note 2 Fig. 11**).

**Supplementary Note 2 Figure 11.**
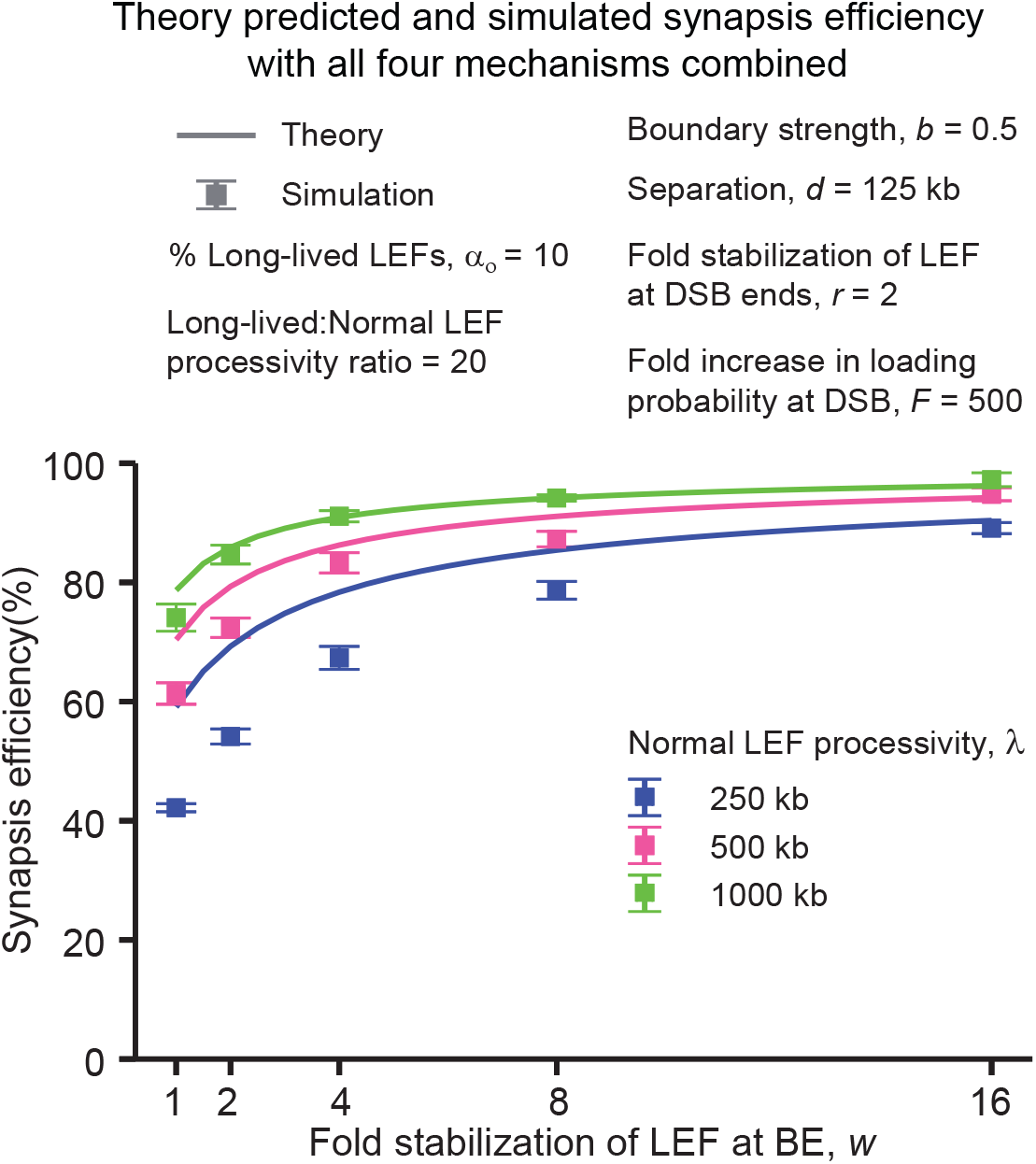
Theory predicts the synapsis efficiency with all four additional mechanisms combined. The error bars represent the standard error of mean, n = 3 independent simulations, with 218 DSB events per simulation.

#### 2.5 Simplified expression for the probability of synapsis with four additional mechanisms

We can derive the simplified expression for the probability of synapsis with all four mechanisms combined.

For *P*_constrain,combined_, we can first apply the same linear approximation (tangent line approximation) of 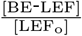 used to obtain Eq.(32):

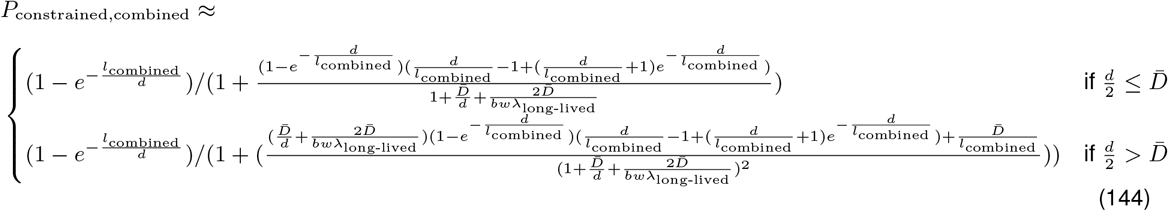

For *P*_end-joining|constrained,combined_, similar to above, we can simplify by considering the scenario where gaps are bridged upon the first try (before the first gap-bridging LEF falls off), with gap length 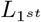 and 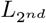 on two sides of the DSB. We assume no gap-bridging LEFs are present within the constraining LEF at the time of DSB occurrence. Then *P*_end-joining|constrained,combined_ can be approximated as the following:

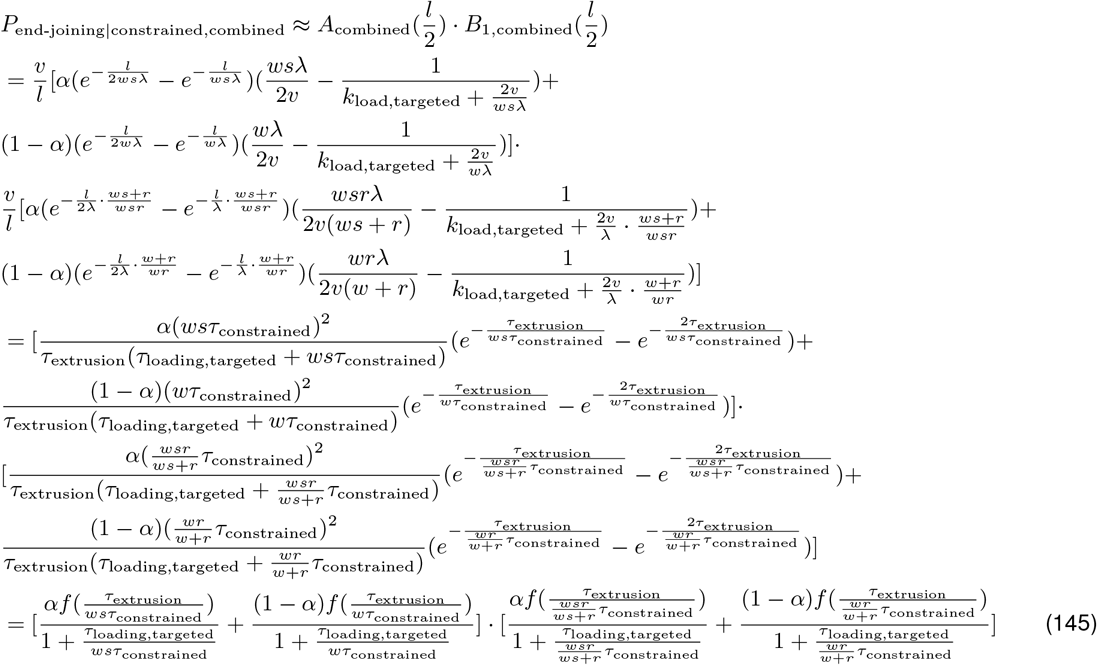

where:

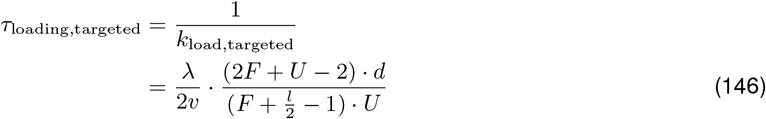

Now we can write the simplified expression for *P*_synapsis, combined_:

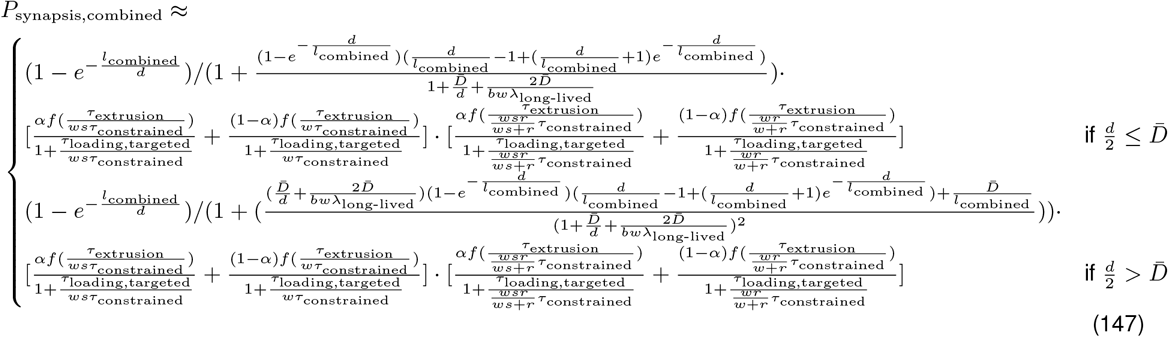

#### 2.6 Two important relative timescales underpinning synapsis efficiency

Since estimates based on experimental evidence suggest that most of the interphase DNA is inside loops at any given time ([15, 17, 18]), *P*_constrained,combined_ is likely close to 1. Thus achieving close-to-perfect synapsis efficiency hinges on *P*_end-joining|constrained,combined_. We next ask what factors underlying *P*_end-joining|constrained,combined_ have the most dominant impact on synapsis efficiency. Notice the recurrent terms, 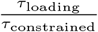 and 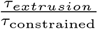, with different prefactors in Eq.(145), we define the following weighted relative timescales:

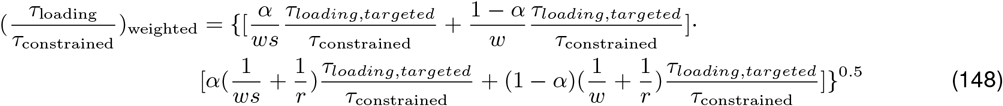

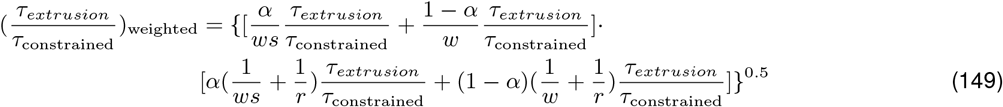

On top of our theoretical framework, we also used 1D simulations to determine whether all four mechanisms could combine to improve synapsis efficiency. We added processivity/separation as an additional dimension, resulting in a 5-dimensional parameter scan with 768 different parameter combinations (see main text **Fig. 6A**). The two weighted relative timescales in Eqs.(148)-(149) can effectively separate the 5-dimensional parameter scan simulation data points based on synapsis efficiency, suggesting lowering the two relative timescales is key to improving synapsis efficiency (**Supplementary Note 2 Fig. 12**).

**Supplementary Note 2 Figure 12.**
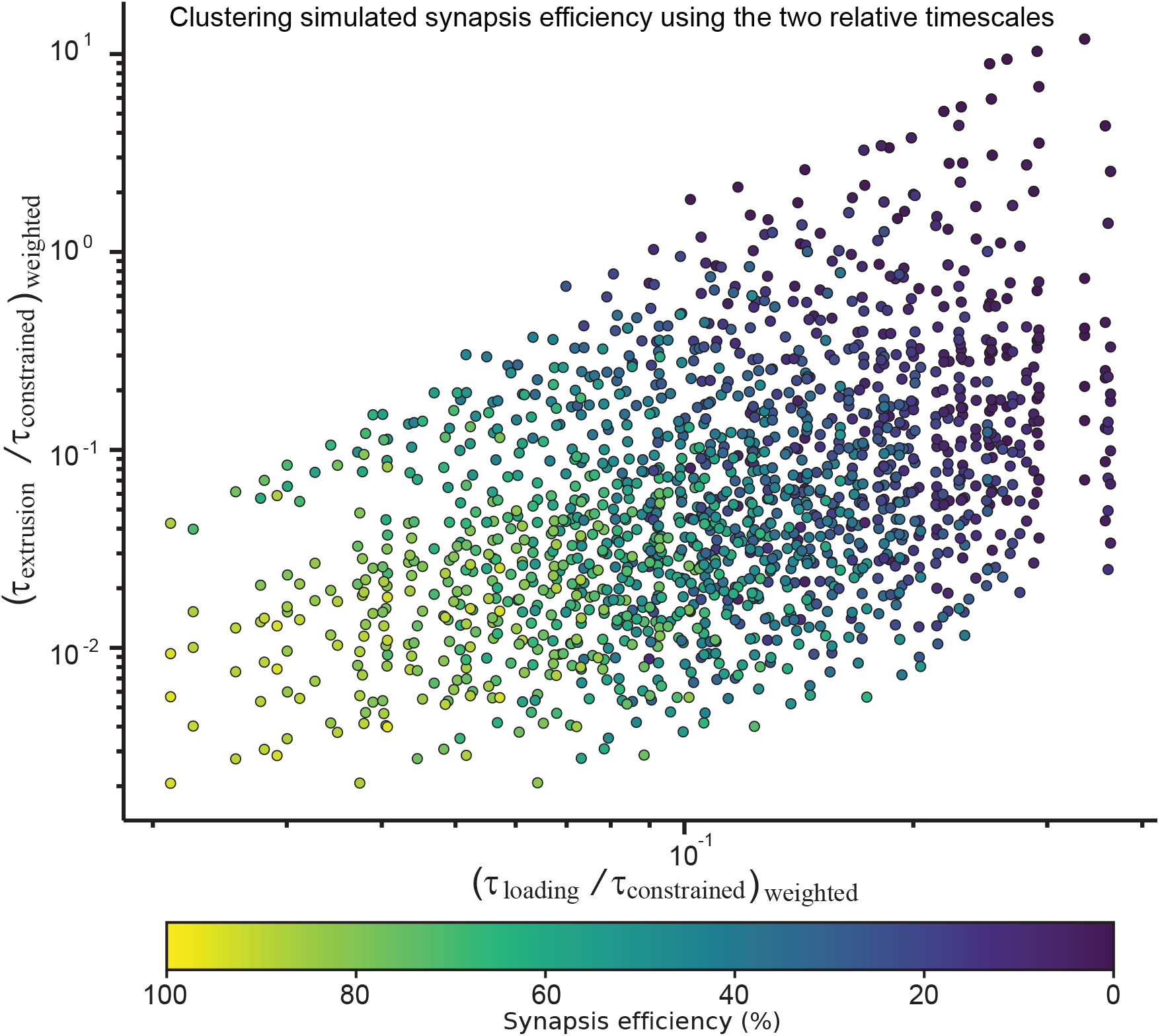
The two relative timescales can effectively cluster simulation data points according to their synapsis efficiency. Same as main text **Fig. 6B**. The color of each data point shows the average synapsis efficiency of n = 3 independent simulations for a given parameter combination, with 218 DSB events per simulation. The two relative timescales for each data point are calculated using Eqs.(148)-(149) based on the input parameters.

#### 2.7 Limitations of our analytical theory

Despite the close agreement between our theory prediction of synapsis efficiency and the simulation results in the parameter space bounded by experimental estimates (see **Supplementary Note 3**), it is worth pointing out several important assumptions and approximations made to simplify the mathematical form of the analytical solution:

1. We assume the prolonged LEF lifetime due to additional mechanisms (BE and DSB stabilization, and long-lived LEFs) does not affect the loading time *Y*. While the accuracy of predictions is largely unaffected by this assumption when *λ > d* as shown above, the assumption no longer holds when *λ* ≤ *d* as the prolonged lifetime for a fraction of LEFs reduces the pool of dynamic LEFs that can be quickly loaded at DSB, leading to overestimation of synapsis efficiency (**Supplementary Note 2 Supplementary Fig. 13**).
2. We assume that only one gap-bridging LEF is bridging the gap on each side of the DSB for the calculation of the first-passage time *T*. This assumption no longer holds when *λ* ≫ *d* and leads to underestimation of synapsis efficiency (top right corner of the heatmaps in **Supplementary Note 2 Fig. 3** where *λ* = 16*d*).
3. We assume if one or more gap-bridging LEFs are present within the constraining LEF at the time of DSB occurrence, one of the two gaps is bridged already. While a seemingly crude approximation to account for the pre-existing gap-bridging LEFs’ impact on synapsis, this assumption does not significantly affect the accuracy of theory prediction.

Perhaps the most significant limitation of our theory is that we neglect the effect of passive 3D diffusion. While we estimate the time it takes for 3D diffusion alone to bring two DSB ends back into proximity is likely too long to be consistent with the synapsis timescale, 3D diffusion likely acts concurrently and synergistically with loop extrusion to further improve synapsis efficiency. When the two DSB ends are brought close enough by loop extrusion, the chance of the two DSB ends randomly encountering each other through passive diffusion also greatly increases. Therefore, the synapsis efficiency predicted by our theory likely delineates the lower bound of the physiological synapsis efficiency.

Finally, we have also made several biological assumptions that abstract the complex synapsis process into a simplified picture more tractable for theoretical analysis:

1. We assume uniform loading probability of LEFs across the genome aside from DSB ends. While it remains unclear to what extent cohesins show preferential loading to certain genomic regions, previous work suggests that cohesin loading may not be uniform across the genome [19–21].
2. We assume uniform LEF extrusion speed throughout the genome and that only BEs may stall LEF extrusion, while it has been shown that other non-BE elements such as RNA polymerases could also act as partial extrusion barriers [19, 22, 23].
3. We assume a uniform probability of DSB formation throughout the genome, while evidence suggests certain sites such as binding sites for CTCF and cohesin are more prone to generate DSBs [24–28]. Since DSB sites close to CTCF likely benefit more from BE stabilization to improve synapsis efficiency, and DSB sites close to cohesin binding sites are more likely to be constrained by LEFs and have gap-bridging LEFs loaded in the gaps to improve synapsis efficiency, our theory might underestimate the physiological synapsis efficiency by assuming a uniform probability of DSB formation.

**Supplementary Note 2 Figure 13.**
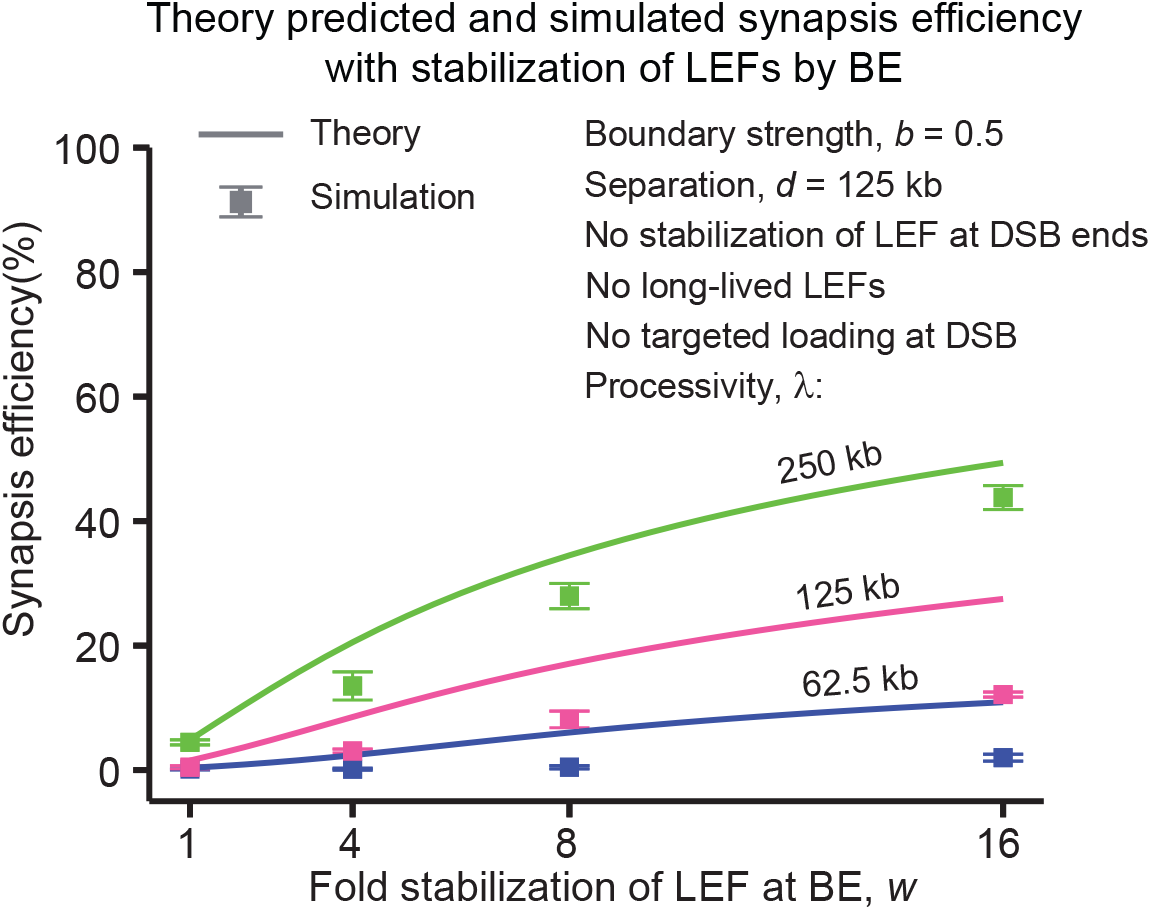
Theory overestimates the synapsis efficiency when *λ ≤ d*. The error bars represent the standard error of mean, n = 3 independent simulations, with 218 DSB events per simulation.

#### 2.8 Conclusion

In conclusion, we built a probabilistic theoretical model with a relatively simple mathematical form, that can predict the simulated synapsis efficiency in the parameter regime with *λ > d* with a fairly good accuracy. Our theory highlights two important roles of loop extrusion in synapsis: LEFs can constrain DSB ends and thereby prevent them from diffusing apart, and gap-bridging LEFs can mediate synapsis by bringing the DSB ends back together. Since DSB ends can only be joined by loop extrusion if the DSB occurs inside a loop, loop coverage of the genome is crucial to achieve high synapsis efficiency. Mechanisms that stabilize LEFs before the DSB occurs such as BE stabilization and the presence of a small fraction of long-lived LEFs could serve to improve the coverage of the genome by LEFs. Provided that the DSB ends are constrained, our theory then points to two relative timescales, 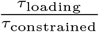 and 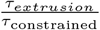that underlie synapsis efficiency. By extending our simple theory with 4 physiologically plausible mechanisms, we show how each of the four mechanisms decreases the two relative timescales and thereby improves synapsis efficiency. The final expressions with all four mechanisms combined illustrate the synergistic effects of the mechanisms. Despite the various approximations made in deriving the theory, the relatively good agreement between our analytical theory and simulations lends credibility to our mechanistic insights obtained from our theoretical model.

Our theory can also serve to guide experimental perturbation of the DSB synapsis machinery for further validation the role of different extended mechanisms discussed above, and to explain observations of the perturbations’ impact on synapsis efficiency. Our theory has direct implications for the dependence of DSB repair efficiency on genomic context. For example, our theory points to the importance of BE stabilization in efficient synapsis, which predicts reduced synapsis efficiency in genomic region that lacks BEs (CTCF binding sites e tc.). Indeed, heterochromatin, which usually has lower density of bound CTCF binding sites, has been shown to be more sensitive to radiation-induced chromosomal aberrations than euchromatin in Chinese hamster cells [29], consistent with our theory prediction.

### 3 Supplementary Note 3

#### Overview

Given the large possible parameter space for individual variables considered in our simulations and modeling as well as the vast parameter combinations generated by permutation, we sought to use prior experimental estimates to generate plausible upper and lower bounds on individual parameter values. In this Supplementary Note, we detail the calculations used to justify the parameter bounds chosen in our study.

#### 3.1 Estimation of the range of LEF separation and processivity

Two studies performed absolute quantification of CTCF and cohesin in HeLa cells [18] and mouse embryonic stem cells (mESCs) [17]. Since in simulations we assume LEFs re-load somewhere on the genome immediately after unloading, we could use the density of chromatin-bound cohesin to estimate the LEF separation, *d* (the inverse of LEF density).

Using fluorescence-correlation spectroscopy (FCS) and fluorescence recovery after photobleaching (FRAP), the chromatin-bound SCC1(subunit of cohesin) copy number has been estimated to be ∼ 160, 000 for HeLa cells [18]. Considering the total HeLa genome length of 7.9 Gb [18], we can estimate the cohesin spearation as the following, assuming cohesin exists as monomeric ring [30]:

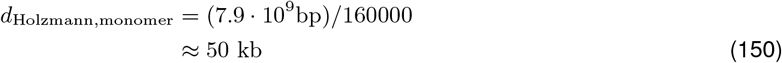

If we assume cohesin extrudes as dimers, then [30]:

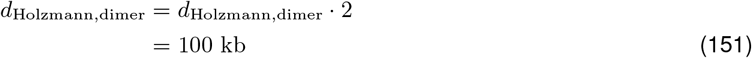

Through a combination of FCS, “in-gel” fluorescence, and flow cytometry, we previously carried out an absolute quantification of CTCF and cohesin in mESCs. We estimated the cohesin density to be 5.3 per Mb (assuming monomeric) or 2.7 per Mb (assuming dimeric) [17, 30], which correspond to the following cohesin separation:

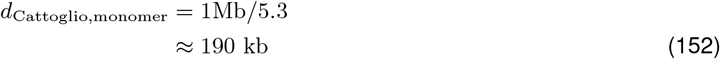

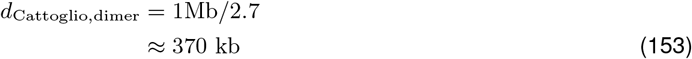

Thus far, absolute quantification of cohesin has only been performed in two mammalian cell types, HeLa and mESC. As can be seen, the density varies substantially between these two cell types, and may vary even more among other cell types. It is therefore associated with significant uncertainty. Nevertheless, these two studies bound the LEF separations in the following range:

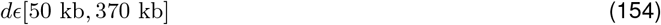

Let, *τ*_*o*_, be the LEF lifetime (residence time). Let *v*_*T*_, be the total unobstructed LEF extrusion speed. Then for the two-sided LEFs considered in our study, *v*_*T*_ = 2*v*, where *v* is the unobstructed extrusion speed in one direction. Then LEF processivity can be computed as the product of total extrusion speed (in both directions) and the LEF lifetime:

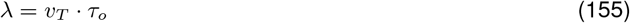

Several single-molecule experiments for condensins and cohesins [31–35] measured the unobstructed total extrusion speed to be in the range of *v*_*T*_ *∈* [0.5 kb/s, 2 kb/s]. Note that the extrusion speed estimated from *in vivo* measurement is often slower [30, 36, 37], likely due to various protein roadblocks bound to DNA including BEs like CTCF. While synapsis time calculated from our simulations is inversely proportional to extrusion speed, synapsis efficiency is independent of extrusion speed. Unless otherwise specified, we use *v*_*T*_ of 1 kb/s for the calculation of synapsis time.

Multiple studies have estimated cohesin’s residence time which varies with cell cycle phase. Focusing on G1, we estimated cohesin’s residence time to be ∼ 22 minutes in mESCs [38], whereas Holzmann *et al*. estimated it to be ∼ 13.7 minutes in HeLa cells [18]. In contrast, the residence time of condensin I was determined by FRAP to be ∼ 2 minutes [39]. Thus the bounds of *v*_*T*_ and *τ*_*o*_ estimated from these studies provide the following range of LEF processivity given Eq.(155):

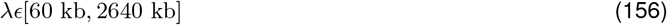

Given the estimated bounds of LEF separation and processivity in Eq.(154) and Eq.(156), to keep the processivity/separation ratio as geometric integer sequence, we use the separation list *d* = [62.5, 125, 250, 500] kb and the processivity list *λ* = [62.5, 125, 250, 500, 1000] kb in the simulations without additional mechanisms. To limit the parameter combinations in the the 5-dimensional parameter scan simulations, we used the separation and processivity combinations (*d, λ*) = [(125, 62.5), (125, 125), (125, 250), (250, 125), (250, 250), (250, 500)] kb, so that processivity/separation ratios of 0.5,1 and 2 are examined.

#### 3.2 Estimation of the fold increase in LEF lifetime upon stabilization by BEs

CTCF can increase cohesin’s chromatin residence time by interacting with cohesin in such a way that it outcompetes cohesin interactions with WAPL, which unloads cohesin from DNA [40]. An independent study found that CTCF could also facilitate cohesin acetylation and prolong the lifetime of acetylated cohesin [41]. Cohesin contains one of the two variant STAG subunits, STAG1 or STAG2 [42]. CTCF stabilizes both cohesin-STAG1 and cohesin-STAG2, but with a lesser extent for cohesin-STAG2 [41]. Cohesin-STAG1’s stable residence time, *τ*_cohesin-STAG1,stable_, and dynamic residence time, *τ*_cohesin-STAG1,dynamic_, during G1 phase have been determined by inverse fluorescence recovery after photobleaching (iFRAP) [41]:

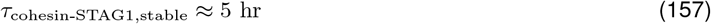

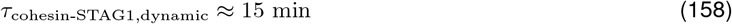

Since the longer lifetime of cohesin-STAG1 could be attributed to both stabilization by CTCF and acetylation of SMC3 (a subunit of cohesin) by ESCO1 and that cohesin-STAG2 is stabilized by CTCF to a lesser extent than cohesin-STAG1 [41], the ratio of the stable residence time of cohesin-STAG1 and the dynamic residence time of cohesin-STAG1 could serve as an upper bound for the fold increase in LEF lifetime due to stabilization by BEs, *w*:

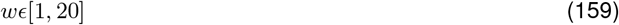

#### 3.3 Estimation of the fraction and lifetime of long-lived LEFs

Long-lived LEFs can be conceptualized as a subpopulations of LEFs with intrinsic longer lifetime due to chemical modifications such as acetylation of the cohesin subunit SMC3. The STAG subunit of cohesin can be composed of either STAG1 or STAG2 given rise to two distinct forms of cohesin. Cohesin-STAG1 is preferentially acetylated during G1 phase, and cohesin-STAG2 contains four times less acetylated SMC3 relative to cohesin-STAG1 [41]. About ∼ 30 − 50% of cohesin-STAG1 is stably bound to chromatin in G1 [38, 41, 43]. The relative levels of STAG1 and STAG2 varies substantially between cell types [44]. Cohesin-STAG1 constitutes about about 25% of total cohesin in HeLa cell [18], about 33% of total cohesins in immortalized mouse embryonic fibroblasts (iMEFs) [45], and about 55% in normal human bronchial epithelial cells (NHBE) [46], with the rest being cohesin-STAG2.

Taken together, if we assume all the stably bound cohesin-STAG1 is acetylated, up to ∼ 30% of all cohesin is then acetylated, which we use as an upper bound for the fraction of long-lived LEFs, *α*_*o*_, since factors other than chemical modifications such as stabilization by CTCF could also contribute to the stably bound cohesin’s longer lifetime [41]:

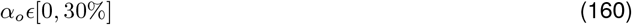

Stably bound SCC1 exhibits about 50 fold increase in lifetime compared with the dynamic fraction of SCC1 [41]. Thus we can bound the fold increase in long-lived LEFs’ lifetime compared with normal LEFs:

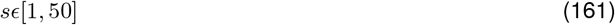

To limit the number of free variables, we use an intermediate value *s* = 20 throughout our study.

#### 3.4 Estimation of the fold increase in LEF lifetime upon stabilization by DSB ends

The ATM kinase at DSB ends is hypothesized to phosphorylate cohesin, thereby increasing cohesin’s lifetime [47]. When cohesin is stabilized by ATM kinase at DSB ends, the higher processivity means that there is higher probability that cohesin can extrude all the way to adjacent BEs. About 1.5 fold enrichment of SCC1, a subunit of cohesin, at BEs was observed in DSB-containing TADs relative to SCC1 count prior to DSB occurrence [47]. We performed simulations with different fold stabilization of LEF at DSB ends, and found the ∼ 1.5 fold enrichment of SCC1 at BEs in DSB-containing TADs corresponds to about 2 to 4 fold stabilization of LEF at DSB ends (**Supplementary Note 3 Fig. 1**). Given the uncertainty associated with the experimental measurement, we use 8 as an upper bound for the fold increase in LEF lifetime due to stabilization by DSB ends, *r*:

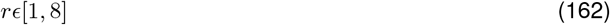

**Supplementary Note 3 Figure 1.**
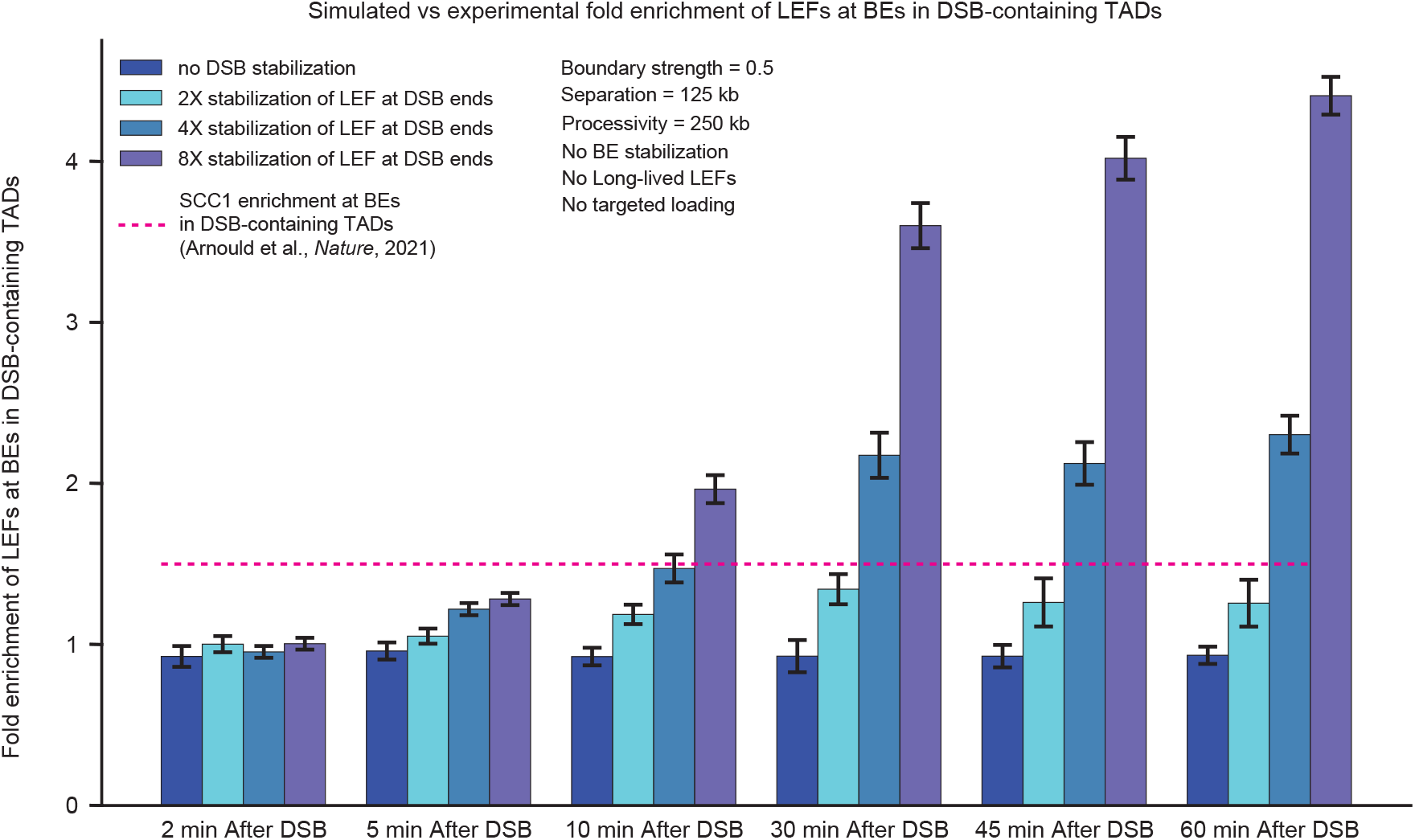
Simulated LEF enrichment at BEs in DSB-containing TADs is consistent with experimental observations. Fold enrichment is calculated as the number of LEFs in the DSB-containing TADs within 5 kb to the BEs at the indicated time points normalized against the number of LEFs in the same regions prior to DSB occurrence. The error bars represent standard error of mean, n = 216 DSB-containing TADs. The pink dashed line represents the SCC1 fold enrichment at BEs in the DSB-containing TAD determined by Arnould et al. [47].

#### 3.5 Estimation of the fold increase in LEF loading probability at DSB

Enrichment of cohesins at DSBs has been reported in several studies [47–50], and this enrichment was recently found to be dependent on cohesin loader NIPBL, as well as ATM and MRN complex recruited to DSB sites [47], pointing to a reactive mechanism that targets cohesin to DSB sites. We performed simulations with different fold increase in LEF loading probability at DSB, and found the ∼ 1.57 fold enrichment of SCC1 in the chromosome 20 DSB-containing TAD of Dlva cells [47] corresponds to about 250 fold increase in LEF loading probability at DSB (**Supplementary Note 3 Fig. 2**). We implemented targeted loading by increasing the loading rate of LEFs within 1 kb of DSB for simplicity, whereas the observed accumulation of LEFs is not limited to the immediate proximity to DSBs but the whole DSB-containing TADs [47], suggesting LEFs might be targeted to larger regions around DSB instead of just the DSB ends. Therefore, we compared the fold enrichment of LEFs in DSB-containing TADs here instead of just comparing the fold enrichment immediately around DSB ends. Given the noise in ChIP-seq experiments and the 2.5-5 times higher fold enrichment of SCC1 within 4kb around DSB reported by Cheblal et al. [50], we use 5000 (corresponding to 7 fold enrichment of LEFs in DSB-containing TADs) as an upper bound for the fold increase in LEF loading probability at DSB, *F*:

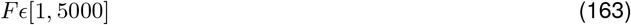

**Supplementary Note 3 Figure 2.**
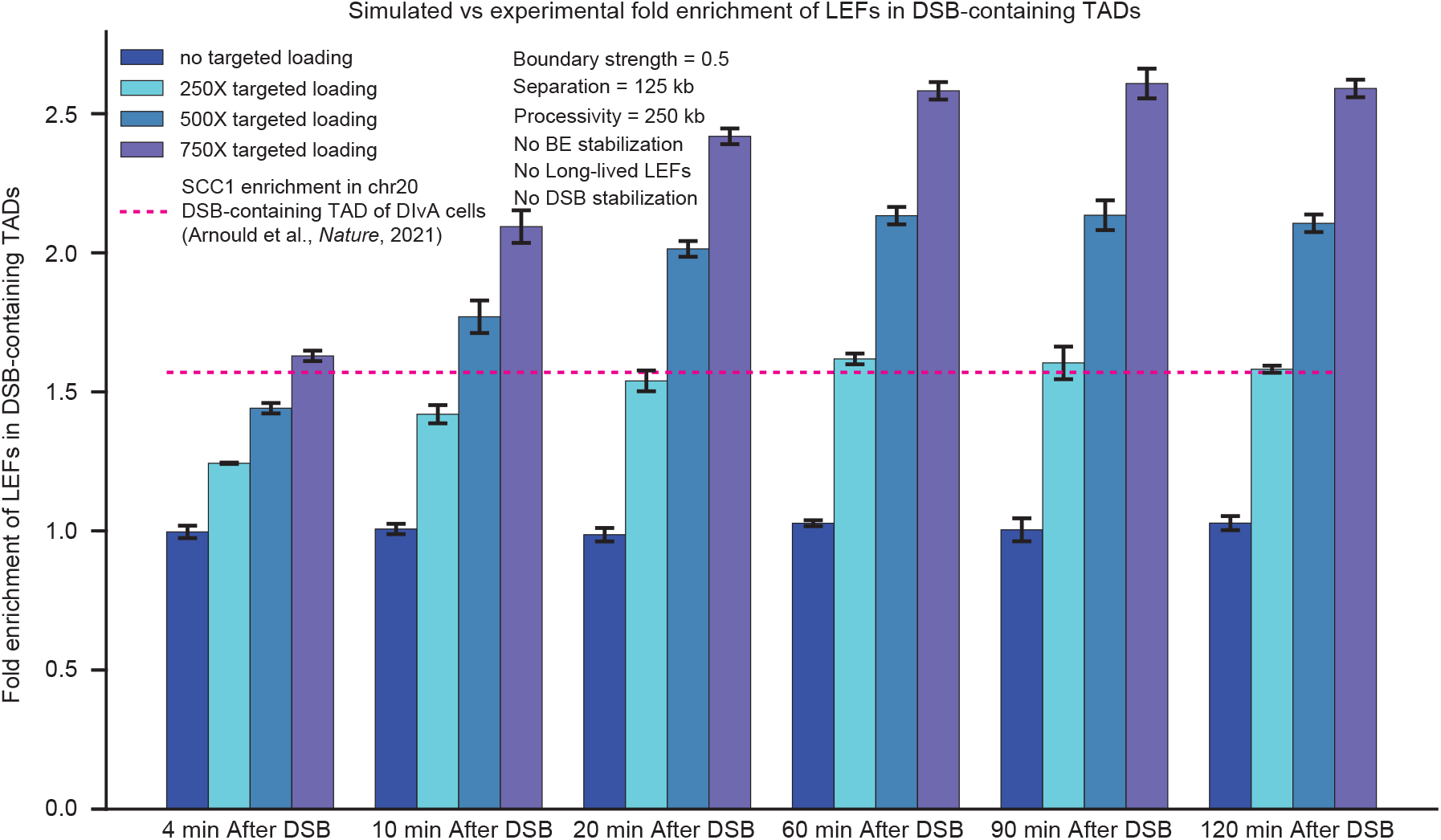
Simulated LEF enrichment in DSB-containing TADs is consistent with experimental observations. Same as **Supplementary Fig. 5A**. Fold enrichment is calculated as the number of LEFs in the DSB-containing TADs at the indicated time points normalized against the number of LEFs in DSB-containing TADs prior to DSB occurrence. The error bars represent standard error of mean, n = 216 DSB-containing TADs. The pink dashed line represents the ChIP-seq SCC1 enrichment in the DSB-containing TAD (TAD boundaries determined by CTCF ChIP-seq data) on chromosome 20 of DIvA cells determined by Arnould et al. [47].

### 4 Supplementary Figures

**Supplementary Figure 1.**
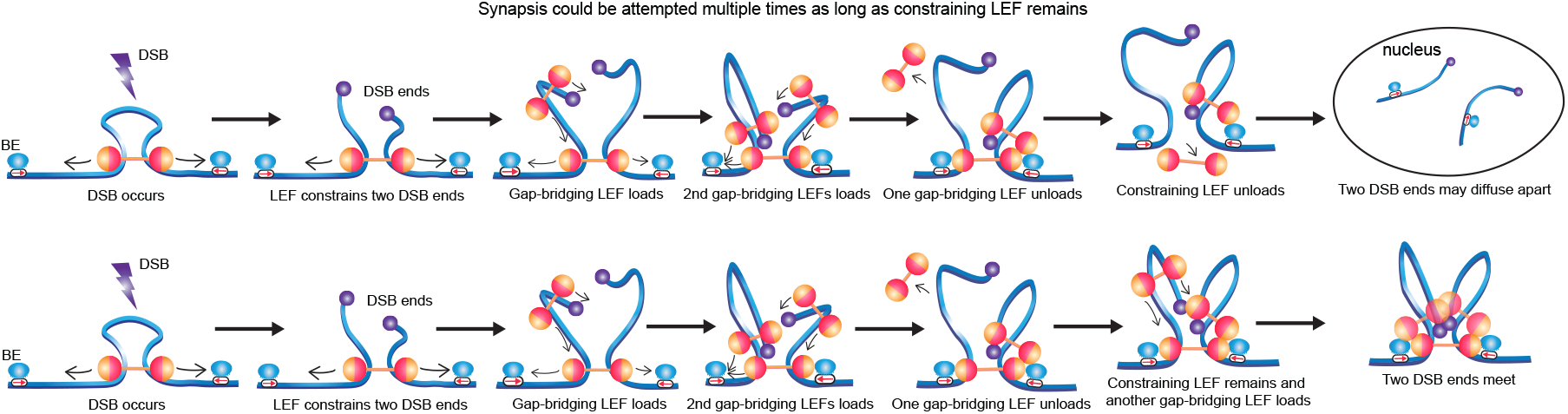
Synapsis could be attempted multiple times as long as constraining LEF remains. Top row: we assume no further synapsis could be attempted once constraining LEF unloads as two DSB ends may diffuse apart. Bottom row: as long as constraining LEF remains, if gap-bridging LEFs unload before synapsis is achieved, additional gap-bridging LEFs could be loaded to attempt synapsis for multiple rounds until successful synapsis is achieved or the constraining LEF unloads.

**Supplementary Figure 2.**
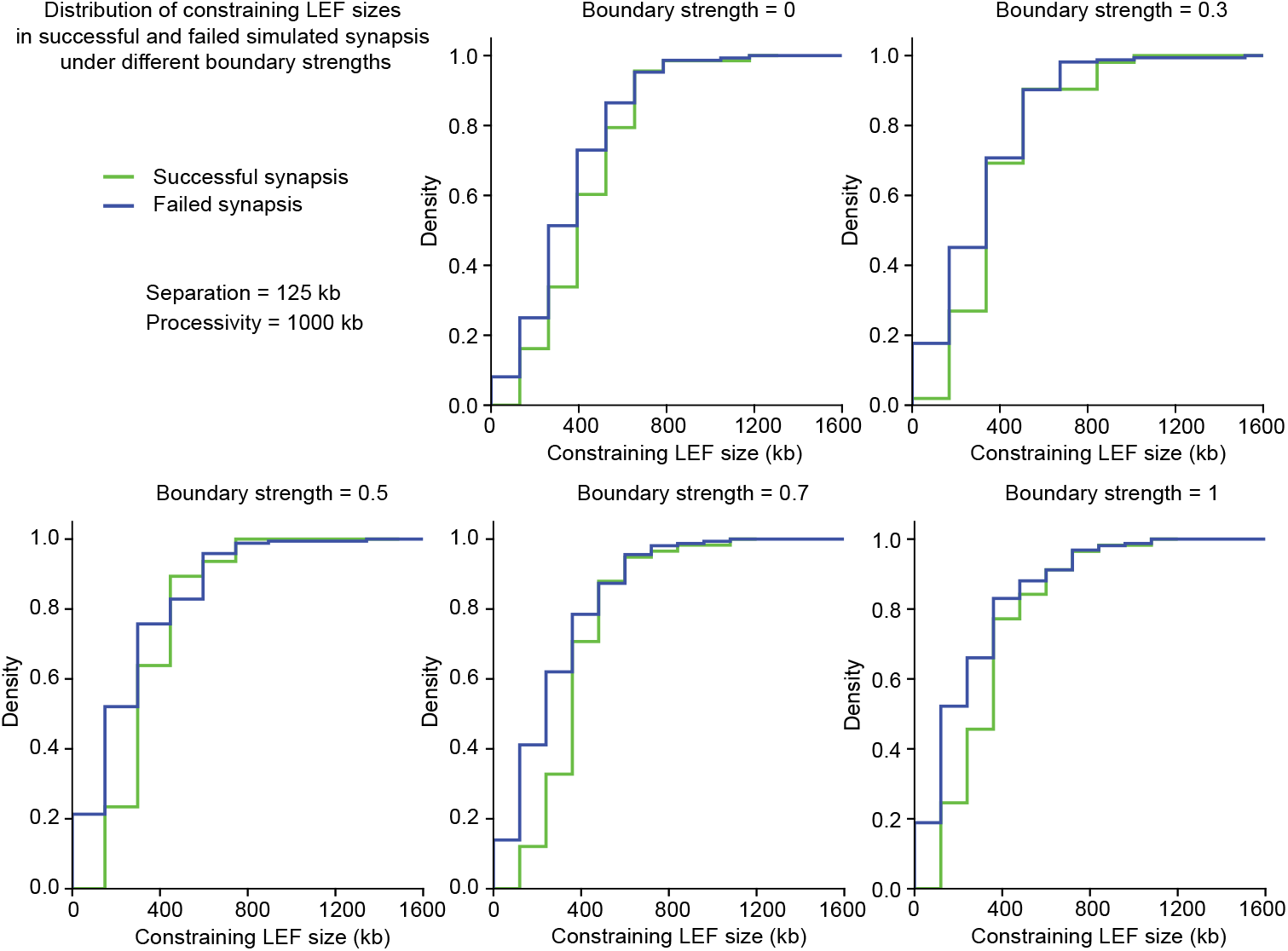
Constraining LEF size does not correlate with boundary strength or whether synapsis succeeds or fails. The cumulative density function (CDF) of constraining LEF size in successful and failed simulated synapsis. Processivity = 250 kb, separation = 250 kb. Constraining LEF size was recorded at the moment when synapsis is achieved or at the moment when constraining LEF unloads for successful and failed synapsis events respectively.

**Supplementary Figure 3.**
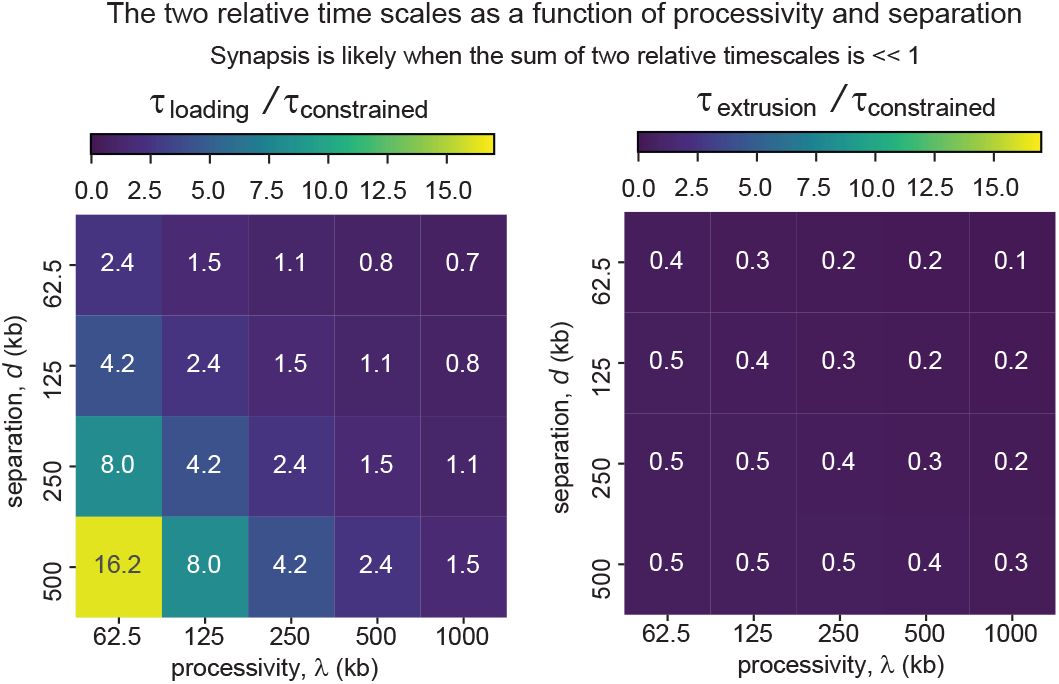
Loading of gap-bridging LEF is the rate limiting step of synapsis. Heatmaps of *τ*_loading_/*τ*_constrained_ (left) and *τ*_extrusion_/*τ*_constrained_ (right) calculated from Eqs.(74)-(76) across different separation and processivity combinations.

**Supplementary Figure 4.**
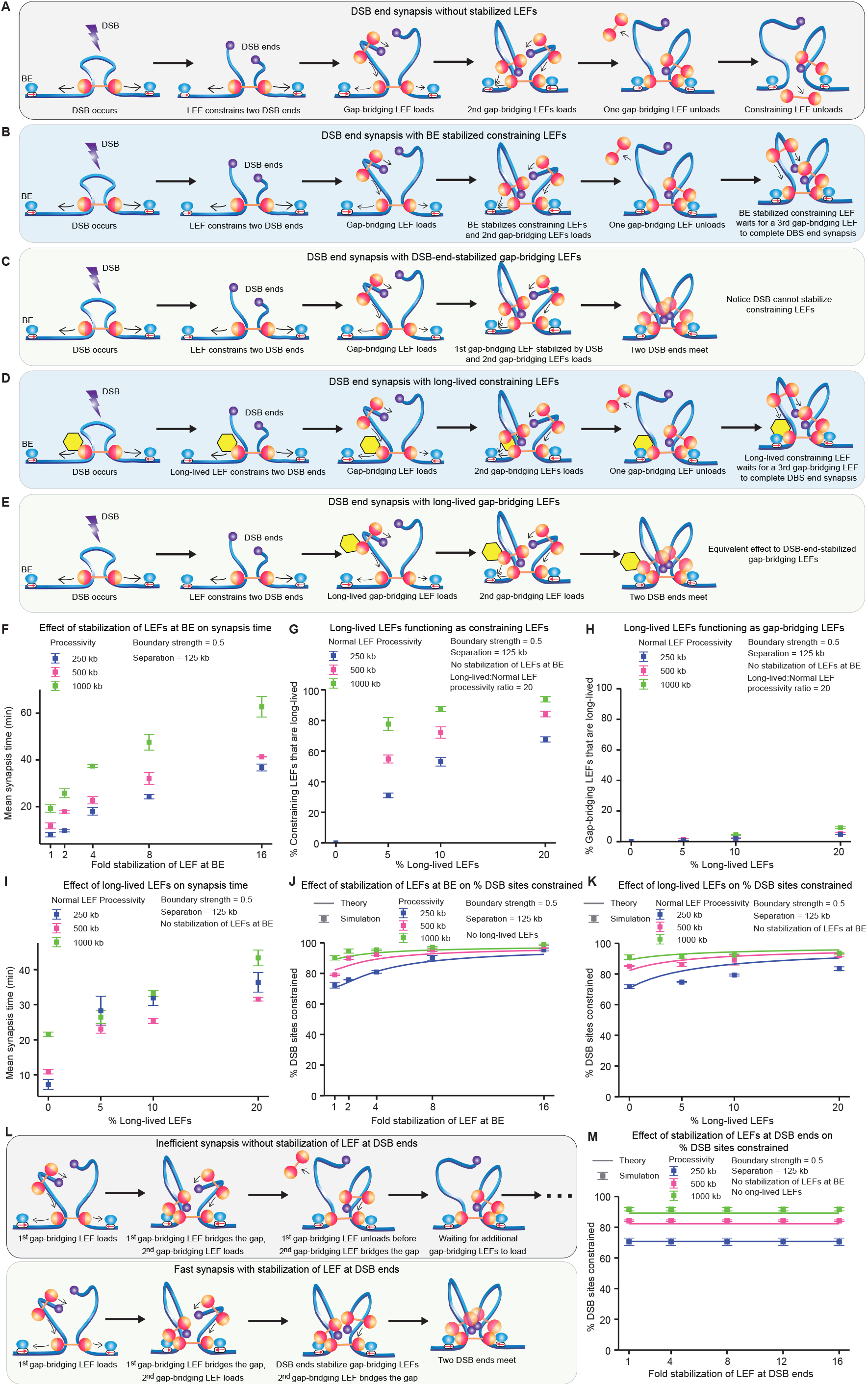
Mechanisms of synapsis facilitated by stabilization of LEFs by BE and DSB, and a small portion of long-lived LEFs. (**A**-**E**) Schematic diagrams of synapsis with LEFs without stabilization (A), with constraining LEFs stabilized by BE (B), with gap-bridging LEFs stabilized by DSB ends (C), with long-lived constraining LEFs (D), or with long-lived gap-bridging LEFs (E). (**F**) The effect of stabilization of LEF at BE on the mean synapsis time. Conversion of simulation time steps to synapsis time assumes total extrusion speed of 1 kb/s. The error bars represent the standard error of mean, n = 3 independent simulations, with 216-218 DSB events per simulation. (**G**) Percentages of constraining LEFs that are long-lived. The error bars represent the standard error of mean, n = 3 independent simulations, with 216-218 DSB events per simulation. (**H**) Percentages of gap-bridging LEFs that are long-lived. The error bars represent the standard error of mean, n = 3 independent simulations, with 216-218 DSB events per simulation. (**I**) The effect of stabilization of LEF at BE on the mean synapsis time. Conversion of simulation time steps to synapsis time assumes total extrusion speed of 1 kb/s. The error bars represent the standard error of mean, n = 3 independent simulations, with 216-218 DSB events per simulation. (**J,K**) The effect of stabilization of LEF at BE (J) and a small subpopulation of long-lived LEFs (K) on the probability of DSB occurring inside a DNA loop. The error bars represent the standard error of mean, n = 3 independent simulations, with 216-218 DSB events per simulation. (**L**) Schematic diagram of synapsis with LEF stabilization by DSB ends, leading to more efficient synapsis. (**M**) DSB stabilization has no impact on the probability of DSB occurring inside a DNA loop. The error bars represent the standard error of mean, n = 3 independent simulations, with 216-218 DSB events per simulation.

**Supplementary Figure 5.**
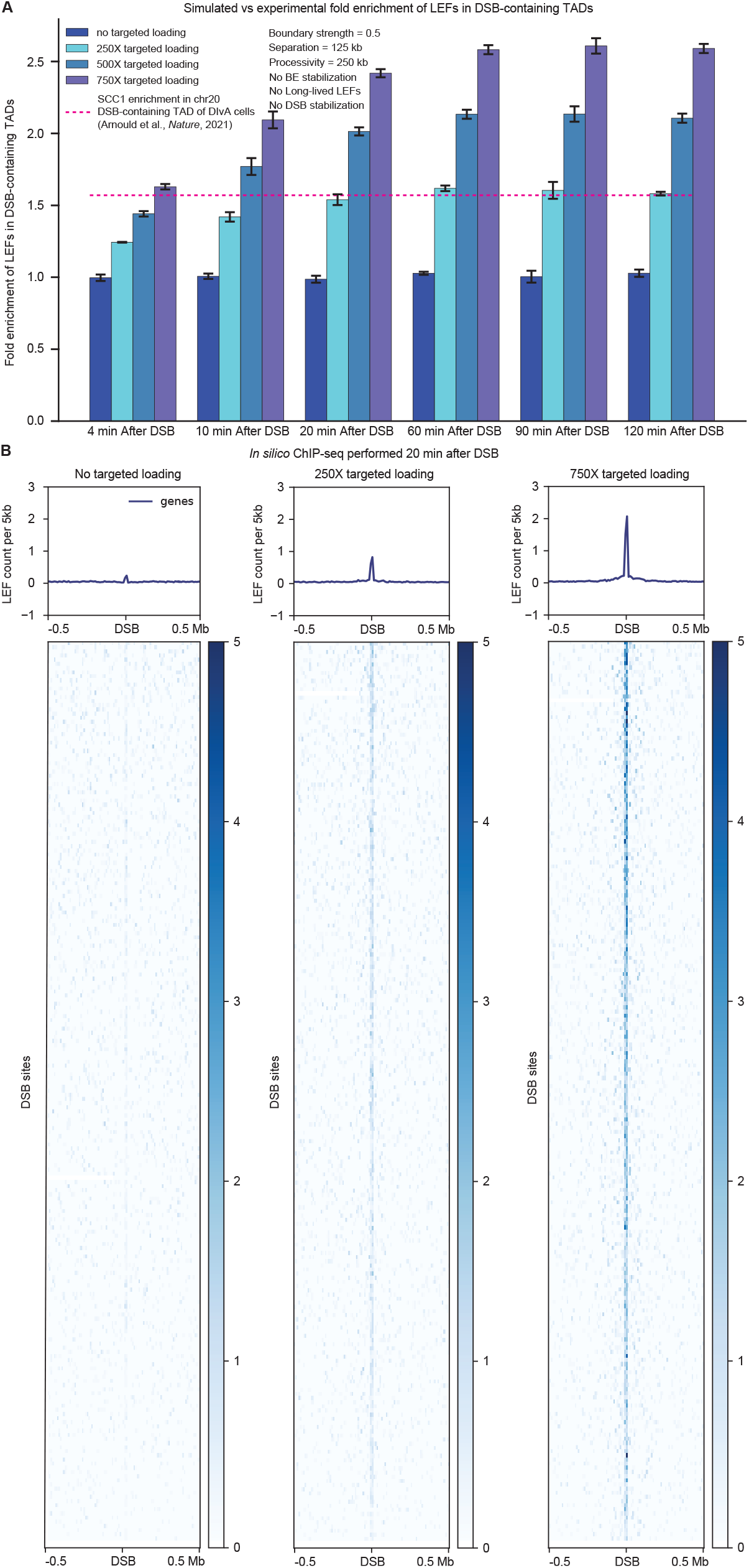
*In silico* ChIP-seq results showing LEF enrichment at DSB consistent with experimental observations. (**A**) Fold enrichment of LEFs in DSB-containing TADs. Fold enrichment is calculated as the number of LEFs in the DSB-containing TADs at the indicated time points normalized against the number of LEFs in DSB-containing TADs prior to DSB occurrence. The error bars represent standard error of mean, n = 216 DSB-containing TADs. The pink dashed line represents the ChIP-seq SCC1 enrichment in the DSB-containing TAD (TAD boundaries determined by CTCF ChIP-seq data) on chromosome 20 of DIvA cells [47]. (**B**) Enrichment heatmaps and metaplots of LEFs at DSB sites 20 minutes post DSB occurrence, aligned by their distance from the DSB site along the genome (Mb). Bin size=5 kb.

**Supplementary Figure 6.**
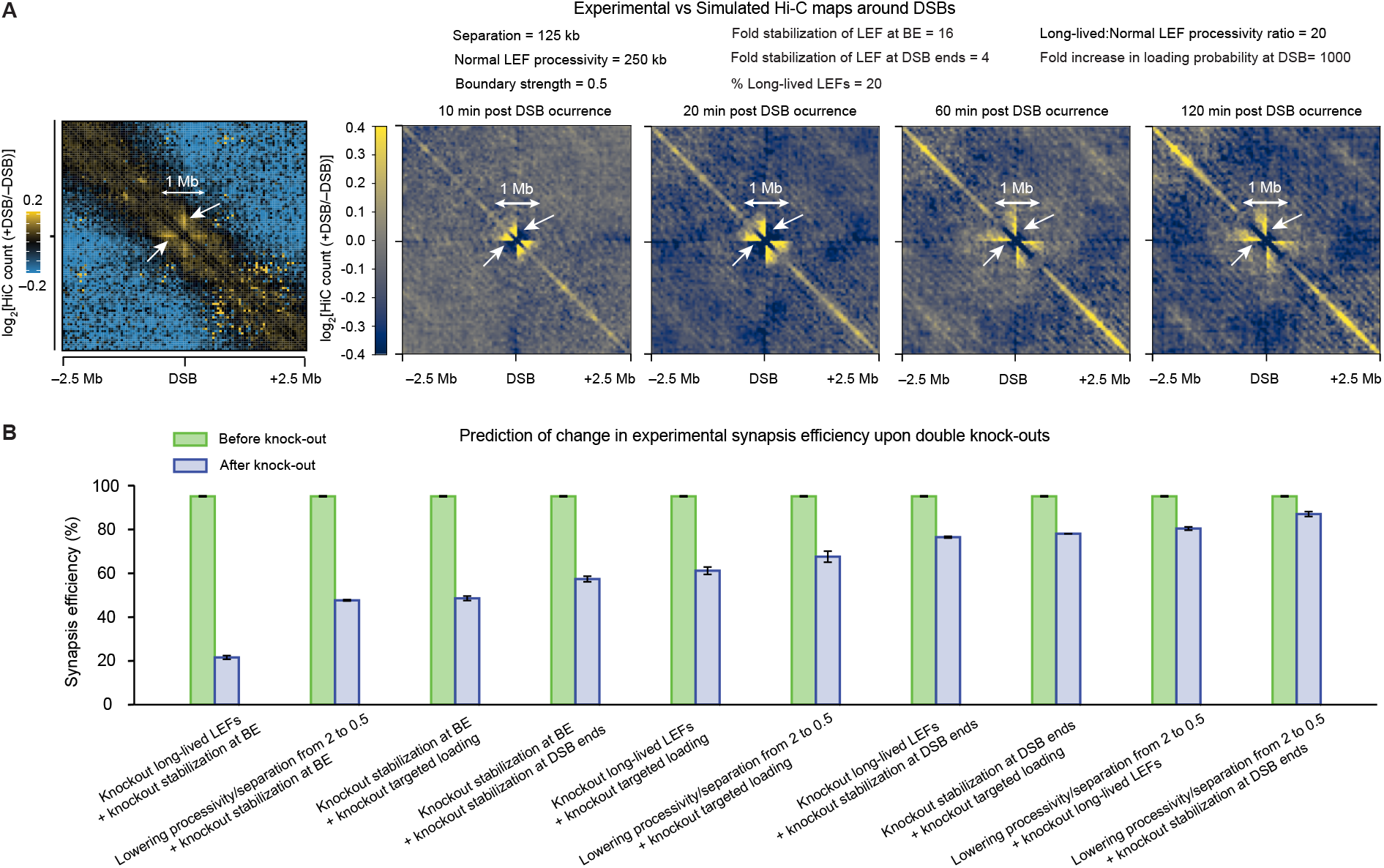
Parameter combination with *≥* 95% synapsis efficiency predicts Hi-C maps with similar pattern as experimental data, and double knock-outs significantly reduces synapsis efficiency for most knock-out combinations. (**A**) The Hi-C contact maps of log_2_[+DSB/-DSB] centered on the DSBs at the indicated time points post DSB occurrence averaged over 603 DSBs (50-kb resolution, 5-Mb window) show a similar pattern to the experimental Hi-C map reused from Fig.2b by Arnould et al. [47] shown in the left. The contact maps are simulated with one parameter combination (indicated in legend) producing *≥* 95% synapsis efficiency. The white arrows highlight the stripe pattern. (**B**) The bar plot shows the synapsis efficiency before and after knocking out two of the five mechanisms discussed in **Fig. 6**. The error bars represent standard error of mean, n = 3 different parameter combinations that achieved *≥* 95% synapsis efficiency before knock-out.

## References

[1] Ross, W. E. & Bradley, M. O. DNA double-strand breaks in mammalian cells after exposure to intercalating agents. Biochimica et Biophysica Acta (BBA)-Nucleic Acids and Protein Synthesis 654, 129–134 (1981).

[2] Ward, J. The yield of DNA double-strand breaks produced intracellularly by ionizing radiation: a review. International journal of radiation biology 57, 1141–1150 (1990).

[3] Arnould, C. & Legube, G. The secret life of chromosome loops upon DNA double-strand break. Journal of molecular biology 432, 724–736 (2020).

[4] Chiarle, R. et al. Genome-wide translocation sequencing reveals mechanisms of chromosome breaks and rearrangements in B cells. Cell 147, 107–119 (2011).

[5] Barlow, J. H. et al. Identification of early replicating fragile sites that contribute to genome instability. Cell 152, 620–632 (2013).

[6] Zhu, Y. et al. qDSB-Seq is a general method for genome-wide quantification of DNA double-strand breaks using sequencing. Nature communications 10, 1–11 (2019).

[7] Vilenchik, M. M. & Knudson, A. G. Endogenous DNA double-strand breaks: production, fidelity of repair, and induction of cancer. Proceedings of the National Academy of Sciences 100, 12871–12876 (2003).

[8] Qiu, Z., Zhang, Z., Roschke, A., Varga, T. & Aplan, P. D. Generation of gross chromosomal rearrangements by a single engineered DNA double strand break. Scientific reports 7, 1–10 (2017).

[9] Taleei, R. & Nikjoo, H. Biochemical DSB-repair model for mammalian cells in G1 and early S phases of the cell cycle. Mutation Research/Genetic Toxicology and Environmental Mutagenesis 756, 206–212 (2013).

[10] Mladenov, E., Magin, S., Soni, A. & Iliakis, G. DNA double-strand-break repair in higher eukaryotes and its role in genomic instability and cancer: Cell cycle and proliferation-dependent regulation. In Seminars in cancer biology, vol. 37, 51–64 (Elsevier, 2016).

[11] Stinson, B. M. & Loparo, J. J. Repair of dna double-strand breaks by the nonhomologous end joining pathway. Annual Review of Biochemistry 90 (2021).

[12] Marcomini, I. et al. Asymmetric processing of DNA ends at a double-strand break leads to unconstrained dynamics and ectopic translocation. Cell reports 24, 2614–2628 (2018).

[13] Shrivastav, M., De Haro, L. P. & Nickoloff, J. A. Regulation of DNA double-strand break repair pathway choice. Cell research 18, 134–147 (2008).

[14] Thompson, L. H. Recognition, signaling, and repair of DNA double-strand breaks produced by ionizing radiation in mammalian cells: the molecular choreography. Mutation Research/Reviews in Mutation Research 751, 158–246 (2012).

[15] Krenning, L., van den Berg, J. & Medema, R. H. Life or death after a break: what determines the choice? Molecular cell 76, 346–358 (2019).

[16] Aleksandrov, R. et al. Protein dynamics in complex dna lesions. Molecular cell 69, 1046–1061 (2018).

[17] Graham, T. G., Walter, J. C. & Loparo, J. J. Two-stage synapsis of DNA ends during non-homologous end joining. Molecular cell 61, 850–858 (2016).

[18] Iarovaia, O. V. et al. Dynamics of double strand breaks and chromosomal translocations. Molecular cancer 13, 1–10 (2014).

[19] Soutoglou, E. et al. Positional stability of single double-strand breaks in mammalian cells. Nature cell biology 9, 675–682 (2007).

[20] Roukos, V. et al. Spatial dynamics of chromosome translocations in living cells. Science 341, 660–664 (2013).

[21] Yu, W. et al. Repair of G1 induced DNA double-strand breaks in S-G2/M by alternative NHEJ. Nature communications 11, 1–15 (2020).

[22] Zhao, B., Rothenberg, E., Ramsden, D. A. & Lieber, M. R. The molecular basis and disease relevance of non-homologous DNA end joining. Nature Reviews Molecular Cell Biology 21, 765–781 (2020).

[23] Zhao, B. et al. The essential elements for the noncovalent association of two DNA ends during NHEJ synapsis. Nature communications 10, 1–12 (2019).

[24] Stinson, B. M., Moreno, A. T., Walter, J. C. & Loparo, J. J. A mechanism to minimize errors during non-homologous end joining. Molecular cell 77, 1080–1091 (2020).

[25] Rothkamm, K., Kruger, I., Thompson, L. H. & Lobrich, M. Pathways of DNA double-strand break repair during the mammalian cell cycle. Molecular and cellular biology 23, 5706–5715 (2003).

[26] Hanscom, T. & McVey, M. Regulation of error-prone dna double-strand break repair and its impact on genome evolution. Cells 9, 1657 (2020).

[27] Alipour, E. & Marko, J. F. Self-organization of domain structures by DNA-loop-extruding enzymes. Nucleic acids research 40, 11202–11212 (2012).

[28] Kim, Y., Shi, Z., Zhang, H., Finkelstein, I. J. & Yu, H. Human cohesin compacts DNA by loop extrusion. Science 366, 1345–1349 (2019).

[29] Golfier, S., Quail, T., Kimura, H. & Brugués, J. Cohesin and condensin extrude DNA loops in a cell cycle-dependent manner. Elife 9, e53885 (2020).

[30] Davidson, I. F. et al. DNA loop extrusion by human cohesin. Science 366, 1338–1345 (2019).

[31] Hansen, A. S. CTCF as a boundary factor for cohesin-mediated loop extrusion: evidence for a multi-step mechanism. Nucleus 11, 132–148 (2020).

[32] Fudenberg, G. et al. Formation of chromosomal domains by loop extrusion. Cell reports 15, 2038–2049 (2016).

[33] Sanborn, A. L. et al. Chromatin extrusion explains key features of loop and domain formation in wild-type and engineered genomes. Proceedings of the National Academy of Sciences 112, E6456–E6465 (2015).

[34] Wang, X., Brandão, H. B., Le, T. B., Laub, M. T. & Rudner, D. Z. Bacillus subtilis SMC complexes juxtapose chromosome arms as they travel from origin to terminus. Science 355, 524–527 (2017).

[35] Davidson, I. F. & Peters, J.-M. Genome folding through loop extrusion by SMC complexes. Nature Reviews Molecular Cell Biology 22, 445–464 (2021).

[36] Ganji, M. et al. Real-time imaging of DNA loop extrusion by condensin. Science 360, 102–105 (2018).

[37] Banigan, E. J. & Mirny, L. A. Loop extrusion: theory meets single-molecule experiments. Current opinion in cell biology 64, 124–138 (2020).

[38] Goloborodko, A., Marko, J. F. & Mirny, L. A. Chromosome compaction by active loop extrusion. Biophysical journal 110, 2162–2168 (2016).

[39] Cattoglio, C. et al. Determining cellular CTCF and cohesin abundances to constrain 3D genome models. Elife 8, e40164 (2019).

[40] Holzmann, J. et al. Absolute quantification of cohesin, CTCF and their regulators in human cells. Elife 8, e46269 (2019).

[41] Taleei, R. & Nikjoo, H. Repair of the double-strand breaks induced by low energy electrons: a modelling approach. International journal of radiation biology 88, 948–953 (2012).

[42] Amitai, A. & Holcman, D. Diffusing polymers in confined microdomains and estimation of chromosomal territory sizes from chromosome capture data. Physical review letters 110, 248105 (2013).

[43] Tiana, G. et al. Structural fluctuations of the chromatin fiber within topologically associating domains. Biophysical journal 110, 1234–1245 (2016).

[44] Miné-Hattab, J. & Rothstein, R. Increased chromosome mobility facilitates homology search during recombination. Nature cell biology 14, 510–517 (2012).

[45] Arnould, C. et al. Loop extrusion as a mechanism for formation of DNA damage repair foci. Nature 590, 660–665 (2021).

[46] Dixon, J. R. et al. Topological domains in mammalian genomes identified by analysis of chromatin interactions. Nature 485, 376–380 (2012).

[47] Lévy-Leduc, C., Delattre, M., Mary-Huard, T. & Robin, S. Two-dimensional segmentation for analyzing Hi-C data. Bioinformatics 30, i386–i392 (2014).

[48] Shin, H. et al. TopDom: an efficient and deterministic method for identifying topological domains in genomes. Nucleic acids research 44, e70–e70 (2016).

[49] Wutz, G. et al. ESCO1 and CTCF enable formation of long chromatin loops by protecting cohesinSTAG1 from WAPL. Elife 9, e52091 (2020).

[50] Li, Y. et al. The structural basis for cohesin–CTCF-anchored loops. Nature 578, 472–476 (2020).

[51] Banigan, E. J. & Mirny, L. A. The interplay between asymmetric and symmetric dna loop extrusion. Elife 9, e63528 (2020).

[52] Cheblal, A. et al. DNA damage-induced nucleosome depletion enhances homology search independently of local break movement. Molecular Cell 80, 311–326 (2020).

[53] Kim, J.-S., Krasieva, T. B., LaMorte, V., Taylor, A. M. R. & Yokomori, K. Specific recruitment of human cohesin to laser-induced DNA damage. Journal of Biological Chemistry 277, 45149–45153 (2002).

[54] Kong, X. et al. Distinct functions of human cohesin-SA1 and cohesin-SA2 in double-strand break repair. Molecular and cellular biology 34, 685–698 (2014).

[55] Liu, Y. et al. Very fast CRISPR on demand. Science 368, 1265–1269 (2020).

[56] Li, K., Bronk, G., Kondev, J. & Haber, J. E. Yeast ATM and ATR kinases use different mechanisms to spread histone H2A phosphorylation around a DNA double-strand break. Proceedings of the National Academy of Sciences 117, 21354–21363 (2020).

[57] Collins, P. L. et al. Dna double-strand breaks induce h2ax phosphorylation domains in a contact-dependent manner. Nature communications 11, 1–9 (2020).

[58] Ochs, F. et al. Stabilization of chromatin topology safeguards genome integrity. Nature 574, 571–574 (2019).

[59] Zhang, Y. et al. The fundamental role of chromatin loop extrusion in physiological V (D) J recombination. Nature 573, 600–604 (2019).

[60] Ba, Z. et al. CTCF orchestrates long-range cohesin-driven V (D) J recombinational scanning. Nature 586, 305–310 (2020).

[61] Dai, H.-Q. et al. Loop extrusion mediates physiological Igh locus contraction for RAG scanning. Nature 590, 338–343 (2021).

[62] Zhang, X. et al. Fundamental roles of chromatin loop extrusion in antibody class switching. Nature 575, 385–389 (2019).

[63] Brandão, H. B. et al. RNA polymerases as moving barriers to condensin loop extrusion. Proceedings of the National Academy of Sciences 116, 20489–20499 (2019).

[64] Pradhan, B. et al. SMC complexes can traverse physical roadblocks bigger than their ring size. bioRxiv (2021).

[65] Brandão, H. B., Ren, Z., Karaboja, X., Mirny, L. A. & Wang, X. DNA-loop-extruding SMC complexes can traverse one another in vivo. Nature Structural & Molecular Biology 1–10 (2021).

[66] Kim, E., Kerssemakers, J., Shaltiel, I. A., Haering, C. H. & Dekker, C. DNA-loop extruding condensin complexes can traverse one another. Nature 579, 438–442 (2020).

[67] Kim, S.-T., Xu, B. & Kastan, M. B. Involvement of the cohesin protein, SMC1, in ATM-dependent and independent responses to DNA damage. Genes & Development 16, 560–570 (2002).

[68] Banigan, E. J., van den Berg, A. A., Brandão, H. B., Marko, J. F. & Mirny, L. A. Chromosome organization by one-sided and two-sided loop extrusion. Elife 9, e53558 (2020).

[69] Ramírez, F. et al. deeptools2: a next generation web server for deep-sequencing data analysis. Nucleic acids research 44, W160–W165 (2016).

[70] Imakaev, M. et al. Iterative correction of Hi-C data reveals hallmarks of chromosome organization. Nature methods 9, 999–1003 (2012).

## References

[1] Zhao, B., Rothenberg, E., Ramsden, D. A. & Lieber, M. R. The molecular basis and disease relevance of non-homologous DNA end joining. Nature Reviews Molecular Cell Biology 21, 765–781 (2020).

[2] Soutoglou, E. et al. Positional stability of single double-strand breaks in mammalian cells. Nature cell biology 9, 675–682 (2007).

[3] Graham, T. G., Walter, J. C. & Loparo, J. J. Two-stage synapsis of DNA ends during non-homologous end joining. Molecular cell 61, 850–858 (2016).

[4] Chen, S. et al. Structural basis of long-range to short-range synaptic transition in NHEJ. Nature 593, 294–298 (2021).

[5] Chaplin, A. K. et al. Cryo-EM of NHEJ supercomplexes provides insights into DNA repair. Molecular cell (2021).

[6] Taleei, R. & Nikjoo, H. Repair of the double-strand breaks induced by low energy electrons: a modelling approach. International journal of radiation biology 88, 948–953 (2012).

[7] Amitai, A. & Holcman, D. Diffusing polymers in confined microdomains and estimation of chromosomal territory sizes from chromosome capture data. Physical review letters 110, 248105 (2013).

[8] Zhu, Y. et al. qDSB-Seq is a general method for genome-wide quantification of DNA double-strand breaks using sequencing. Nature communications 10, 1–11 (2019).

[9] Vilenchik, M. M. & Knudson, A. G. Endogenous DNA double-strand breaks: production, fidelity of repair, and induction of cancer. Proceedings of the National Academy of Sciences 100, 12871–12876 (2003).

[10] Roukos, V. et al. Spatial dynamics of chromosome translocations in living cells. Science 341, 660–664 (2013).

[11] Tiana, G. et al. Structural fluctuations of the chromatin fiber within topologically associating domains. Biophysical journal 110, 1234–1245 (2016).

[12] Shukron, O., Hauer, M. & Holcman, D. Two loci single particle trajectories analysis: constructing a first passage time statistics of local chromatin exploration. Scientific reports 7, 1–11 (2017).

[13] Miné-Hattab, J. & Rothstein, R. Increased chromosome mobility facilitates homology search during recombination. Nature cell biology 14, 510–517 (2012).

[14] Dion, V., Kalck, V., Horigome, C., Towbin, B. D. & Gasser, S. M. Increased mobility of double-strand breaks requires Mec1, Rad9 and the homologous recombination machinery. Nature cell biology 14, 502–509 (2012).

[15] Goloborodko, A., Marko, J. F. & Mirny, L. A. Chromosome compaction by active loop extrusion. Biophysical journal 110, 2162–2168 (2016).

[16] Banigan, E. J. & Mirny, L. A. Limits of chromosome compaction by loop-extruding motors. Physical Review X 9, 031007 (2019).

[17] Cattoglio, C. et al. Determining cellular CTCF and cohesin abundances to constrain 3D genome models. Elife 8, e40164 (2019).

[18] Holzmann, J. et al. Absolute quantification of cohesin, CTCF and their regulators in human cells. Elife 8, e46269 (2019).

[19] Lengronne, A. et al. Cohesin relocation from sites of chromosomal loading to places of convergent transcription. Nature 430, 573–578 (2004).

[20] Newkirk, D. A. et al. The effect of nipped-B-like (Nipbl) haploinsufficiency on genome-wide cohesin binding and target gene expression: modeling Cornelia de Lange syndrome. Clinical epigenetics 9, 1–20 (2017).

[21] Davidson, I. F. & Peters, J.-M. Genome folding through loop extrusion by SMC complexes. Nature Reviews Molecular Cell Biology 22, 445–464 (2021).

[22] Busslinger, G. A. et al. Cohesin is positioned in mammalian genomes by transcription, CTCF and Wapl. Nature 544, 503–507 (2017).

[23] Brandão, H. B. et al. RNA polymerases as moving barriers to condensin loop extrusion. Proceedings of the National Academy of Sciences 116, 20489–20499 (2019).

[24] Canela, A. et al. Genome organization drives chromosome fragility. Cell 170, 507–521 (2017).

[25] Shastri, N. et al. Genome-wide identification of structure-forming repeats as principal sites of fork collapse upon ATR inhibition. Molecular cell 72, 222–238 (2018).

[26] Canela, A. et al. Topoisomerase II-induced chromosome breakage and translocation is determined by chromosome architecture and transcriptional activity. Molecular cell 75, 252–266 (2019).

[27] Gothe, H. J. et al. Spatial chromosome folding and active transcription drive DNA fragility and formation of oncogenic MLL translocations. Molecular cell 75, 267–283 (2019).

[28] Madabhushi, R. et al. Activity-induced DNA breaks govern the expression of neuronal early-response genes. Cell 161, 1592–1605 (2015).

[29] Puerto, S. et al. Induction, processing and persistence of radiation-induced chromosomal aberrations involving hamster euchromatin and heterochromatin. Mutation Research/Genetic Toxicology and Environmental Mutagenesis 469, 169–179 (2000).

[30] Hansen, A. S. CTCF as a boundary factor for cohesin-mediated loop extrusion: evidence for a multi-step mechanism. Nucleus 11, 132–148 (2020).

[31] Ganji, M. et al. Real-time imaging of DNA loop extrusion by condensin. Science 360, 102–105 (2018).

[32] Kong, M. et al. Human condensin i and ii drive extensive ATP-dependent compaction of nucleosome-bound DNA. Molecular cell 79, 99–114 (2020).

[33] Golfier, S., Quail, T., Kimura, H. & Brugués, J. Cohesin and condensin extrude DNA loops in a cell cycle-dependent manner. Elife 9, e53885 (2020).

[34] Kim, Y., Shi, Z., Zhang, H., Finkelstein, I. J. & Yu, H. Human cohesin compacts DNA by loop extrusion. Science 366, 1345–1349 (2019).

[35] Davidson, I. F. et al. DNA loop extrusion by human cohesin. Science 366, 1338–1345 (2019).

[36] Rao, S. S. et al. Cohesin loss eliminates all loop domains. Cell 171, 305–320 (2017).

[37] Gibcus, J. H. et al. A pathway for mitotic chromosome formation. Science 359 (2018).

[38] Hansen, A. S., Pustova, I., Cattoglio, C., Tjian, R. & Darzacq, X. CTCF and cohesin regulate chromatin loop stability with distinct dynamics. Elife 6, e25776 (2017).

[39] Walther, N. et al. A quantitative map of human Condensins provides new insights into mitotic chromosome architecture. Journal of Cell Biology 217, 2309–2328 (2018).

[40] Li, Y. et al. The structural basis for cohesin–CTCF-anchored loops. Nature 578, 472–476 (2020).

[41] Wutz, G. et al. ESCO1 and CTCF enable formation of long chromatin loops by protecting cohesinSTAG1 from WAPL. Elife 9, e52091 (2020).

[42] Arruda, N. L. et al. Distinct and overlapping roles of STAG1 and STAG2 in cohesin localization and gene expression in embryonic stem cells. Epigenetics & chromatin 13, 1–17 (2020).

[43] Gerlich, D., Koch, B., Dupeux, F., Peters, J.-M. & Ellenberg, J. Live-cell imaging reveals a stable cohesin-chromatin interaction after but not before DNA replication. Current Biology 16, 1571–1578 (2006).

[44] Cuadrado, A. & Losada, A. Specialized functions of cohesins STAG1 and STAG2 in 3D genome architecture. Current opinion in genetics & development 61, 9–16 (2020).

[45] Remeseiro, S., Cuadrado, A., Gómez-López, G., Pisano, D. G. & Losada, A. A unique role of cohesin-SA1 in gene regulation and development. The EMBO journal 31, 2090–2102 (2012).

[46] Kojic, A. et al. Distinct roles of cohesin-SA1 and cohesin-SA2 in 3D chromosome organization. Nature structural & molecular biology 25, 496–504 (2018).

[47] Arnould, C. et al. Loop extrusion as a mechanism for formation of DNA damage repair foci. Nature 590, 660–665 (2021).

[48] Caron, P. et al. Cohesin protects genes against γH2AX induced by DNA double-strand breaks. PLoS genetics 8, e1002460 (2012).

[49] Meisenberg, C. et al. Repression of transcription at DNA breaks requires cohesin throughout interphase and prevents genome instability. Molecular cell 73, 212–223 (2019).

[50] Cheblal, A. et al. DNA damage-induced nucleosome depletion enhances homology search independently of local break movement. Molecular Cell 80, 311–326 (2020).

